# Synergistic, Multi-level Understanding of Psychedelics: Three Systematic Reviews and Meta-analyses of Their Pharmacology, Neuroimaging and Phenomenology

**DOI:** 10.1101/2023.10.06.561183

**Authors:** Kenneth Shinozuka, Katarina Jerotic, Pedro Mediano, Alex T. Zhao, Katrin H. Preller, Robin Carhart-Harris, Morten L. Kringelbach

**Affiliations:** Centre for Eudaimonia and Human Flourishing, Linacre College, University of Oxford, Oxford, OX3 9BX, UK; Department of Psychiatry, University of Oxford, Oxford, OX3 7JX, UK; Oxford Mathematics of Consciousness and Applications Network (OMCAN), University of Oxford, Oxford, OX2 6GG, UK; Department of Computing, Imperial College London, London, SW7 2RH, UK; Department of Statistics and Data Science (Alumnus), The Wharton School, University of Pennsylvania, Philadelphia, PA 19104, USA; Departments of Psychiatry, Neuroscience, and Psychology, Yale University, New Haven, CT 06511, USA; Department of Psychiatry, Psychotherapy and Psychosomatics, Psychiatric University Hospital Zurich, University of Zurich, Zurich, 8032, Switzerland; Centre for Psychedelic Research, Imperial College London, London, SW7 2AZ, UK; Department of Neurology, University of California, San Francisco, San Francisco, CA 94158, USA; Department of Neurology, Psychiatry and Behavioral Sciences, University of California, San Francisco, CA 94143, USA; Center for Music in the Brain, Department of Clinical Medicine, Aarhus University, Aarhus, 8000, Denmark

**Keywords:** psychedelics, DMT, ayahuasca, LSD, psilocybin, meta-analysis, fMRI, functional connectivity, pharmacology, receptor agonism, GPCR, phenomenology, subjective experience, consciousness

## Abstract

Serotonergic psychedelics induce altered states of consciousness and have shown potential for treating a variety of neuropsychiatric disorders, including depression and addiction. Yet their modes of action are not fully understood. Here, we provide a novel, synergistic understanding of psychedelics arising from systematic reviews and meta-analyses of three hierarchical levels of analysis: 1) subjective experience (phenomenology), 2) neuroimaging and 3) molecular pharmacology. Phenomenologically, medium and high doses of LSD yield significantly higher ratings of visionary restructuralisation than psilocybin on the 5-dimensional Altered States of Consciousness Scale. Our neuroimaging results reveal that, in general, psychedelics significantly strengthen between-network functional connectivity (FC) while significantly diminishing within-network FC. Pharmacologically, LSD induces significantly more inositol phosphate formation at the 5-HT_2A_ receptor than DMT and psilocin, yet there are no significant between-drug differences in the selectivity of psychedelics for the 5-HT_2A_, 5-HT_2C_, or D_2_ receptors, relative to the 5-HT_1A_ receptor. Our meta-analyses link DMT, LSD, and psilocybin to specific neural fingerprints at each level of analysis. The results show a highly non-linear relationship between these fingerprints. Overall, our analysis highlighted the high heterogeneity and risk of bias in the literature. This suggests an urgent need for standardising experimental procedures and analysis techniques, as well as for more research on the emergence between different levels of psychedelic effects.

## Section 1. Introduction

Psychedelics, derived from the Greek words “mind” and “manifesting,” are hallucinogenic drugs that profoundly alter consciousness. The “classic” psychedelics are serotonergic substances that include lysergic acid diethylamide (LSD), psilocybin (the primary psychoactive ingredient in magic mushrooms), and N,N-dimethyltryptamine (DMT, the main psychoactive chemical in ayahuasca). Recently, the term “psychedelic” has been applied to other mind-expanding yet non-serotonergic drugs, including ketamine and 3,4-methylenedioxymethampetamine (MDMA, also known as ecstasy). For the sake of this paper, we will only consider the classic psychedelics.

Psychedelics have been utilised by early cultures for millennia within diverse sociocultural contexts, as well as spiritual and healing rituals (Garcia & Maia, 2022; Nichols, 2016; Tupper et al., 2015). Recently, researchers have come to recognise that psychedelics may be effective tools in the treatment of psychiatric disorders such as depression (Muttoni et al., 2019; Nutt et al., 2020) and addiction (Tófoli & de Araujo, 2016). It is worth noting that, in the contemporary era of psychedelic research, psychedelics are typically only administered as adjuncts to therapy (hence the phrase “psychedelic-assisted therapy”), which may confound the therapeutic effects of the drugs themselves (Goodwin et al., 2024).

The effects of psychedelic substances have been examined at several levels. Pharmacological research has measured their interaction with various receptors in the brain (e.g. Glennon et al., 1984; Watts et al., 1995), while other research domains have explored the subjective experience, or phenomenology, of a psychedelic ‘trip’. Neuroimaging research has examined changes in brain activity and functional connectivity under the influence of psychedelics, yet there are still relatively few studies and a plethora of different analysis techniques. Here, we report the findings of three systematic meta-analyses focused on synthesising the evidence about three classical psychedelics: DMT, LSD, and psilocybin. The main aim is to analyse the literature across three levels of description: (1) phenomenology—to discuss key differences in the states of consciousness elicited by psychedelics; (2) functional neuroimaging—to compare the changes in brain activity, functional connectivity, and entropy that are induced by psychedelics; and (3) pharmacology—to discuss the binding affinities and functional activity of psychedelics with respect to various serotonergic and dopaminergic receptors.

### Section 1.1. Phenomenology

Psychedelics produce altered states of consciousness that are characterised by visual hallucinations, ego death, or a breakdown in one’s sense of self, and spiritual or “mystical” experiences (Griffiths et al., 2018; Yaden et al., 2021). The Altered States of Consciousness (ASC) scale is one popular questionnaire that captures several of these subjective effects. It measures these states along five different dimensions: oceanic boundlessness (e.g. a sense of interconnectedness), anxious ego dissolution, visionary restructuralisation (e.g. visual hallucinations), auditory alterations, and reduction of vigilance. Subsequent factor analyses led to the extraction of 11 lower-order dimensions from the 5D-ASC, resulting in the development of the 11D-ASC: experience of unity, spiritual experience, blissful state, insightfulness, disembodiment, impaired control and cognition, anxiety, elementary imagery, complex imagery, audio-visual synesthesia, and changed meaning of percepts (Studerus et al., 2010). Because the ASC is the most common subjective questionnaire in the literature on the neuroimaging of psychedelics, we performed a meta-analysis on both the 5D- and 11D-ASC scores of DMT, LSD, and psilocybin.

However, the ASC is just one of many scales for rating the subjective effects of psychedelics. The multiplicity of scales makes it difficult to comprehensively evaluate the literature on the phenomenology of psychedelics. To date, only two studies have used a single set of scales to compare the subjective effects of LSD and psilocybin in the same group of participants (Holze et al., 2022; Ley et al., 2023). More recent research has attempted to draw direct comparisons by qualitatively examining larger datasets, such as the Erowid database of trip reports (Ballentine et al., 2022; Coyle et al., 2012; Qiu & Minda, 2021; Sanz et al., 2018; Zamberlan et al., 2018). The key findings from these comparative studies are discussed in *Section S3*.

### Section 1.2. Neuroimaging

The first functional magnetic resonance imaging (fMRI) study of psychedelics, published in 2012 (Carhart-Harris, Erritzoe, et al., 2012), was an exploratory analysis of psilocybin-induced changes in cerebral blood flow and BOLD activity in healthy human participants. Since then, dozens of fMRI studies have been performed on human participants under the influence of ayahausca, LSD, psilocybin and DMT, including several studies on depressed patients (Carhart-Harris et al., 2017; Daws et al., 2022; Doss et al., 2021; Mertens et al., 2020; Roseman et al., 2018; Wall et al., 2022).

The fMRI studies assess three different metrics of brain activity: BOLD activation, connectivity, and entropy. Studies on BOLD activation analyse changes in the trajectory of BOLD timeseries with a rapidly-acting psychedelic, such as intravenously-administered psilocybin. Connectivity studies can examine undirected, instantaneous correlations between different regions (a type of functional connectivity) or model the experimental factors that modulate directed connections between regions, using dynamic causal modeling (a tool for measuring effective connectivity). The entropy of spontaneous brain activity is best assessed with MEG and EEG, but efforts to compute entropy on the spontaneous BOLD signal, as well as the entropy rate (Lempel-Ziv complexity) and indices of criticality, have also been attempted (see *Section S1.2* for a full list of citations). While studies on BOLD activation and functional connectivity are common for most subject areas within cognitive neuroscience, entropy is a rather unique feature of the psychedelic neuroimaging literature. The studies on entropy were motivated by Robin Carhart-Harris’ influential Entropic Brain Hypothesis (EBH) (Carhart-Harris et al., 2014; Carhart-Harris, 2018), which states that psychedelics alter consciousness by elevating the entropy of spontaneous brain activity across time.

We performed a quantitative meta-analysis of the pairwise functional connectivity data using a novel algorithm. We initially attempted to conduct a quantitative meta-analysis of the BOLD activation data with the GingerALE method, which examines common clusters of BOLD activity across studies. However, we found that the data was too heterogeneous for the results to be valid. From the outset, we decided to provide a qualitative discussion of the studies on entropy, due to the wide range of methods in that section of the literature.

We chose to perform a meta-analysis exclusively on the fMRI data and not on any of the other modalities used to measure brain activity, such as PET, SPECT, MEG, and EEG. There were too few primary PET (4), SPECT (4), or MEG (2) datasets to merit a meta-analysis; by comparison, 30 fMRI studies have collected original data. While there are many more primary EEG datasets (25) from the post-1960s era (and even more from the 1950s-1960s era) than PET, SPECT, or MEG, most of the EEG studies do not spatially localise the brain activity recorded at EEG electrodes. Without the raw data, which may be difficult to access when some of the studies are nearly 20 years old, we cannot source-localise the EEG activity ourselves. On the other hand, most fMRI studies report the spatial coordinates of the brain regions that become more or less active or connected on psychedelics. In general, EEG has higher temporal resolution than fMRI (Haufe et al., 2018), so a future meta-analysis that focuses on the temporal, rather than spatial, characteristics of brain activity on psychedelics could consider the EEG rather than fMRI data.

### Section 1.3. Pharmacology

All classic psychedelics activate serotonin receptors. In particular, their interaction with the serotonin-2A (5-HT_2A)_ receptor is their primary mechanism of action (González-Maeso et al., 2007; López-Giménez & González-Maeso, 2018). Multiple studies have demonstrated that blocking the 5-HT_2A_ receptor with antagonists such as ketanserin eliminates their subjective effects (Preller et al., 2017; Quednow et al., 2012; Vollenweider et al., 1998). 5-HT_2A_ expression is profuse in human cortex, with one study finding highest expression in the posterior cingulate cortex and V1 (Beliveau et al., 2017). In both humans and monkeys, 86-100% of glutamatergic cells in layers II-V of cortex express mRNA encoding 5-HT_2A_ receptors (de Almeida & Mengod, 2007). 5-HT_2A_ has also been identified in many subcortical regions (Nichols, 2016), though human PET imaging implies lower expression than in the cortex (Beliveau et al., 2017).

Although the consciousness-altering effects of psychedelics do primarily arise from their action at the 5-HT_2A_ receptor, different psychedelics display different binding profiles. For instance, unlike DMT and psilocybin, LSD has moderate affinity for dopamine receptors (de Vos et al., 2021). Differences in binding affinities could partially account for the distinctive phenomenology of each psychedelic. There is also some evidence that the 5-HT_1A_ and 5-HT_2C_ receptors may mediate the effects of psychedelics (Canal & Murnane, 2017; Pokorny et al., 2016).

However, binding affinity paints an incomplete picture of the pharmacology of psychedelics. Another key aspect is their functional activity at receptors. Serotonin and dopamine receptors tend to be G protein-coupled receptors (GPCRs) (Gurevich et al., 2016; Shah et al., 2020), which perform essential roles in neurophysiological processes like smell, taste, light perception, and more (de Oliveira et al., 2019). GPCRs regulate these processes through G protein signaling pathways, which are initiated when agonists bind to GPCRs and activate G proteins (Tuteja, 2009). Different families of G proteins are selective for particular signaling pathways (Giulietti et al., 2014). In a phenomenon known as biased agonism, agonists can induce certain conformations in GPCRs, which then selectively activate certain pathways and not others (Andresen, 2011). 5-HT_2_ receptors couple preferentially to a family of G proteins known as Gα_q/11_, which activates the enzyme phospholipase C (PLC). This enzyme then catalyses the synthesis of the secondary messenger inositol triphosphate (IP_3_, a species of inositol phosphate [IP]) , which subsequently leads to the release or “mobilisation” of calcium from the endoplasmic reticulum (López-Giménez & González-Maeso, 2018). Psychedelics are also known to recruit β-arrestin proteins (Pottie, Cannaert, et al., 2020; Pottie, Dedecker, et al., 2020; Rodriguiz et al., 2021; Schmitz et al., 2022). These proteins block the interaction between G proteins and the GPCR and remove the GPCR from the cell membrane, while also coupling GPCRs to signaling proteins without activating G proteins; thus, β-arrestin promotes alternative signaling pathways (Bohn & Schmid, 2010). Functional GPCR assays on psychedelics have tended to focus on the three aspects of GPCR signaling described above: (1) IP formation, (2) calcium mobilisation, and (3) β-arrestin (specifically β-arrestin2) recruitment.

We performed a meta-analysis of the selective affinity of DMT, LSD, and psilocin for the 5-HT_2A_, 5-HT_2C_, and D_2_ receptors, relative to the 5-HT_1A_ receptor. Additionally, we conducted a meta-analysis of their functional activity at the 5-HT_2A_ receptor, as measured by the three aforementioned assays.

## Section 2. Results

We present the results of our meta-analysis of the phenomenology, neuroimaging, and pharmacology of three classical psychedelics: DMT, LSD, and psilocybin. At each level, we measured the alignment between the corresponding results and the Yeo networks, which facilitated comparisons between the three hierarchical levels.

### Section 2.1. Phenomenology

We performed a meta-analysis on the 5D- and 11D-ASC scores of DMT, LSD, and psilocybin. Our literature search identified *n* = 44 phenomenology studies that met the inclusion criteria, including 5 studies on DMT (5D-ASC: *n* = 3; 11D-ASC: *n* = 3), 14 studies on LSD (5D-ASC: *n* = 9; 11D-ASC: *n* = 12), and 25 studies on psilocybin (5D-ASC: *n* = 12; 11D-ASC: *n* = 17). The eleven dimensions of the 11D-ASC are subscales of three dimensions in the 5D-ASC: oceanic boundlessness (OB), anxious ego dissolution (AED), and visionary restructuralisation (VR). Nevertheless, the 11D-ASC questionnaire contains fewer items than the 5D-ASC questionnaire, so 5D-ASC data cannot be directly compared to 11D-ASC data. Hence, we ran separate meta-analyses for the 5D-ASC and 11D-ASC data. In order to account for the measurement of multiple subjective dimensions, sometimes with more than one dose or drug, in individual studies, we performed a multilevel random-effects meta-analysis (Assink & Wibbelink, 2016; Harrer et al., 2021).

DMT was administered intravenously (IV) in the phenomenological studies, whereas LSD and psilocybin were administered orally (we excluded phenomenological studies that used IV LSD and psilocybin). IV administration yields different pharmacokinetics, which are known to influence the subjective experience of the drug (Luan et al., 2023; Vogt et al., 2023). Therefore, while we conducted meta-analyses of all three drugs, we only examined significant between-drug differences for ASC ratings of LSD and psilocybin. Additionally, we only compared similar doses of LSD and psilocybin. We assumed 20 mg psilocybin is equivalent to 0.01 mg LSD (Ley et al., 2023), and we defined low, medium, and high doses based on the literature (Carbonaro et al., 2018; Hasler et al., 2004; Vollenweider et al., 2007).

Pooled 5D-ASC scores for DMT, LSD, and psilocybin are shown in **Figure 1** (numerical data are given in **Table S1**). Between-drug differences for LSD and psilocybin are displayed in the left column of **Figure 1a-c**, and within-drug differences are displayed in the right column. At medium doses (LSD: 0.075-0.109 mg, psilocybin: 15-21 mg), 5D-ASC scores were significantly greater for LSD than psilocybin in the OB dimension (*p* = 0.0283), which corresponds to feelings of interconnectedness, and the VR dimension (*p* = 0.0468), which measures the quality and intensity of visual hallucinations. We also observed a significant difference between LSD and psilocybin in the VR dimension (*p* = 0.0417) at high doses (LSD: ≥0.010 mg, psilocybin: ≥22 mg). There were no significant differences at low doses (LSD: 0.050-0.074 mg, psilocybin: 8-14 mg).

**Figure 1.**
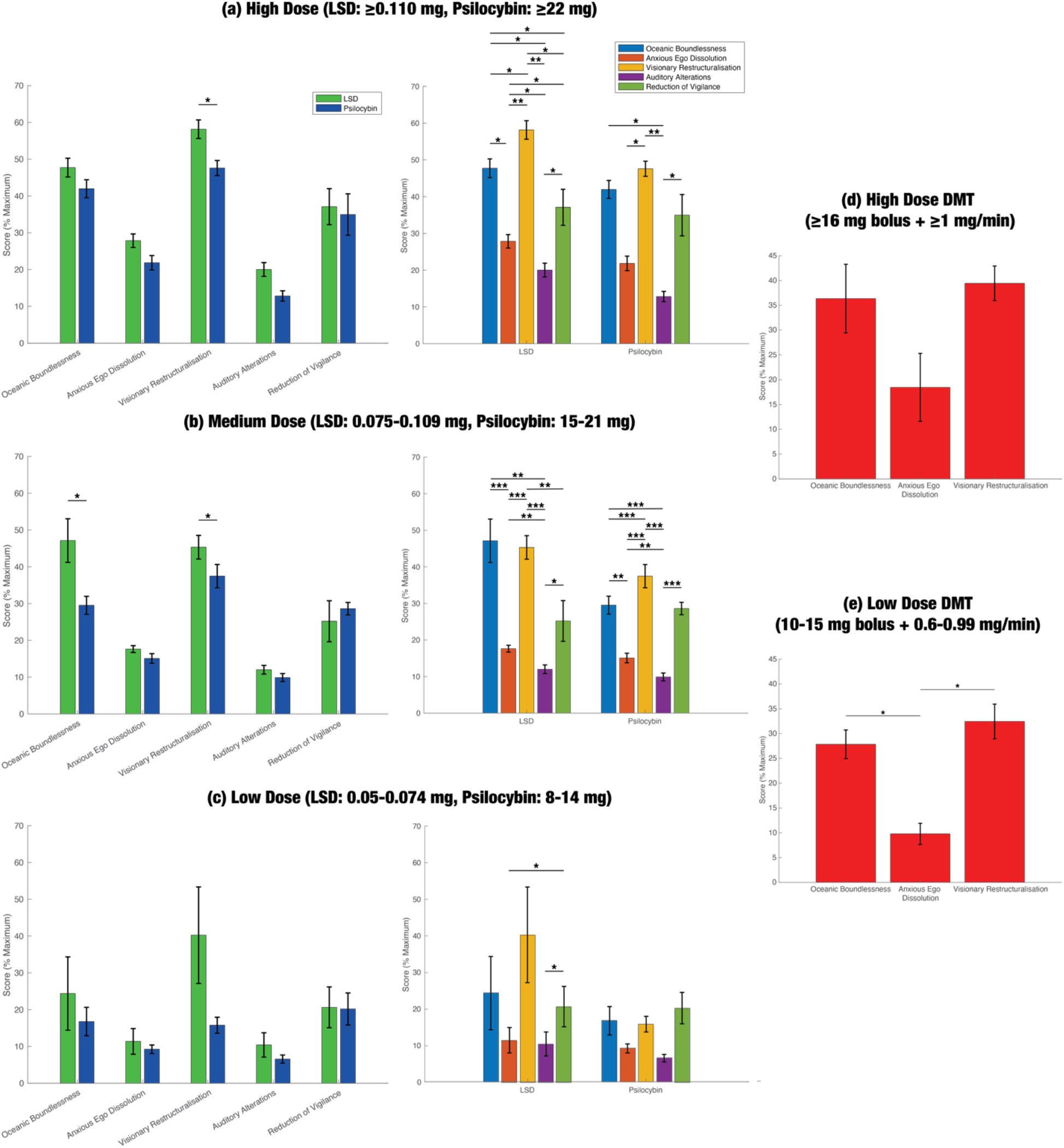
Meta-analysis of the 5-Dimensional Altered States of Consciousness (5D-ASC) data reveals few significant differences between psychedelics, but many more significant differences within psychedelics. The 5D-ASC is one of the most common scales for assessing the subjective effects of psychedelics. It measures altered states of consciousness along five different dimensions: oceanic boundlessness (OB; a feeling of interconnectedness), anxious ego dissolution (AED), visionary restructuralisation (VR; the quality and intensity of visual hallucinations), auditory alterations (AA), and reduction of vigilance (RV). Our literature search identified 23 studies that reported 5D-ASC data. We performed a multilevel random-effects meta-analysis in order to account for the lack of statistical independence between measurements of different dimensions within the same group of participants. Within-study and between-study heterogeneity were estimated with the restricted maximum likelihood procedure. Using a Correlated and Hierarchical Effects model to account for within-study correlations in sampling error, we analysed the effect of dose and drug (DMT, LSD, or psilocybin) on pooled 5D-ASC scores. DMT was administered intravenously in phenomenological studies, whereas LSD and psilocybin were administered orally. Thus, DMT has different pharmacokinetics from LSD and psilocybin, which impacts the subjective experience, so we only compared significant between-drug differences for LSD and psilocybin. We only compared the two drugs for similar doses, assuming that 0.1 mg LSD = 20 mg psilocybin (Ley *et al*., 2023). (a-c, left column) Between-drug comparisons of pooled 5D-ASC scores for LSD and psilocybin. LSD almost always ranked higher than psilocybin, but differences only reached significance in the VR dimension for high and medium doses and in the OB dimension for medium dose. (a-c, right column) Within-drug comparisons for LSD and psilocybin. VR and OB received significantly higher scores than AED and AA, for both LSD and psilocybin. (d-e) Within-drug comparisons for DMT. At low doses, OB and VR ranked significantly higher than AED. Only one study measured ASC scores for the AA and RV dimensions on DMT, so we did not perform a meta-analysis on these dimensions for DMT. Numerical data for the above figure are given in Table S1. We also conducted a meta-analysis of the 11-dimensional ASC scale, which is also widely used in the literature; the results are displayed in Figure S3 and Table S2. * < 0.05, ** < 0.01, *** < 0.001.

Within-drug differences between the five subjective dimensions were quite similar. A clear separation can be seen between two groups of subjective categories: (1) OB, VR, RV, and (2) AA and AED. At medium doses, VR and OB scores were significantly higher than both AED and AA scores for both LSD and psilocybin. For DMT, there were no significant differences between dimensions at high doses (≥16 mg bolus injection, followed by continuous infusion of 1 mg/min), but for low doses (10-15 mg bolus + 0.6-0.99 mg/min continuous infusion), AED ranked significantly lower than both OB and VR. (Note that only one study measured AA and RV scores for DMT (Vogt et al., 2023), so we did not perform a meta-analysis on these dimensions for DMT.)

The results of the 11D-ASC meta-analysis were not very consistent with those of the 5D-ASC meta-analysis (**Figure S3**; **Table S2**). Note that we could not include DMT in the 11D-ASC meta-analysis because two of the three studies on DMT did not report standard errors, which are necessary for pooling the subjective data. At high doses, there were no significant differences between LSD and psilocybin in any of the dimensions, and standard errors were very large for psilocybin in some dimensions, particularly in some OB and VR subscales. At medium doses, where standard errors tended to be lower, we observed that two of the OB subscales (experience of unity and insightfulness), one of the AED subscales (impaired control and cognition), and three of the VR subscales (complex imagery, audio-visual synaesthesia, and changed meaning of percepts) ranked significantly higher for LSD than psilocybin. In line with the 5D-ASC results, we did not find any significant differences between LSD and psilocybin at low doses. However, unlike the 5D-ASC analysis, the 11D-ASC analysis showed that psilocybin rated higher than LSD in most of the dimensions; that being said, standard errors were very high for both LSD and psilocybin, so our estimates are not reliable. For medium doses, within-drug comparisons for the 11D-ASC analysis generally aligned with those of the 5D-ASC analysis.

We did not find any significant relationship between the pooled 5D-ASC scores and three methodological covariates that were defined *a priori*: prior psychedelic use, the time that the questionnaire was administered relative to the drug, and the presence or absence of a task in the experiment. Additionally, there was significant residual heterogeneity in the pooled 5D-and 11D-ASC scores (*p* < 0.0001 in both cases). Unfortunately, we were unable to obtain reliable estimates of heterogeneity specifically attributable to within-study and between-study differences. For three of the 5D-ASC scales, we found evidence of significant publication bias with Egger’s regression test, which determines whether small studies report disproportionately high effect sizes (OB: *p* = 0.0187, AED: *p* < 0.0001, VR: *p* = 0.6279, AA: *p* < 0.0001, RV: *p* = 0.6532). The results of our risk-of-bias assessments for individual studies are shown in **Table S3**. Due to the moderate-to-serious risk of bias, significant residual heterogeneity, high probability of publication bias, and indirectness (some studies measured subjective effects while participants were performing a specific task, as opposed to during resting-state), our certainty in the body of phenomenological evidence is low.

In order to directly compare the phenomenology of psychedelics to their neuroimaging and pharmacological profiles (*Section 2.2* and *Section 2.3*), we sought to determine the “phenomenology profiles” of DMT, LSD, and psilocybin based on the neural correlates of their subjective effects. We performed a separate literature search of studies that measured correlations between ASC ratings and fMRI activity. We determined the Yeo network that contained each brain region that exhibited a significant correlation. The phenomenology profiles were defined by a weighted combination of the mean correlations between each ASC scale and each Yeo network, in which the weights were the pooled ASC ratings of high doses of each psychedelic for the corresponding scale.

Because there were few significant differences in the pooled ASC scores, the phenomenology profiles of the psychedelics look very similar (**Figure 2**). However, these profiles may not reflect the true neural correlates of the subjective effects of psychedelics. Many studies performed correlations between ASC ratings and brain regions that were active only in specific tasks, but the tasks varied strongly in the literature. There need to be more studies that link whole-brain fMRI activity to ASC ratings before we can estimate the phenomenology profiles with confidence.

**Figure 2.**
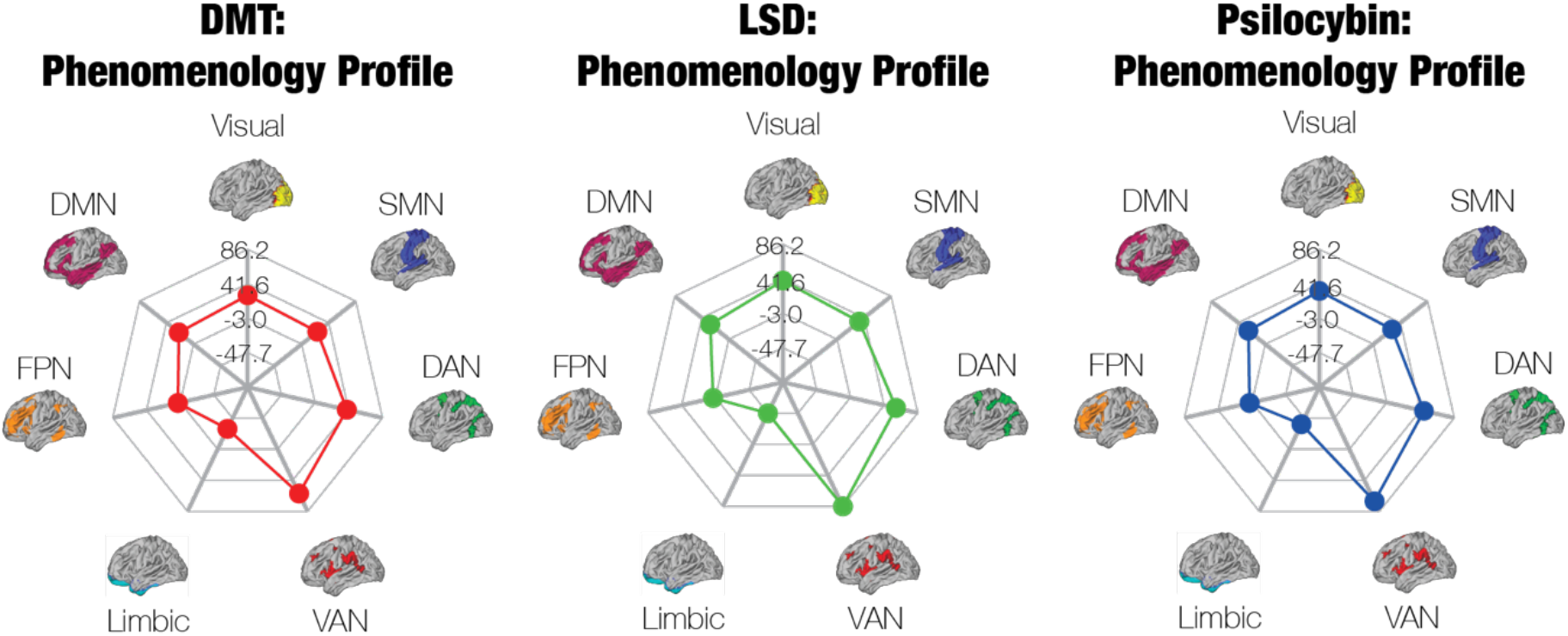
Phenomenology profiles of the psychedelics demonstrate broad similarity between the neural correlates of their subjective effects. We identified 14 studies that measured correlations between ASC ratings and fMRI activity or pairwise connectivity, including three studies on ketamine. We determined the Yeo network that contained each brain region that exhibited a significant correlation, and then we averaged across the correlations associated with each Yeo network, resulting in the mean correlation between each Yeo network and each ASC scale. The phenomenology profiles were defined by a weighted combination of the mean correlations for each ASC scale, in which the weights were the pooled ASC ratings of high doses of each psychedelic for the corresponding scale. Because the pooled ASC ratings are similar across psychedelics, the phenomenology profiles are very alike one another as well. However, the lack of data on the neural correlates of the ASC ratings limits our confidence in the validity of these profiles.

Finally, we also aimed to relate the phenomenology of different psychedelics by qualitatively synthesising the findings of papers that directly compared the subjective experiences of DMT/ayahuasca, LSD, and psilocybin. *Section S3* contains a comprehensive summary of their major aims, analysis approaches and findings.

### Section 2.2. Neuroimaging

As stated above, the fMRI studies on psychedelics measure three different characteristics of brain activity: BOLD activation, entropy, and connectivity. We identified *n* = 17 studies on BOLD activation, including *n* = 3 studies on ayahuasca or DMT, *n* = 7 studies on LSD, and *n* = 4 studies on psilocybin. Initially, we used the GingerALE method to determine common clusters of BOLD activation across studies. We observed that psilocybin was associated with a cluster of activity in visual cortices, whereas LSD tended to affect more frontal areas. However, three of the four studies on psilocybin displayed visual stimuli to participants. Therefore, our results were likely confounded by the tasks that were used in the literature. We chose not to report the results of our GingerALE analysis because they are likely misleading.

The *n* = 12 fMRI studies on entropy and criticality in the psychedelic literature are very heterogeneous, employing a wide variety of measures on dissimilar variables of interest. A quantitative meta-analysis was extremely challenging, so we opted instead to perform a qualitative review of this literature, which is reported in *Section S4*.

The studies on connectivity were the only segment of the fMRI literature for which we determined that we could conduct a valid meta-analysis. These studies tended to use the same resting-state experimental procedure, and many of them employed similar methods for measuring connectivity. However, one major obstacle was the variety of parcellations in the literature. Since each study defined regions of interest (ROIs) in a different way, it was not possible to simply perform a weighted average of connectivity values across studies. (This weighted average method is essentially the core technique of the phenomenology and pharmacology meta-analyses.) An alternative approach is to “re-reference” the data to a coarse-grained parcellation: the Yeo networks, of which there are only seven in the brain <u>(Thomas Yeo et al., 2011</u>). That is, for every pair of functionally connected ROIs in the literature, we determine the corresponding Yeo networks that contain them. If the connection is positive (it increases under psychedelics), then the aggregate FC between those Yeo networks is incremented; otherwise, it is decremented. Our method for weighting each connection in the literature is described in *Section S2.2.2.* We determined significance by repeatedly applying our method to an independent resting-state fMRI dataset of sober, healthy individuals (Van Essen et al., 2013), which resulted in a null distribution (**Figure S4**).

This method is capable of synthesising studies on *pairwise* FC, such as seed-to-seed, seed-to-voxel, and within- and between-network FC analyses using independent components analysis (see **Table S5** for a full description of the inputs). The algorithm cannot accommodate global, structural metrics of connectivity such as global brain connectivity (GBC) or graph-theoretic modularity. Additionally, the algorithm does not capture any information about the direction of connectivity between regions; therefore, we did not incorporate studies of effective connectivity or directed FC. (However, the vast majority of FC studies measured undirected FC.)

Many FC studies are secondary analyses of the same primary dataset. For instance, the primary FC data from Carhart-Harris et al. (2016) was re-analysed at least seven times with different methods in subsequent studies. To mitigate bias, we only inputted the most informative analysis of each unique dataset into the meta-analysis algorithm, such that each dataset was only represented once in the meta-analysis. Our method for selecting the most informative analysis is described in *Section S2.2.2*.

Based on these criteria and others, we ultimately identified *n* = 12 studies on connectivity that were eligible for our meta-analysis, including *n* = 3 studies on ayahuasca or DMT, *n* = 3 studies on LSD, and *n* = 6 studies on psilocybin. (There are 22 other studies on connectivity that we qualitatively review in *Section S5*.) Our estimates of aggregate FC between Yeo networks, across all psychedelics, are shown in **Figure 3**. Psychedelics significantly strengthened between-network connectivity across most pairs of networks. Within-network connectivity significantly decreased in the visual network, ventral attention network (VAN), and default mode network (DMN), yet significantly increased in the dorsal attention network (DAN) and frontoparietal network (FPN). Connectivity between the limbic network and other networks was generally deemed insignificant, likely due to the small size of this network in the Yeo parcellation and therefore low number of seed regions in the literature that predominantly overlap with it.

**Figure 3.**
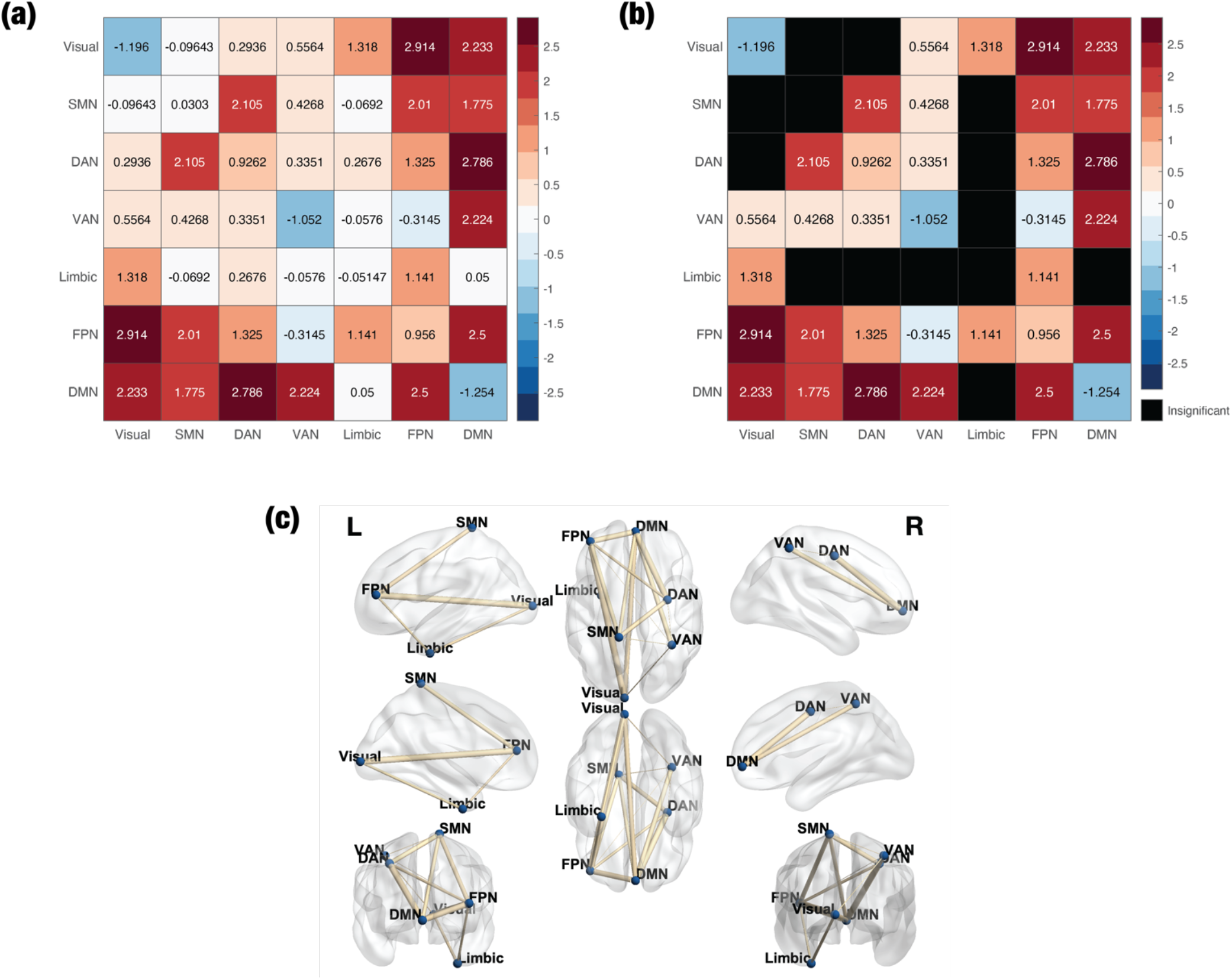
Meta-analysis of the functional connectivity (FC) data indicates that psychedelics potently increase between-network FC. To perform a meta-analysis on the FC data, we determined the Yeo networks that contained each pair of functionally connected regions of interest (ROIs) in the literature and then performed a weighted sum of the number of significant connections between Yeo networks. We included n = 12 studies in this meta-analysis, after excluding studies that did not measure pairwise FC and secondary analyses of identical datasets. (a) The aggregate FC matrix shows the overall connectivity between pairs of Yeo networks. (b) Several connections were deemed to be insignificant relative to a null distribution that was formed from an independent resting-state fMRI dataset collected by the Human Connectome Project. (c) A rendering of (b) on the surface of the brain, created with the BrainNet Viewer (Xia *et al*., 2013).

In addition to aggregating FC across all psychedelics, we performed our analysis separately on each individual psychedelic (**Figure S5**). All three of the psychedelics significantly reduced the within-network FC of the visual network while generally elevating between-network FC. Intriguingly, LSD significantly increased FC within the DMN, but this is largely due to a high number of significant connections that were identified in a single study (Bedford et al., 2023) between regions that are typically not considered to be core elements of the DMN yet are nevertheless classified by the Yeo parcellation as regions within the DMN, such as the frontal pole and inferior frontal gyrus (Liu et al., 2023; Martín-Signes et al., 2021; Rai et al., 2021). LSD significantly elevated FC between the limbic network and visual network, as well as between the limbic network and FPN, whereas FC between the limbic network and all other networks was insignificant for ayahuasca/DMT and psilocybin. Ayahuasca/DMT was the only psychedelic to significantly reduce FC between the visual network and SMN, as well as between the visual network and FPN. Thus, only ayahuasca/DMT was associated with negative total FC of the visual network (sum of FC between the visual network and all other networks), while total FC of the VAN was negative for only psilocybin, though this result was driven by the findings of just a single study (Barrett, Krimmel, et al., 2020) (**Figure 4**).

**Figure 4.**
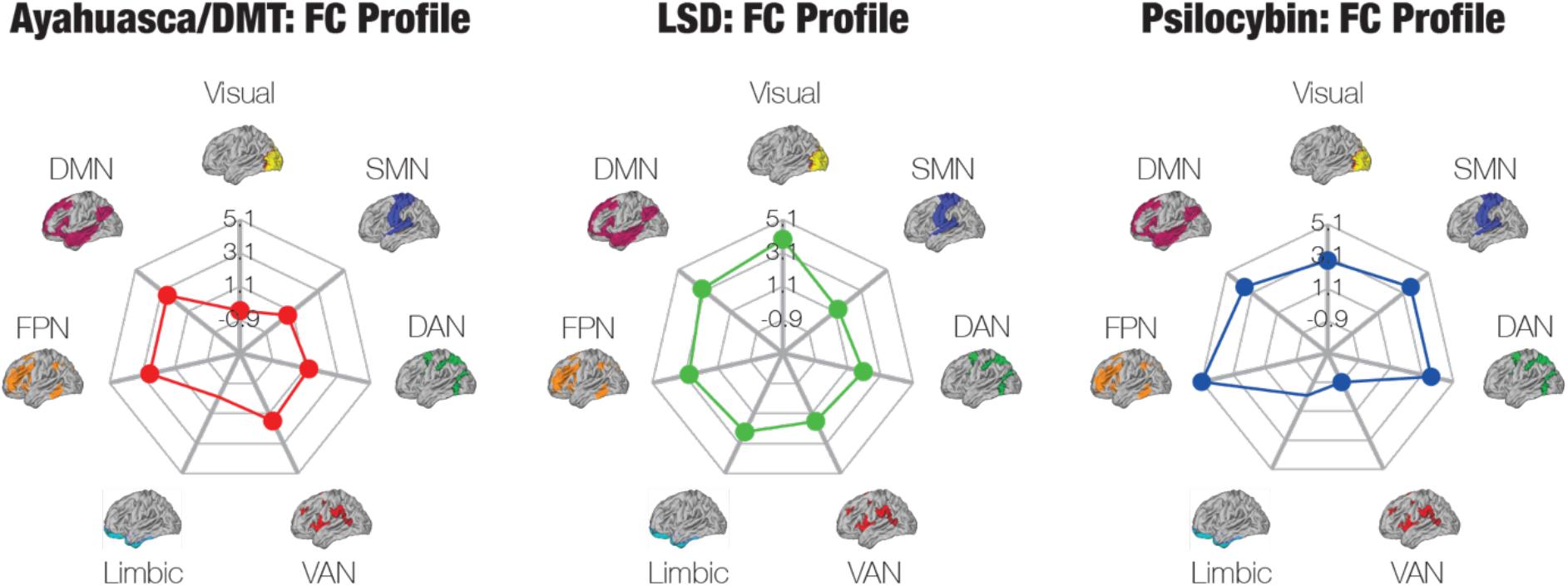
Functional connectivity (FC) profiles show unique FC patterns for each psychedelic. Out of the n = 12 studies that we examined in our quantitative FC meta-analysis, n = 3 were studies on ayahuasca/DMT (ayahuasca: n = 2; DMT: n = 1), n = 3 on LSD, and n = 6 on psilocybin. Each FC profile contains the total FC of each network, which was obtained by taking the sum of the rows of the corresponding aggregate FC matrices (Figure S5). (The units of the profiles are arbitrary.) The psychedelics display distinct FC profiles; for instance, LSD strongly elevates the total FC of the limbic network, whereas FC between the limbic network and all other networks was insignificant for ayahuasca/DMT and psilocybin (hence there is no point on the limbic network for the respective spider plots).

Figures S6-S8 display the results of our meta-analysis on the FC of subcortical regions. Because several studies did not measure subcortical FC, most of the subcortical-to-subcortical and subcortical-to-cortical edges in the aggregate FC matrix are insignificant. Nevertheless, the significant edges indicate that psychedelics elevate connectivity from two regions in the subcortex – the anterior thalamus and cerebellum – to cortex, as well as from the anterior thalamus to some other subcortical structures. Compared to the other psychedelics, LSD vastly elevated the FC between the anterior thalamus and both the cortex and subcortex.

Finally, there was no significant association between our results and any of the methodological covariates in the literature that we identified *a priori*: the width of the Gaussian kernel used to smooth the data, the use of FSL or SPM, the technique used to regress out white matter, the use of scrubbing to correct head motion, and route (oral or intravenous) and relative timing of drug administration. The results of our risk-of-bias assessment for individual studies is shown in **Table S6**. Because of the moderate-to-serious risk of bias, our certainty in the body of FC literature is low.

### Section 2.3. Pharmacology

We performed two separate meta-analyses on the pharmacology of psychedelics. The first assessed the selective affinity of DMT, LSD, and psilocin for the 5-HT_2A_, 5-HT_2C_, and D_2_ receptors, relative to 5-HT_1A_, as well as selective affinity for the 5-HT_1A_, 5-HT_2C_, and D_2_ receptors, relative to 5-HT_2A_. The second examined the relative functional activity of the psychedelics at the 5-HT_2A_ receptor, as captured in three different assays of GPCR signaling.

#### Section 2.3.1. Affinity

We first present our meta-analysis of the selective affinity data. Selectivity is defined as the ratio between the *K*_i_ of each psychedelic for a receptor of interest, relative to a reference receptor. *K*_i_ refers to the inhibition constant, which reflects the concentration of a drug that is needed to inhibit the binding of another ligand, for instance a radioactively-labeled ligand (radioligand), to the receptor of interest; it is inversely related to binding affinity (Sykes et al., 2019). We initially chose 5-HT_1A_ to be the reference receptor. We decided to analyse selectivity rather than absolute *K*_i_ because the latter is biased by the potency of the drug. Since LSD is much more potent than DMT and psilocin (Rickli et al., 2016), the meta-analysis would have simply revealed that LSD has higher affinity for all receptors if we were to examine the absolute *K*_i_ instead.

Our literature search identified *n* = 14 studies on selectivity, including *n* = 6 studies on DMT, *n* = 9 studies on LSD, and *n* = 5 studies on psilocin. We performed a random-effects meta-analysis, the results of which are displayed in Figure 5a (numerical data are given in **Table S7**). We found no significant between-drug differences in selectivity for any of the receptors – 5-HT_2A_, 5-HT_2C_, and D_2_ – relative to the 5-HT_1A_ receptor. Standard errors were very large for all of the pooled selectivity values, and heterogeneity was very high for the 5-HT_2A_ selectivity data (*I*^2^ = 93.69%) and for the 5-HT_2C_ selectivity data (*I*^2^ = 99.14%). Nevertheless, it is clear that all drugs display less selectivity for D_2_ than they do for 5-HT_2C_ and 5-HT_2A_. Each individual drug, especially LSD and psilocin, is more selective for 5-HT_2A_ than 5-HT_1A_. DMT and psilocin are about as equally selective for 5-HT_2C_ as they are for 5-HT_1A_, but LSD is much less selective for 5-HT_2C_ than for 5-HT_1A_.

**Figure 5.**
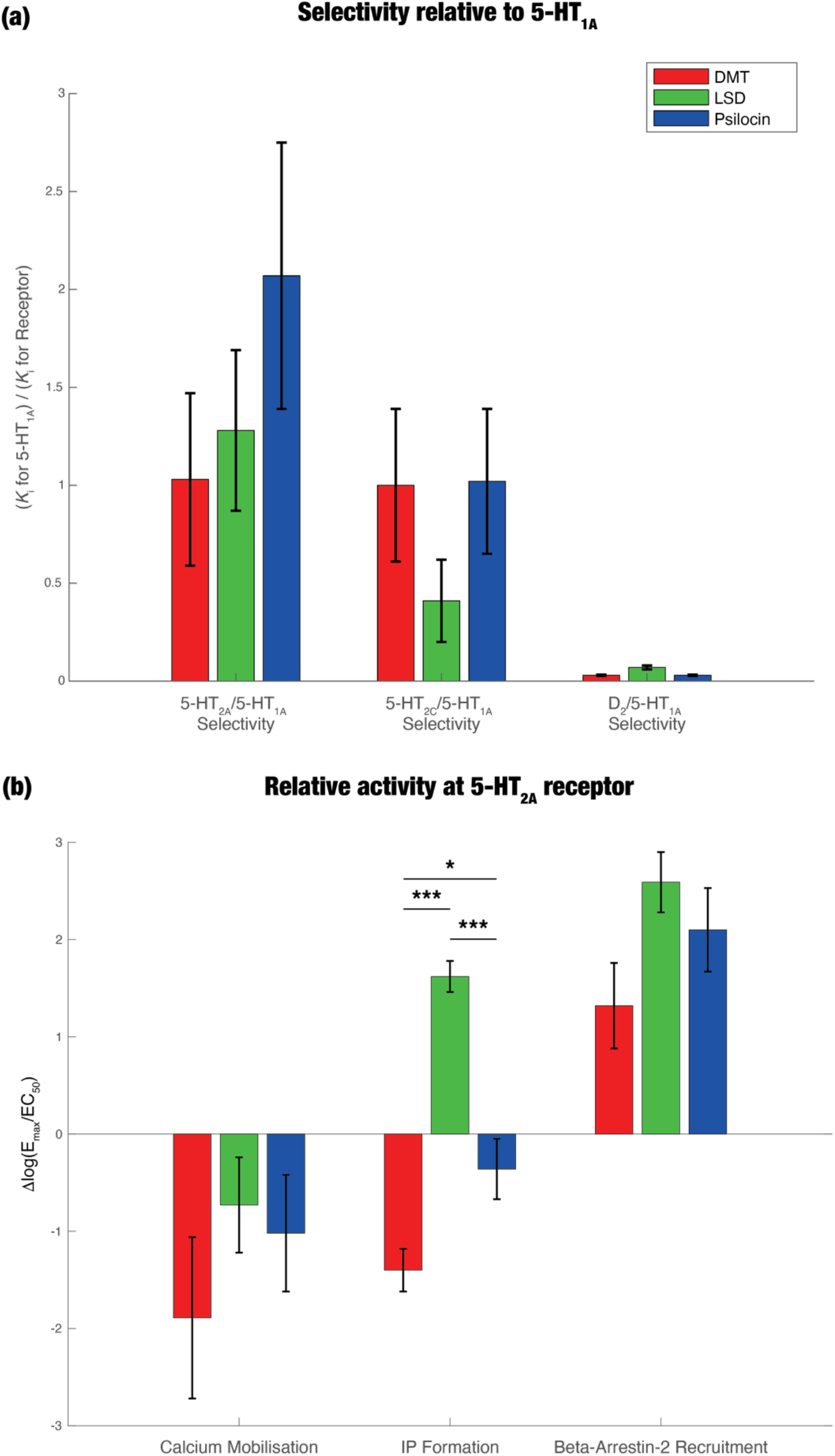
Pharmacology meta-analysis reveals that there are no significant differences in selectivity between psychedelics relative to 5-HT_1A_ and that LSD induces significantly higher relative activity at the inositol phosphate (IP) formation pathway. For both the selectivity and relative activity data, we created random-effects models that modeled between-study variance. (a) Selectivity is the ratio between the binding affinity (measured as K_i_) for a receptor of interest and the affinity for a reference receptor. We measured selectivity for three different receptors – 5-HT_2A_, 5-HT_2C_, and D_2_ – relative to 5-HT_1A_. Our literature search identified n = 14 studies on selectivity, including n = 6 studies on DMT, n = 9 studies on LSD, and n = 5 studies on psilocin. We did not find any significant between-drug differences in selectivity for any of the three receptors. (b) Relative activity is a measure of the cellular signalling that is elicited when a drug binds to a receptor. It is calculated here as Δlog(E_max_/EC_50_), where E_max_ is the maximal effect of the drug relative to a reference ligand and EC_50_ is the concentration needed to elicit 50% of the maximal effect (Kenakin, 2017). We found n = 18 studies on functional activity, including n = 6 studies on DMT, n = 13 studies on LSD, and n = 6 studies on psilocin. We measured relative activity at three different signaling pathways: calcium mobilisation, IP formation, and β-arrestin2 recruitment. IP formation was the only pathway that exhibited any significant between-drug differences; LSD elicited significantly higher activity than both DMT and psilocin. * < 0.05, ** < 0.01, *** < 0.001.

We constructed another random-effects model in which the radioligand for the receptor of interest was included as a covariate; we refer to this as the “full” model and the previous one, in which the influence of the radioligand was not modeled, as the “reduced” model. For 5-HT_2A_, the use of the radioligands [^3^H]-ketanserin (*p* = 0.0141) and [^3^H]-spiperone (*p* = 0.0271) significantly influenced the pooled selectivity estimates. Use of [^3^H]-mesulgerine (*p* = 0.0292), but not [^3^H]-ketanserin (*p* = 0.0872), significantly affected our estimates of selectivity for 5-HT_2C_. According to the corrected Akaike’s information criterion, the full model performed significantly better than the reduced model for 5-HT_2A_ selectivity (*p* < 0.0001), whereas the reverse was true for 5-HT_2C_ selectivity (*p* = 0.0024).

We assessed publication bias due to small study effects and found significant evidence for bias in the 5-HT_2A_ (*p* = 0.0001) and 5-HT_2C_ (*p* = 0.0004) affinity data. (There were not enough studies on D_2_ affinity to measure bias.) In *Section S6.3*, we qualitatively describe some major confounders in the affinity literature. Due to the large imprecision of the results and high probability of publication bias, our certainty in the body of evidence about selectivity is low.

However, when selectivity was measured relative to the 5-HT_2A_ receptor, there were significant between-drug differences (**Figure S9**; numerical results shown in **Table S8**). (Note that there were three additional studies, one for DMT and two for LSD.) In particular, DMT was significantly more selective for the 5-HT_2C_ receptor than both LSD (*p* = 0.001) and psilocin (*p* = 0.0035). In addition, selectivity for the D_2_ receptor was significantly higher for DMT than for psilocin (*p* = 0.001). Heterogeneity was once again high for the 5-HT_2C_ selectivity data (*I*^2^ = 98.56%) and for the 5-HT_1A_ selectivity (*I*^2^ = 99.91%). Surprisingly, there was no evidence for small-study bias in either the 5-HT_1A_ (*p* = 0.1683) or the 5-HT_2C_ case (*p* = 0.3604).

To relate the selective affinities to the neuroimaging and phenomenology of psychedelics, we first determined the expression of the 5-HT_2A_ and D_2_ receptors in the Yeo networks, based on an atlas of PET maps (Hansen et al., 2022) (unfortunately, this did not include maps of the 5-HT_2C_ receptor). Then, we defined the “pharmacology profile” as the weighted sum of the expression patterns, in which the weights were the selectivity of each psychedelic for the corresponding receptor, relative to the 5-HT_1A_ receptor (Figure 6). Because selectivity for the 5-HT_2A_ receptor is around two orders of magnitude higher than for the D_2_ receptor, the pharmacology profile predominantly reflects selectivity for 5-HT_2A_. Since psilocin’s selectivity for 5-HT_2A_ is greater than that of LSD and DMT, psilocin has the “largest” pharmacology profile. 5-HT_2A_ is expressed most in the DMN; thus, selectivity for 5-HT_2A_ may affect brain activity the most in the DMN. Subcortical pharmacology profiles are shown in **Figure S9** and indicate that 5-HT_2A_ receptor expression is highest in the hippocampus.

**Figure 6.**
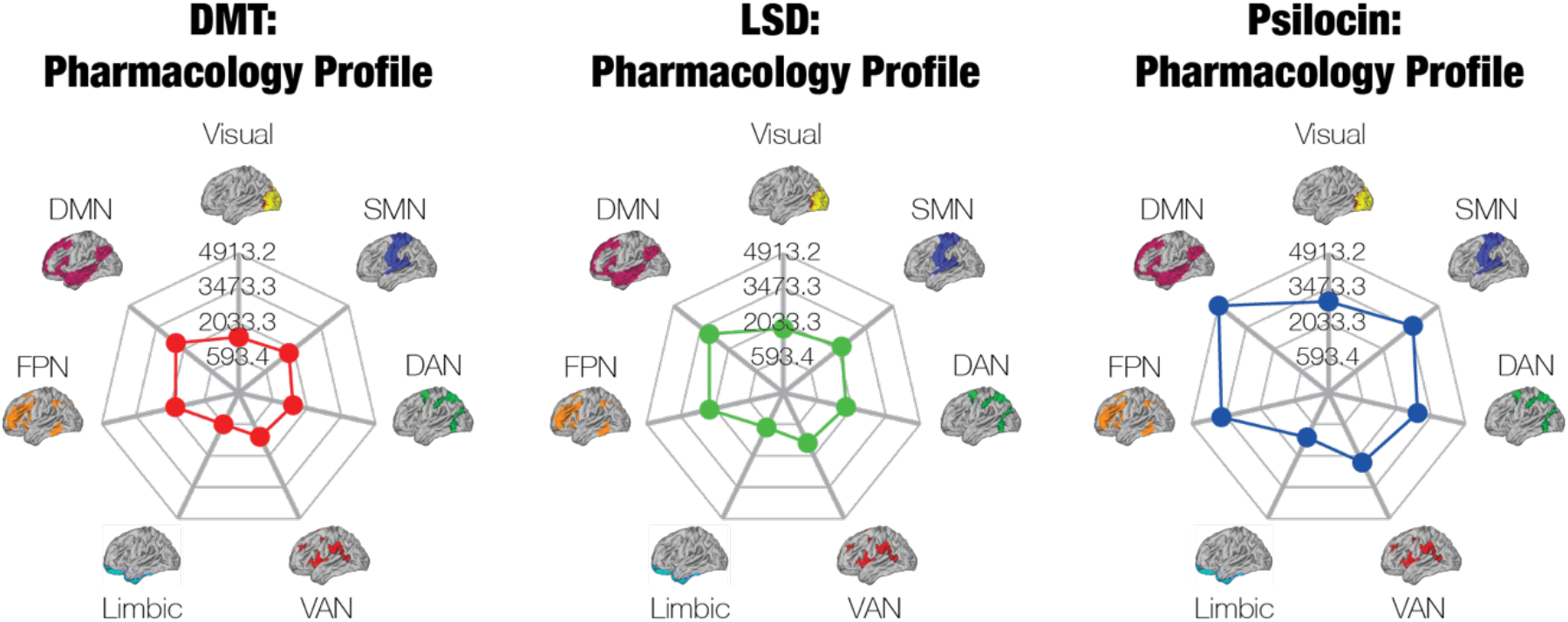
Pharmacology profiles primarily reveal the distribution of 5-HT_2A_ receptors in the Yeo networks, as well as the (insignificantly) higher selectivity of psilocin for the 5-HT_2A_ receptor compared to DMT and LSD. Based on an available PET atlas of the 5-HT_2A_ and D_2_ receptors (Hansen *et al*., 2022), we created the pharmacology profile of each psychedelic. The profiles show the expression of those two receptors in the Yeo networks, weighted by the selectivity of the corresponding psychedelics for those receptors. Because the selectivity for 5-HT_2A_ is two orders of magnitude higher than the selectivity for D_2_, the profiles are dominated by 5-HT_2A_ receptor expression. Since psilocin has the most selectivity for 5-HT_2A_ (relative to 5-HT_1A_), followed by LSD then DMT, psilocin’s pharmacology profile is the “largest.”

#### Section 2.3.2. Functional activity

Binding affinity is only one facet of the pharmacology of psychedelics. Another key aspect is their functional activity: the responses that they elicit in receptors after binding to them. While there are many ways to measure functional activity, we focused on GPCR signaling through three different pathways: (1) IP formation, (2) calcium mobilisation, and (3) β-arrestin2 recruitment. We limited our analysis to functional activity at specifically the 5-HT_2A_ receptor, as there was much more data about this receptor in humans. The variable of interest in our analysis was Δlog(*E*_max_/EC_50_).

We found *n =* 18 relevant studies on functional activity, including *n* = 6 studies on DMT, *n* = 13 studies on LSD, and *n* = 6 studies on psilocin. We performed a random-effects meta-analysis. For the IP formation and calcium mobilisation assays, the reference ligand was serotonin; for the β-arrestin2 recruitment assays, the reference ligand was chosen to be mescaline, as this was the only reference ligand used in experiments on all three psychedelics. The pooled relative activity values indicate that LSD induces significantly more IP formation than DMT (*p* < 0.0001) and psilocin (*p* = 0.0002) (Figure 5b; numerical data given in **Table S8**). Additionally, DMT elicits significantly more IP formation than psilocin (*p* = 0.0203). There were no significant between-drug differences for the other two pathways. Heterogeneity was very high for all three pathways (IP formation: *I*^2^ = 95.35%, calcium mobilisation: *I*^2^ = 92.66%, β-arrestin2 recruitment: *I*^2^ = 91.05%). There was no significant publication bias due to small study effects for the literature on IP formation (*p* = 0.3460). (There were too few studies on the other pathways to assess publication bias.) Because of the large standard errors of the pooled estimates and high unexplained heterogeneity, our certainty in the body of evidence about functional activity is low, despite the lack of evidence for publication bias.

## Section 3. Discussion

Here we examine the effects of psychedelics at three levels: 1) subjective experience (phenomenology), 2) functional connectivity (neuroimaging), and 3) the interaction of psychedelics with serotonin and dopamine receptors (pharmacology). At each level, we performed a quantitative meta-analysis and computed the alignment between the results and the seven Yeo networks. The latter network analysis enabled us to directly compare the effects of three psychedelics – ayahuasca/DMT, LSD, and psilocybin – within and between levels.

### Section 3.1. Unifying the levels of analysis

For the phenomenology literature, we conducted a meta-analysis of both the 5-dimensional and 11-dimensional versions of the Altered States of Consciousness (ASC) scale, a common questionnaire for measuring changes in subjective experience on psychedelics. At both medium and high doses, LSD ranked significantly higher than psilocybin in “visionary restructuralisation,” a dimension capturing the quality and intensity of visual hallucinations. Additionally, at medium doses, LSD was associated with significantly higher scores in the “oceanic boundlessness” dimension, which captures feelings of interconnectedness.

The whole-brain neuroimaging data consisted of three subsets of data: BOLD activation, entropy, and FC. We did not report a meta-analysis of the BOLD activation or entropy literature due to the heterogeneity of experimental procedures and analysis methods, respectively; however, we did conduct a qualitative review of the entropy literature (*Section S4*). To perform a meta-analysis on the FC data, we essentially re-parcellated the published data into the Yeo networks and then calculated a weighted sum of the connections between Yeo networks. LSD strongly elevated FC between the visual network and the other networks, whereas ayahuasca/DMT increased FC most between transmodal networks (specifically, the FPN and DMN) and unimodal networks (i.e. SMN and visual).

For pharmacology, we conducted two meta-analyses: one of selectivity and another of relative functional activity. There were no significant differences between psychedelics in selectivity for the 5-HT_2A_, 5-HT_2C_, or D_2_ receptors relative to the 5-HT_1A_ receptor. Compared to both DMT and psilocin, LSD elicited significantly more relative activity in IP formation assays. We did not observe any significant between-drug differences for the calcium mobilisation and β-arrestin2 recruitment assays.

Directly comparing the pharmacology, neuroimaging, and phenomenology profiles of psychedelics (Figure 7) reveals a weak one-to-one relationship between the three levels of analysis, which is unsurprising given the highly non-linear, complex, and reciprocal (Kringelbach et al., 2020) interactions between them. However, the results of the meta-analyses at each level may reveal some insights into the nature of these interactions.

**Figure 7.**
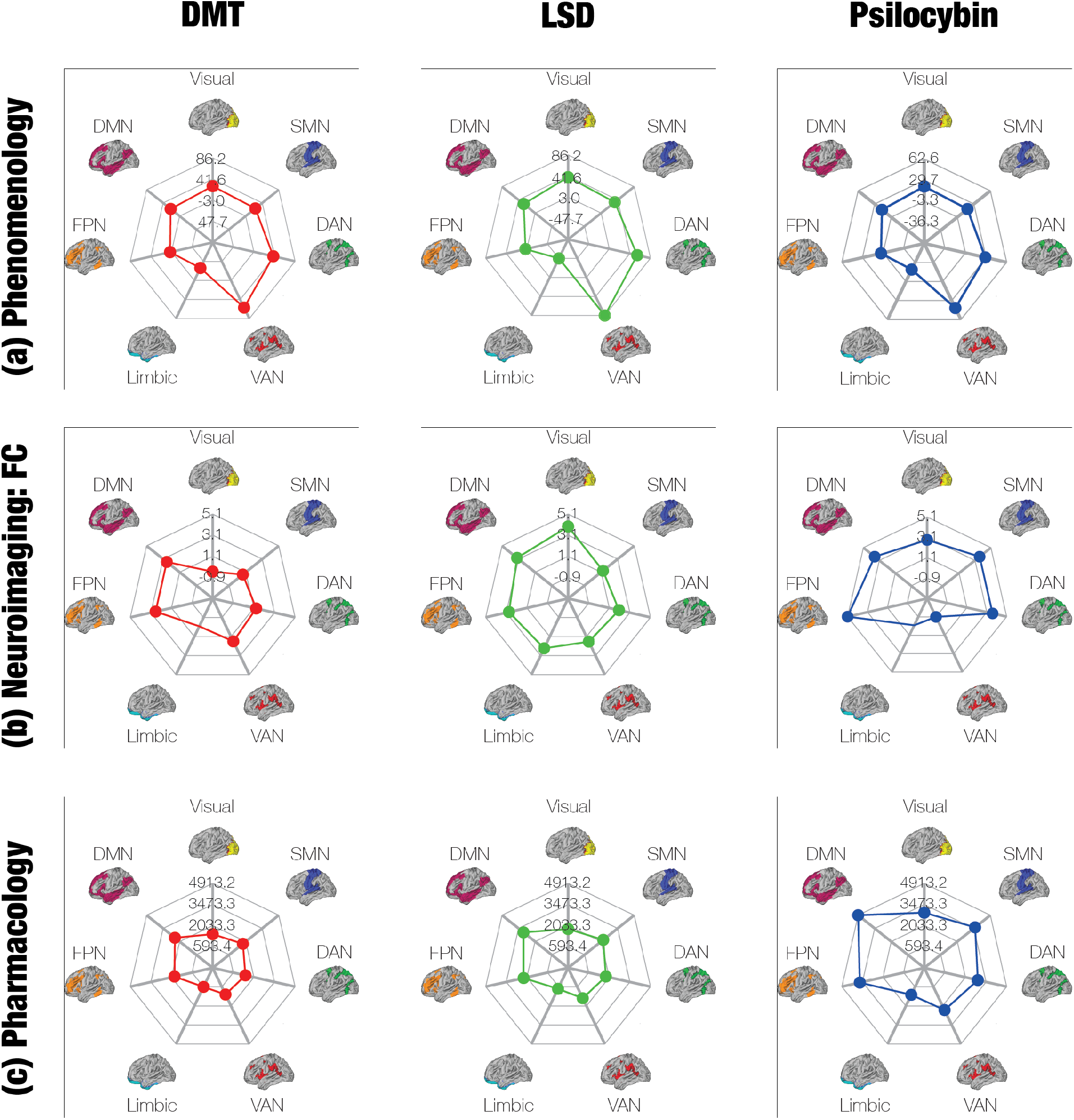
Our multi-level analysis of psychedelic effects highlights the nonlinear relationship between their pharmacology, neuroimaging, and phenomenology. The effects of psychedelics form a tripartite hierarchy, consisting of subjective experience (a, phenomenology), functional connectivity (b, neuroimaging), and their selective affinity for receptors (c, pharmacology). Here, we show the neural correlates of each level of the hierarchy in the seven Yeo networks. In brief, each profile was derived from (a) neural correlates of the subjective dimensions of the ASC scale, (b) summing the aggregate FC between each network and other networks, and (c) the distribution of each receptor, weighted by each psychedelic’s selectivity for that receptor. Clearly, there is a very weak correspondence between the different levels of the hierarchy, revealing the highly non-linear relationship between phenomenology, neuroimaging, and pharmacology. Note: we analysed data on both DMT and ayahuasca in our FC meta-analysis, but only on DMT for the other meta-analyses. We examined data on psilocin in our pharmacology meta-analysis and on psilocybin in both of the other meta-analyses.

On a superficial level, there do appear to be some direct correspondences between the phenomenology, neuroimaging, and pharmacology. Phenomenologically, we only found one significant difference between psychedelics that was consistent across medium and high doses: LSD induces more visionary restructuralisation than psilocybin does. Neurobiologically, LSD enhanced connectivity between the visual network and all other networks more than the other psychedelics.

The relationship between the pharmacology results and the above findings is more difficult to establish. Our results show that LSD increases IP formation, whereas DMT and psilocin decrease it. IP formation occurs whenever Gα_q_ proteins are recruited to initiate cellular signaling. Furthermore, there is evidence that Gα_q_ activation is necessary for 5-HT_2A_ agonists to bring about hallucinogenic effects. When administered the psychedelic (±)1-(2,5-dimethoxy-4-iodophenyl)-2-aminopropane (DOI), mice that lack the gene for Gα_q_ proteins do not exhibit behaviours (head twitches) that typically indicate hallucinations (Garcia et al., 2007). More recently, activation of Gα_q_ pathways by 5-HT_2A_ agonists has been shown to predict head-twitch behaviour, whereas 5-HT_2A_ agonists that are biased for the β-arrestin2 pathway do not induce head twitches (Wallach et al., 2023). There appears to be a threshold level of Gα_q_ activation that must be surpassed in order for a 5-HT_2A_ agonist to induce hallucinogenic effects (Wallach et al., 2023). Intriguingly, the pro-psychotic properties of drugs can be predicted from the extent to which they increase Gα_q_ activation (relative to a different G-protein pathway) when these drugs interact with a complex formed between the 5-HT_2A_ receptor and a metabotropic glutamate receptor (Fribourg et al., 2011). In other words, the more that a drug upsets the normal balance of Gα_q_ signaling, the more likely it is to engender hallucinations. However, these studies only establish the presence of hallucinations elicited by Gα_q_ signaling. They do not indicate whether Gα_q_ signaling induces *visual* hallucinations in particular. Thus, based on the available evidence, we cannot yet conclude that the pharmacology, neuroimaging, and phenomenology literature all convergently indicate that LSD has a uniquely strong effect on visual awareness, compared to the other psychedelics.

There do not appear to be any tools at the moment for relating the pharmacology of drugs to the precise nature of the subjective experiences that they produce. Methods for examining the relationship between neuroimaging and phenomenology are currently limited to correlations and linear regressions (Burt et al., 2021; Carhart-Harris et al., 2016), which do not capture causality. There is a need to develop techniques for elucidating the causal relationships between phenomenology, neuroimaging, and pharmacology.

### Section 3.2. Relating the results of the meta-analysis to the literature

#### Section 3.2.1. Phenomenology

We sought to verify the results of our phenomenology meta-analysis by comparing them to the two experimental studies that measured ASC ratings of LSD and psilocybin in the same group of participants. Holze et al. (2022) gave a medium (15 mg) and a high (30 mg) dose of psilocybin, as well as a medium (0.01 mg) and a high (0.02 mg) dose of LSD, and then compared both 5D- and 11D-ASC scores. It is worth noting that the doses of psilocybin and LSD in this study are not equivalent, since 0.01 mg LSD = 20 mg psilocybin (Ley et al., 2023). When comparing 15 mg psilocybin to 0.01 mg LSD, mean 5D-ASC and 11D-ASC scores tended to be higher for LSD than psilocybin. However, the only differences that reached significance were in the OB and VR scales for the 5D-ASC analysis, and in the complex imagery and audio-visual synaesthesia subscales for the 11D-ASC analysis. In our meta-analysis, LSD and psilocybin were significantly different in both OB and VR ratings, as well as in complex imagery and audio-visual synaesthesia ratings, at medium doses. Ley et al. (2023) compared the subjective effects of medium doses of LSD (0.01 mg) and psilocybin (20 mg). Here, the doses are equivalent, unlike in the previous study. The study found no significant differences between LSD and psilocybin in any of the dimensions, both in the 5D and 11D analyses. Therefore, the results of our meta-analysis are not entirely supported by studies that conducted direct comparisons between psychedelics.

#### Section 3.2.2. Neuroimaging

One major finding in the neuroimaging literature is that psychedelics desegregate and disintegrate brain networks. Segregation is defined as the lack of FC *between* brain regions, while integration denotes FC *within* a network. Carhart-Harris et al. (2016) found that LSD led to disintegration and desegregation for most RSNs. This finding has been independently confirmed in several other studies (Barrett, Doss, et al., 2020; Dai et al., 2023; Girn et al., 2022; Lebedev et al., 2015; Madsen et al., 2021; Müller, Dolder, et al., 2018; Roseman et al., 2014). In a review of three studies, Müller, Liechti, et al. (2018) found good convergence in reports of desegregation. A recent study, which was published after our meta-analysis was completed, utilised a novel logitudinal precision functional mapping approach, in which participants were scanned 18 times before, during, and after psilocybin administration (Siegel et al., 2024). It also reported massive decreases in anticorrelations between networks (desegregation), as well as in correlations within networks (disintegration). Our meta-analysis corroborates the desegregation hypothesis and provides mixed support for the disintegration hypothesis. Psychedelics significantly elevated FC between all non-identical pairs of brain networks except the FPN and VAN. Within-network FC significantly decreased for the visual network, VAN, and DMN, yet it significantly increased for the DAN and FPN. The reduction in within-VAN FC and amplification of within-DAN FC may be consistent with the behavioural neurophysiology and phenomenology of psychedelics, respectively. Broadly speaking, the VAN redirects attention to salient stimuli, whereas the DAN is responsible for sustaining attention (Vossel et al., 2014). In agreement with the VAN FC results, there is some evidence, albeit mixed, that psychedelics reduce mismatch negativity, or the brain’s response to surprising stimuli (Duerler et al., 2021; Heekeren et al., 2008; Timmermann et al., 2018), though the MMN is primarily generated by regions outside the VAN (Garrido et al., 2009). Whereas closed-eyes mental imagery tends to be very fleeting when people are sober, it is often much more stable and vivid on psychedelics (Dittrich, 1998; Kometer & Vollenweider, 2018; Studerus et al., 2010). We speculate that this increase in stability could be attributed to a vast enhancement of the ability to freely allocate attentional resources on psychedelics, as has been proposed in the past (Lyon, 2023). That is, external stimuli typically demand attention, which limits the brain’s capacity to freely allocate attention to spontaneously generated, closed-eyes mental imagery. Psychedelics may reduce the competition between external stimuli and closed-eyes mental imagery for sustained attention, which could explain an increase in the within-network FC of the DAN. Intriguingly, ayahuasca and DMT are the only psychedelics to reduce FC between the visual network and the DAN. This could be consistent with the fact that only ayahuasca and DMT appear to give rise to *open-eye* “breakthrough” experiences, which feature extraordinarily rich and realistic visual hallucinations (Kometer & Vollenweider, 2018). Indeed, the loss of connectivity from the visual network, and consequently of visual input from the environment, may make it possible to sustain attention on complex visual hallucinations even while the eyes are open.

Meanwhile, increases in connectivity within the FPN could account for the therapeutic effects of psychedelics. The FPN is a network that initiates and flexibly adjusts cognitive control in response to feedback from the environment (Marek & Dosenbach, 2018). Depression compromises executive control and cognitive flexibility (Gabrys et al., 2018; Kraft et al., 2023); hence, it reduces FC within the FPN (Alexopoulos et al., 2012; Hwang et al., 2015). Psilocybin has shown promise for treating depression (Davis et al., 2021; Goodwin et al., 2022; Raison et al., 2023; Tabaac et al., 2024; Von Rotz et al., 2023), and while this meta-analysis was only conducted on data from healthy participants, our finding that psychedelics elevate within-FPN FC may still be able to explain their antidepressant effects.

The significant increases in within-FPN and within-DAN FC indicate that the effects of psychedelics on within-network FC are more nuanced than total disintegration across the whole brain. Furthermore, the explanatory power of desegregation and disintegration remains unclear. While there is a correlation between desegregation/disintegration and the subjective experience of ego dissolution (Tagliazucchi et al., 2016), there is no evidence for a *causal* link. There appear to be parallels between the conscious experience of interconnectedness and the “interconnectedness” of the brain on psychedelics, but further research is needed to demonstrate that the connection between the two is more than merely semantic.

Another common finding in the literature is that psychedelics decrease activity and connectivity within the DMN, a network that is generally implicated in self-awareness, self-reflection, and other self-referential cognitive processes (Davey et al., 2016; Davey & Harrison, 2018; Molnar-Szakacs & Uddin, 2013; Moran et al., 2013). This popular ’meme’ likely has its origin in the first resting-state fMRI study of a psychedelic, where decreased blood flow and BOLD signal were observed in a pattern of regions resembling the DMN (Carhart-Harris et al., 2012). Eleven years later, evidence continues to indicate that psychedelics potently affect the regularity of population brain activity in DMN regions (Timmermann et al., 2023). Since a commonly reported experience on psychedelics is the dissolution of the self, a phenomenon known as “ego death” (Moreton et al., 2020; Nour et al., 2016; Pahnke, 1969), it would superficially make sense that psychedelics dysregulate the DMN. Many studies have found significant correlations between subjective ratings of ego dissolution and DMN disintegration (Atasoy et al., 2017; Barrett, Krimmel, et al., 2020; Luppi et al., 2021; Madsen et al., 2021; Palhano-Fontes et al., 2015; Pasquini et al., 2020; Roseman et al., 2014; Smigielski et al., 2019; Tagliazucchi et al., 2014, 2016), though other analyses did not find evidence for a relationship between the two (Lebedev et al., 2015; Müller, Dolder, et al., 2018).

However, claims of “de-activation” or worse, “shutting-off” of the DMN as signatures of psychedelic action are wrong and misleading. Other neuroimaging studies by Carhart-Harris et al. (2012, 2013, 2016) and Roseman et al. (2014) discovered that psychedelics affect the *connectivity* of the DMN, decreasing within-DMN FC (“disintegration”) while increasing DMN coupling with other resting-state networks (RSNs) (“desegregation”). Our neuroimaging meta-analysis confirms that psychedelics significantly decrease connectivity within the DMN while significantly elevating its connectivity with all other networks. Furthermore, the reduction in within-network FC was greater for the DMN than for any other network, though it was not that much larger than that of the visual network. Although we found that LSD increased within-DMN FC, this result is likely attributable to connectivity between regions that are not key hubs of the DMN. Our phenomenology meta-analysis revealed that all psychedelics induced positive experiences of depersonalisation and derealisation, as captured by the oceanic boundlessness dimension of the 5D-ASC scale; these experiences are related to the feeling of ego death.

Nevertheless, there are many cases in which within-DMN connectivity may decrease without giving rise to the subjective effects that characterise psychedelics. Several other drugs such as amphetamines and alcohol, as well as drug addiction in general, reduce within-DMN connectivity in humans (Schrantee et al., 2016; Weber et al., 2014; Zhang & Volkow, 2019), even though they typically do not lead to ego dissolution (Doss et al., 2020). Certain psychological states that exacerbate self-rumination, such as maladaptive self-focused attention, are *not* associated with hyperconnectivity in the DMN (Fang et al., 2022), and one study found that some tasks requiring self-related judgments were associated with a *suppression* of within-DMN connectivity (van Buuren et al., 2010). That being said, a recent meta-analysis of 14 studies did endorse an association between rumination and the DMN (Zhou et al., 2020).

Finally, it is worth noting that a recent systematic review of the neuroimaging literature on psychedelics concluded that it was not possible to perform a meta-analysis due to the heterogeneity of the studies, as well as other methodological concerns (Linguiti et al., 2023). Overall, we agree that it is challenging to perform a standard meta-analysis on the psychedelic neuroimaging literature. However, several steps were taken in our analysis to overcome the concerns that Linguiti and colleagues raised. Unlike Linguiti and colleagues, we separated studies on functional connectivity from studies on entropy and BOLD activation. As stated above, the BOLD activation studies subjected participants to such a wide variety of tasks that it was impossible to conduct a meta-analysis on this segment of the literature. However, almost all of the functional connectivity studies that we included in our meta-analysis conducted resting-state recordings. Linguiti and colleagues correctly note that, even among the resting-state recordings, there experimental procedures and analysis methods are very heterogeneous. We ran subgroup analyses to confirm that these covariates, such as route of administration (oral or intravenous), timing of drug administration relative to the duration of the drug’s acute effects, and various preprocessing techniques, did not impact the results of our meta-analysis. Linguiti and colleagues also observed that there is a high sample overlap in the published neuroimaging studies; that is, there are many studies that reused or re-analysed data from previously published research. In our meta-analysis, we ensured that we included only one analysis of each unique dataset in the literature. Because of this and other exclusion criteria, we entered data from only 12 studies into our quantitative meta-analysis, whereas Linguiti and colleagues included 91 studies. We do agree with Linguiti and colleagues that at least one of our included studies failed to adequately control for Type I error (Kaelen et al., 2016) and that at least another one did not sufficiently report head motion correction (Grimm et al., 2018). Nevertheless, overall, we believe that there is enough consistency among some studies in the neuroimaging literature to conduct a meaningful meta-analysis.

#### Section 3.2.3. Pharmacology

The results of our affinity meta-analysis aligned well with studies that performed direct comparisons of selectivity between different psychedelics (Eshleman et al., 2014; Janowsky et al., 2014; Kozell et al., 2023; Pierce & Peroutka, 1989; Rickli et al., 2016). In line with our findings, all of these studies showed that LSD is more selective than DMT for 5-HT_2A_, relative to 5-HT_1A_. Two studies also observed that psilocin is more selective than LSD for 5-HT_2A_ (Kozell et al., 2023; Rickli et al., 2016). Our meta-analysis demonstrated that DMT is more selective than LSD for 5-HT_2C_, a conclusion that is supported by three of four studies (Eshleman et al., 2014; Janowsky et al., 2014; Kozell et al., 2023). Note that none of these studies measured the statistical significance of between-drug differences in selectivity.

Our findings about functional activity were also congruent with studies that measured the effect of multiple psychedelics on IP formation at the 5-HT_2A_ receptor (Braden & Nichols, 2007; Eshleman et al., 2014; Janowsky et al., 2014; Kozell et al., 2023). We found that LSD’s relative activity at this pathway was significantly greater than that of psilocin, which was in turn significantly larger than that of DMT. All four studies showed that LSD had higher relative activity at this pathway than DMT. Two studies also reported that psilocin’s relative activity was lower than that of LSD, but larger than that of DMT (Braden & Nichols, 2007; Kozell et al., 2023).

### Section 3.3. Recommendations for future research

We encourage researchers to develop tools for modelling the nonlinear relationships between the pharmacology, neuroimaging, and phenomenology of psychedelics, the three hierarchical levels of analysis that are at the heart of this meta-analysis. While some researchers have attempted to examine the associations between each level (Ballentine et al., 2022; Lawn et al., 2022), their techniques are model-free and only measure *correlations* between the effects of psychedelics, rather than elucidating the *causal mechanisms* that underpin them.

Recent whole-brain models have made significant advances in discovering these causal mechanisms (see Kringelbach & Deco (2020) for a general review). Whole-brain models consist of nodes that approximate local or mesoscopic neuronal dynamics through mean field models of coupled excitatory-inhibitory interactions (Deco et al., 2014; Honey et al., 2007) or Hopf models of bifurcations into and out of sustained oscillatory activity (Deco et al., 2017; Freyer et al., 2012). Crucially, whole-brain models have enabled researchers to identify the specific regions that should be stimulated in order to force transitions between different states of consciousness, such as from sleep to wakefulness (Deco et al., 2019). Whole-brain models have also been applied to psychedelics; in particular, the 5-HT_2A_ receptor density in each area of the brain was used to fit a mean field model of fMRI data on LSD, capturing the nonlinear interactions between connectivity and the distribution of 5-HT_2A_ receptors (Deco et al., 2018). This model was recently extended to determine the associations between global brain connectivity on LSD and categories of subjective experience (Burt et al., 2021). The model has also been applied to the relationship between neurotransmitters (namely, serotonin) and neuronal regions on psilocybin; in particular, Kringelbach et al. (2020) coupled a mean field model of each neuronal region to the release-and-reuptake dynamics of serotonin concentration. Importantly, this coupled neuronal-neurotransmitter model exhibited a better fit to the empirical FC than a model of solely the neuronal activity. Overall, whole-brain models display great promise for linking neurotransmitter release, which is partly determined by pharmacology, to the brain regions that causally mediate, rather than merely correlate with, the effects of psychedelics. Most recently, whole-brain models have also been combined with another cutting-edge method, turbulence, to determine the susceptibility of the brain to external stimuli under psychedelics (Cruzat et al., 2022).

This meta-analysis was only performed on data from healthy participants, yet there is emerging research on the neuroimaging and phenomenology of psychedelics in clinical populations (R. L. Carhart-Harris et al., 2017; Doss et al., 2021; Roseman, Nutt, et al., 2018). Future models that bridge the three levels of analysis should also be applied to the clinical data in order to shed light on the therapeutic mechanisms of psychedelics. It is essential to determine whether these models would be able to predict clinical outcomes. Recent research has succeeded in predicting treatment response to psilocybin for depression based on changes in the functional hierarchy of the brain (Deco et al., 2024). These methods may be even more effective if they incorporated data on phenomenology and pharmacology. However, as mentioned in the Introduction, psychedelic treatments are always administered in conjunction with therapy or some form of “psychological support,” which the models must also take into account.

## Section 4. Conclusion

This is the first meta-analysis of the literature on the phenomenology, neuroimaging, and pharmacology of psychedelics. We assess and compare these three hierarchical levels of analysis across three different classical psychedelics: DMT, LSD, and psilocybin. Thus, we performed comparisons not only between psychedelics, but also between the levels of the hierarchy.

We found that the different levels of the hierarchy exhibited a weak one-to-one correspondence with one another. This is not surprising, as the interactions between phenomenology, neuroimaging, and pharmacology are highly non-linear and complex. We encourage future research to develop tools for modelling these relationships in order to improve our scientific understanding of psychedelics.

## Acknowledgments

K.S. is funded by the Clarendon Fund, a Department of Psychiatry studentship, the John Henry Jones Scholarship from Balliol College, and the Research Council U.K. P.M. is funded by the Wellcome Trust (grant no. 210920/Z/18/Z). K.P. is funded by the Swiss National Science Foundation (P2ZHP1\161626). R.C-H is funded by a Ralph Metzner endowment. M.K. is supported by the Centre for Eudaimonia and Human Flourishing (funded by the Pettit and Carlsberg Foundations) and Center for Music in the Brain (funded by the Danish National Research Foundation, DNRF117).

The authors would like to thank Michael Angyus for his help with improving the methodology for the functional connectivity meta-analysis.

## Code and Data Availability

Code and data used for the phenomenology and pharmacology analyses are available upon request by emailing the corresponding author

## Study Registration and Protocol

This study was not registered. A review protocol was not prepared.

## References

Alexopoulos, G. S., Hoptman, M. J., Kanellopoulos, D., Murphy, C. F., Lim, K. O., & Gunning, F. M. (2012). Functional connectivity in the cognitive control network and the default mode network in late-life depression. Journal of Affective Disorders, 139(1), 56–65. 10.1016/j.jad.2011.12.002

Andresen, B. T. (2011). A Pharmacological Primer of Biased Agonism. Endocrine, Metabolic & Immune Disorders Drug Targets, 11(2), 92–98. 10.2174/187153011795564179

Atasoy, S., Roseman, L., Kaelen, M., Kringelbach, M. L., Deco, G., & Carhart-Harris, R. L. (2017). Connectome-harmonic decomposition of human brain activity reveals dynamical repertoire re-organization under LSD. Scientific Reports, 7(1), Article 1. 10.1038/s41598-017-17546-0

Ballentine, G., Friedman, S. F., & Bzdok, D. (2022). Trips and neurotransmitters: Discovering principled patterns across 6850 hallucinogenic experiences. Science Advances, 8(11), eabl6989. 10.1126/sciadv.abl6989

Barrett, F. S., Doss, M. K., Sepeda, N. D., Pekar, J. J., & Griffiths, R. R. (2020). Emotions and brain function are altered up to one month after a single high dose of psilocybin. Scientific Reports, 10(1), 2214. 10.1038/s41598-020-59282-y

Barrett, F. S., Krimmel, S. R., Griffiths, R. R., Seminowicz, D. A., & Mathur, B. N. (2020). Psilocybin acutely alters the functional connectivity of the claustrum with brain networks that support perception, memory, and attention. NeuroImage, 218, 116980. 10.1016/j.neuroimage.2020.116980

Bedford, P., Hauke, D. J., Wang, Z., Roth, V., Nagy-Huber, M., Holze, F., Ley, L., Vizeli, P., Liechti, M. E., Borgwardt, S., Müller, F., & Diaconescu, A. O. (2023). The effect of lysergic acid diethylamide (LSD) on whole-brain functional and effective connectivity. Neuropsychopharmacology, 48(8), Article 8. 10.1038/s41386-023-01574-8

Beliveau, V., Ganz, M., Feng, L., Ozenne, B., Højgaard, L., Fisher, P. M., Svarer, C., Greve, D. N., & Knudsen, G. M. (2017). A High-Resolution In Vivo Atlas of the Human Brain’s Serotonin System. The Journal of Neuroscience: The Official Journal of the Society for Neuroscience, 37(1), 120–128. 10.1523/JNEUROSCI.2830-16.2016

Bohn, L. M., & Schmid, C. L. (2010). Serotonin receptor signaling and regulation via β-arrestins. Critical Reviews in Biochemistry and Molecular Biology, 45(6), 555. 10.3109/10409238.2010.516741

Braden, M. R., & Nichols, D. E. (2007). Assessment of the roles of serines 5.43(239) and 5.46(242) for binding and potency of agonist ligands at the human serotonin 5-HT2A receptor. Molecular Pharmacology, 72(5), 1200–1209. 10.1124/mol.107.039255

Burt, J. B., Preller, K. H., Demirtas, M., Ji, J. L., Krystal, J. H., Vollenweider, F. X., Anticevic, A., & Murray, J. D. (2021). Transcriptomics-informed large-scale cortical model captures topography of pharmacological neuroimaging effects of LSD. eLife, 10, e69320. 10.7554/eLife.69320

Canal, C. E., & Murnane, K. S. (2017). The serotonin 5-HT2C receptor and the non-addictive nature of classic hallucinogens. Journal of Psychopharmacology, 31(1), 127–143. 10.1177/0269881116677104

Carbonaro, T. M., Johnson, M. W., Hurwitz, E., & Griffiths, R. R. (2018). Double-blind comparison of the two hallucinogens psilocybin and dextromethorphan: Similarities and differences in subjective experiences. Psychopharmacology, 235(2), 521–534. 10.1007/s00213-017-4769-4

Carhart-Harris, R. L. (2018). The entropic brain—Revisited. Neuropharmacology, 142, 167–178. 10.1016/j.neuropharm.2018.03.010

Carhart-Harris, R. L., Erritzoe, D., Williams, T., Stone, J. M., Reed, L. J., Colasanti, A., Tyacke, R. J., Leech, R., Malizia, A. L., Murphy, K., Hobden, P., Evans, J., Feilding, A., Wise, R. G., & Nutt, D. J. (2012). Neural correlates of the psychedelic state as determined by fMRI studies with psilocybin. Proceedings of the National Academy of Sciences, 109(6), 2138–2143. 10.1073/pnas.1119598109

Carhart-Harris, R. L., Leech, R., Erritzoe, D., Williams, T. M., Stone, J. M., Evans, J., Sharp, D. J., Feilding, A., Wise, R. G., & Nutt, D. J. (2013). Functional connectivity measures after psilocybin inform a novel hypothesis of early psychosis. Schizophrenia Bulletin, 39(6), 1343–1351. 10.1093/schbul/sbs117

Carhart-Harris, R. L., Muthukumaraswamy, S., Roseman, L., Kaelen, M., Droog, W., Murphy, K., Tagliazucchi, E., Schenberg, E. E., Nest, T., Orban, C., Leech, R., Williams, L. T., Williams, T. M., Bolstridge, M., Sessa, B., McGonigle, J., Sereno, M. I., Nichols, D., Hellyer, P. J., … Nutt, D. J. (2016). Neural correlates of the LSD experience revealed by multimodal neuroimaging. Proceedings of the National Academy of Sciences of the United States of America, 113(17), 4853–4858. 10.1073/pnas.1518377113

Carhart-Harris, R. L., Roseman, L., Bolstridge, M., Demetriou, L., Pannekoek, J. N., Wall, M. B., Tanner, M., Kaelen, M., McGonigle, J., Murphy, K., Leech, R., Curran, H. V., & Nutt, D. J. (2017). Psilocybin for treatment-resistant depression: fMRI-measured brain mechanisms. Scientific Reports, 7(1), Article 1. 10.1038/s41598-017-13282-7

Carhart-Harris, R., Leech, R., Hellyer, P., Shanahan, M., Feilding, A., Tagliazucchi, E., Chialvo, D., & Nutt, D. (2014). The entropic brain: A theory of conscious states informed by neuroimaging research with psychedelic drugs. Frontiers in Human Neuroscience, 8. https://www.frontiersin.org/article/10.3389/fnhum.2014.00020

Coyle, J. R., Presti, D. E., & Baggott, M. J. (2012). Quantitative Analysis of Narrative Reports of Psychedelic Drugs (arXiv:1206.0312). arXiv. 10.48550/arXiv.1206.0312

Cruzat, J., Perl, Y. S., Escrichs, A., Vohryzek, J., Timmermann, C., Roseman, L., Luppi, A. I., Ibañez, A., Nutt, D., Carhart-Harris, R., Tagliazucchi, E., Deco, G., & Kringelbach, M. L. (2022). Effects of classic psychedelic drugs on turbulent signatures in brain dynamics. Network Neuroscience, 6(4), 1104– 1124. 10.1162/netn_a_00250

Dai, R., Larkin, T. E., Huang, Z., Tarnal, V., Picton, P., Vlisides, P. E., Janke, E., McKinney, A., Hudetz, A. G., Harris, R. E., & Mashour, G. A. (2023). Classical and non-classical psychedelic drugs induce common network changes in human cortex. NeuroImage, 273, 120097. 10.1016/j.neuroimage.2023.120097

Davey, C. G., & Harrison, B. J. (2018). The brain’s center of gravity: How the default mode network helps us to understand the self. World Psychiatry, 17(3), 278–279. 10.1002/wps.20553

Davey, C. G., Pujol, J., & Harrison, B. J. (2016). Mapping the self in the brain’s default mode network. NeuroImage, 132, 390–397. 10.1016/j.neuroimage.2016.02.022

Davis, A. K., Barrett, F. S., May, D. G., Cosimano, M. P., Sepeda, N. D., Johnson, M. W., Finan, P. H., & Griffiths, R. R. (2021). Effects of Psilocybin-Assisted Therapy on Major Depressive Disorder: A Randomized Clinical Trial. JAMA Psychiatry, 78(5), 481–489. 10.1001/jamapsychiatry.2020.3285

Daws, R. E., Timmermann, C., Giribaldi, B., Sexton, J. D., Wall, M. B., Erritzoe, D., Roseman, L., Nutt, D., & Carhart-Harris, R. (2022). Increased global integration in the brain after psilocybin therapy for depression. Nature Medicine, 28(4), Article 4. 10.1038/s41591-022-01744-z

de Almeida, J., & Mengod, G. (2007). Quantitative analysis of glutamatergic and GABAergic neurons expressing 5-HT(2A) receptors in human and monkey prefrontal cortex. Journal of Neurochemistry, 103(2), 475–486. 10.1111/j.1471-4159.2007.04768.x

de Oliveira, P. G., Ramos, M. L. S., Amaro, A. J., Dias, R. A., & Vieira, S. I. (2019). Gi/o-Protein Coupled Receptors in the Aging Brain. Frontiers in Aging Neuroscience, 11. https://www.frontiersin.org/articles/10.3389/fnagi.2019.00089

de Vos, C. M. H., Mason, N. L., & Kuypers, K. P. C. (2021). Psychedelics and Neuroplasticity: A Systematic Review Unraveling the Biological Underpinnings of Psychedelics. Frontiers in Psychiatry, 12. https://www.frontiersin.org/article/10.3389/fpsyt.2021.724606

Deco, G., Cruzat, J., Cabral, J., Knudsen, G. M., Carhart-Harris, R. L., Whybrow, P. C., Logothetis, N. K., & Kringelbach, M. L. (2018). Whole-Brain Multimodal Neuroimaging Model Using Serotonin Receptor Maps Explains Non-linear Functional Effects of LSD. Current Biology, 28(19), 3065–3074.e6. 10.1016/j.cub.2018.07.083

Deco, G., Cruzat, J., Cabral, J., Tagliazucchi, E., Laufs, H., Logothetis, N. K., & Kringelbach, M. L. (2019). Awakening: Predicting external stimulation to force transitions between different brain states. Proceedings of the National Academy of Sciences, 116(36), 18088–18097. 10.1073/pnas.1905534116

Deco, G., Kringelbach, M. L., Jirsa, V. K., & Ritter, P. (2017). The dynamics of resting fluctuations in the brain: Metastability and its dynamical cortical core. Scientific Reports, 7(1), 3095. 10.1038/s41598-017-03073-5

Deco, G., Ponce-Alvarez, A., Hagmann, P., Romani, G. L., Mantini, D., & Corbetta, M. (2014). How Local Excitation–Inhibition Ratio Impacts the Whole Brain Dynamics. Journal of Neuroscience, 34(23), 7886– 7898. 10.1523/JNEUROSCI.5068-13.2014

Deco, G., Sanz Perl, Y., Johnson, S., Bourke, N., Carhart-Harris, R. L., & Kringelbach, M. L. (2024). Different hierarchical reconfigurations in the brain by psilocybin and escitalopram for depression. Nature Mental Health, 1–15. 10.1038/s44220-024-00298-y

Dittrich, A. (1998). The standardized psychometric assessment of altered states of consciousness (ASCs) in humans. Pharmacopsychiatry, 31 Suppl 2, 80–84. 10.1055/s-2007-979351

Doss, M. K., May, D. G., Johnson, M. W., Clifton, J. M., Hedrick, S. L., Prisinzano, T. E., Griffiths, R. R., & Barrett, F. S. (2020). The Acute Effects of the Atypical Dissociative Hallucinogen Salvinorin A on Functional Connectivity in the Human Brain. Scientific Reports, 10(1), Article 1. 10.1038/s41598-020-73216-8

Doss, M. K., Považan, M., Rosenberg, M. D., Sepeda, N. D., Davis, A. K., Finan, P. H., Smith, G. S., Pekar, J. J., Barker, P. B., Griffiths, R. R., & Barrett, F. S. (2021). Psilocybin therapy increases cognitive and neural flexibility in patients with major depressive disorder. Translational Psychiatry, 11(1), Article 1. 10.1038/s41398-021-01706-y

Duerler, P., Brem, S., Fraga-González, G., Neef, T., Allen, M., Zeidman, P., Stämpfli, P., Vollenweider, F. X., & Preller, K. H. (2021). Psilocybin Induces Aberrant Prediction Error Processing of Tactile Mismatch Responses-A Simultaneous EEG-FMRI Study. Cerebral Cortex (New York, N.Y.: 1991), 32(1), 186–196. 10.1093/cercor/bhab202

Eshleman, A. J., Forster, M. J., Wolfrum, K. M., Johnson, R. A., Janowsky, A., & Gatch, M. B. (2014). Behavioral and neurochemical pharmacology of six psychoactive substituted phenethylamines: Mouse locomotion, rat drug discrimination and in vitro receptor and transporter binding and function. Psychopharmacology, 231(5), 875–888. 10.1007/s00213-013-3303-6

Fang, A., Baran, B., Beatty, C. C., Mosley, J., Feusner, J. D., Phan, K. L., Wilhelm, S., & Manoach, D. S. (2022). Maladaptive self-focused attention and default mode network connectivity: A transdiagnostic investigation across social anxiety and body dysmorphic disorders. Social Cognitive and Affective Neuroscience, 17(7), 645–654. 10.1093/scan/nsab130

Freyer, F., Roberts, J. A., Ritter, P., & Breakspear, M. (2012). A Canonical Model of Multistability and Scale-Invariance in Biological Systems. PLOS Computational Biology, 8(8), e1002634. 10.1371/journal.pcbi.1002634

Fribourg, M., Moreno, J. L., Holloway, T., Provasi, D., Baki, L., Mahajan, R., Park, G., Adney, S. K., Hatcher, C., Eltit, J. M., Ruta, J. D., Albizu, L., Li, Z., Umali, A., Shim, J., Fabiato, A., MacKerell, A. D., Brezina, V., Sealfon, S. C., … Logothetis, D. E. (2011). Decoding the Signaling of a GPCR Heteromeric Complex Reveals a Unifying Mechanism of Action of Antipsychotic Drugs. Cell, 147(5), 1011–1023. 10.1016/j.cell.2011.09.055

Gabrys, R. L., Tabri, N., Anisman, H., & Matheson, K. (2018). Cognitive Control and Flexibility in the Context of Stress and Depressive Symptoms: The Cognitive Control and Flexibility Questionnaire. Frontiers in Psychology, 9, 2219. 10.3389/fpsyg.2018.02219

Garcia, A. C. M., & Maia, L. de O. (2022). The therapeutic potential of psychedelic substances in Hospice and Palliative Care. Progress in Palliative Care, 30(1), 1–3. 10.1080/09699260.2022.2001140

Garcia, E. E., Smith, R. L., & Sanders-Bush, E. (2007). Role of G(q) protein in behavioral effects of the hallucinogenic drug 1-(2,5-dimethoxy-4-iodophenyl)-2-aminopropane. Neuropharmacology, 52(8), 1671–1677. 10.1016/j.neuropharm.2007.03.013

Garrido, M. I., Kilner, J. M., Stephan, K. E., & Friston, K. J. (2009). The mismatch negativity: A review of underlying mechanisms. Clinical Neurophysiology, 120(3), 453–463. 10.1016/j.clinph.2008.11.029

Girn, M., Roseman, L., Bernhardt, B., Smallwood, J., Carhart-Harris, R., & Nathan Spreng, R. (2022). Serotonergic psychedelic drugs LSD and psilocybin reduce the hierarchical differentiation of unimodal and transmodal cortex. NeuroImage, 256, 119220. 10.1016/j.neuroimage.2022.119220

Giulietti, M., Vivenzio, V., Piva, F., Principato, G., Bellantuono, C., & Nardi, B. (2014). How much do we know about the coupling of G-proteins to serotonin receptors? Molecular Brain, 7(1), 49. 10.1186/s13041-014-0049-y

Glennon, R. A., Titeler, M., & McKenney, J. D. (1984). Evidence for 5-HT2 involvement in the mechanism of action of hallucinogenic agents. Life Sciences, 35(25), 2505–2511. 10.1016/0024-3205(84)90436-3

González-Maeso, J., Weisstaub, N. V., Zhou, M., Chan, P., Ivic, L., Ang, R., Lira, A., Bradley-Moore, M., Ge, Y., Zhou, Q., Sealfon, S. C., & Gingrich, J. A. (2007). Hallucinogens recruit specific cortical 5-HT(2A) receptor-mediated signaling pathways to affect behavior. Neuron, 53(3), 439–452. 10.1016/j.neuron.2007.01.008

Goodwin, G. M., Aaronson, S. T., Alvarez, O., Arden, P. C., Baker, A., Bennett, J. C., Bird, C., Blom, R. E., Brennan, C., Brusch, D., Burke, L., Campbell-Coker, K., Carhart-Harris, R., Cattell, J., Daniel, A., DeBattista, C., Dunlop, B. W., Eisen, K., Feifel, D., … Malievskaia, E. (2022). Single-Dose Psilocybin for a Treatment-Resistant Episode of Major Depression. The New England Journal of Medicine, 387(18), 1637–1648. 10.1056/NEJMoa2206443

Goodwin, G. M., Malievskaia, E., Fonzo, G. A., & Nemeroff, C. B. (2024). Must Psilocybin Always “Assist Psychotherapy”? American Journal of Psychiatry, 181(1), 20–25. 10.1176/appi.ajp.20221043

Griffiths, R. R., Johnson, M. W., Richards, W. A., Richards, B. D., Jesse, R., MacLean, K. A., Barrett, F. S., Cosimano, M. P., & Klinedinst, M. A. (2018). Psilocybin-occasioned mystical-type experience in combination with meditation and other spiritual practices produces enduring positive changes in psychological functioning and in trait measures of prosocial attitudes and behaviors. Journal of Psychopharmacology (Oxford, England), 32(1), 49–69. 10.1177/0269881117731279

Grimm, O., Kraehenmann, R., Preller, K. H., Seifritz, E., & Vollenweider, F. X. (2018). Psilocybin modulates functional connectivity of the amygdala during emotional face discrimination. European Neuropsychopharmacology: The Journal of the European College of Neuropsychopharmacology, 28(6), 691–700. 10.1016/j.euroneuro.2018.03.016

Gurevich, E. V., Gainetdinov, R. R., & Gurevich, V. V. (2016). G protein-coupled receptor kinases as regulators of dopamine receptor functions. Pharmacological Research, 111, 1–16. 10.1016/j.phrs.2016.05.010

Hansen, J. Y., Shafiei, G., Markello, R. D., Smart, K., Cox, S. M. L., Nørgaard, M., Beliveau, V., Wu, Y., Gallezot, J.-D., Aumont, É., Servaes, S., Scala, S. G., DuBois, J. M., Wainstein, G., Bezgin, G., Funck, T., Schmitz, T. W., Spreng, R. N., Galovic, M., … Misic, B. (2022). Mapping neurotransmitter systems to the structural and functional organization of the human neocortex. Nature Neuroscience, 25(11), Article 11. 10.1038/s41593-022-01186-3

Hasler, F., Grimberg, U., Benz, M. A., Huber, T., & Vollenweider, F. X. (2004). Acute psychological and physiological effects of psilocybin in healthy humans: A double-blind, placebo-controlled dose-effect study. Psychopharmacology, 172(2), 145–156. 10.1007/s00213-003-1640-6

Haufe, S., DeGuzman, P., Henin, S., Arcaro, M., Honey, C. J., Hasson, U., & Parra, L. C. (2018). Elucidating relations between fMRI, ECoG, and EEG through a common natural stimulus. NeuroImage, 179, 79–91. 10.1016/j.neuroimage.2018.06.016

Heekeren, K., Daumann, J., Neukirch, A., Stock, C., Kawohl, W., Norra, C., Waberski, T. D., & Gouzoulis-Mayfrank, E. (2008). Mismatch negativity generation in the human 5HT2A agonist and NMDA antagonist model of psychosis. Psychopharmacology, 199(1), 77–88. 10.1007/s00213-008-1129-4

Holze, F., Ley, L., Müller, F., Becker, A. M., Straumann, I., Vizeli, P., Kuehne, S. S., Roder, M. A., Duthaler, U., Kolaczynska, K. E., Varghese, N., Eckert, A., & Liechti, M. E. (2022). Direct comparison of the acute effects of lysergic acid diethylamide and psilocybin in a double-blind placebo-controlled study in healthy subjects. Neuropsychopharmacology, 47(6), 1180–1187. 10.1038/s41386-022-01297-2

Honey, C. J., Kötter, R., Breakspear, M., & Sporns, O. (2007). Network structure of cerebral cortex shapes functional connectivity on multiple time scales. Proceedings of the National Academy of Sciences, 104(24), 10240–10245. 10.1073/pnas.0701519104

Hwang, J. W., Egorova, N., Yang, X. Q., Zhang, W. Y., Chen, J., Yang, X. Y., Hu, L. J., Sun, S., Tu, Y., & Kong, J. (2015). Subthreshold depression is associated with impaired resting-state functional connectivity of the cognitive control network. Translational Psychiatry, 5(11), e683. 10.1038/tp.2015.174

Janowsky, A., Eshleman, A. J., Johnson, R. A., Wolfrum, K. M., Hinrichs, D. J., Yang, J., Zabriskie, T. M., Smilkstein, M. J., & Riscoe, M. K. (2014). Mefloquine and Psychotomimetics Share Neurotransmitter Receptor and Transporter Interactions In Vitro. Psychopharmacology, 231(14), 2771–2783. 10.1007/s00213-014-3446-0

Kaelen, M., Roseman, L., Kahan, J., Santos-Ribeiro, A., Orban, C., Lorenz, R., Barrett, F. S., Bolstridge, M., Williams, T., Williams, L., Wall, M. B., Feilding, A., Muthukumaraswamy, S., Nutt, D. J., & Carhart-Harris, R. (2016). LSD modulates music-induced imagery via changes in parahippocampal connectivity. European Neuropsychopharmacology: The Journal of the European College of Neuropsychopharmacology, 26(7), 1099–1109. 10.1016/j.euroneuro.2016.03.018

Kometer, M., & Vollenweider, F. X. (2018). Serotonergic Hallucinogen-Induced Visual Perceptual Alterations. In A. L. Halberstadt, F. X. Vollenweider, & D. E. Nichols (Eds.), Behavioral Neurobiology of Psychedelic Drugs (pp. 257–282). Springer. 10.1007/7854_2016_461

Kozell, L. B., Eshleman, A. J., Swanson, T. L., Bloom, S. H., Wolfrum, K. M., Schmachtenberg, J. L., Olson, R. J., Janowsky, A., & Abbas, A. I. (2023). Pharmacologic Activity of Substituted Tryptamines at 5-Hydroxytryptamine (5-HT)2A Receptor (5-HT2AR), 5-HT2CR, 5-HT1AR, and Serotonin Transporter. The Journal of Pharmacology and Experimental Therapeutics, 385(1), 62–75. 10.1124/jpet.122.001454

Kraft, B., Bø, R., Jonassen, R., Heeren, A., Ulset, V. S., Stiles, T. C., & Landrø, N. I. (2023). The association between depression symptoms and reduced executive functioning is primarily linked by fatigue. Psychiatry Research Communications, 3(2), 100120. 10.1016/j.psycom.2023.100120

Kringelbach, M. L., Cruzat, J., Cabral, J., Knudsen, G. M., Carhart-Harris, R., Whybrow, P. C., Logothetis, N. K., & Deco, G. (2020). Dynamic coupling of whole-brain neuronal and neurotransmitter systems. Proceedings of the National Academy of Sciences, 117(17), 9566–9576. 10.1073/pnas.1921475117

Kringelbach, M. L., & Deco, G. (2020). Brain States and Transitions: Insights from Computational Neuroscience. Cell Reports, 32(10), 108128. 10.1016/j.celrep.2020.108128

Lawn, T., Dipasquale, O., Vamvakas, A., Tsougos, I., Mehta, M. A., & Howard, M. A. (2022). Differential contributions of serotonergic and dopaminergic functional connectivity to the phenomenology of LSD. Psychopharmacology, 239(6), 1797–1808. 10.1007/s00213-022-06117-5

Lebedev, A. V., Lövdén, M., Rosenthal, G., Feilding, A., Nutt, D. J., & Carhart-Harris, R. L. (2015). Finding the self by losing the self: Neural correlates of ego-dissolution under psilocybin. Human Brain Mapping, 36(8), 3137–3153. 10.1002/hbm.22833

Ley, L., Holze, F., Arikci, D., Becker, A. M., Straumann, I., Klaiber, A., Coviello, F., Dierbach, S., Thomann, J., Duthaler, U., Luethi, D., Varghese, N., Eckert, A., & Liechti, M. E. (2023). Comparative acute effects of mescaline, lysergic acid diethylamide, and psilocybin in a randomized, double-blind, placebo-controlled cross-over study in healthy participants. Neuropsychopharmacology, 1–9. 10.1038/s41386-023-01607-2

Linguiti, S., Vogel, J. W., Sydnor, V. J., Pines, A., Wellman, N., Basbaum, A., Eickhoff, C. R., Eickhoff, S. B., Edwards, R. R., Larsen, B., McKinstry-Wu, A., Scott, J. C., Roalf, D. R., Sharma, V., Strain, E. C., Corder, G., Dworkin, R. H., & Satterthwaite, T. D. (2023). Functional imaging studies of acute administration of classic psychedelics, ketamine, and MDMA: Methodological limitations and convergent results. Neuroscience & Biobehavioral Reviews, 154, 105421. 10.1016/j.neubiorev.2023.105421

Liu, Q., Gao, F., Wang, X., Xia, J., Yuan, G., Zheng, S., Zhong, M., & Zhu, X. (2023). Cognitive inflexibility is linked to abnormal frontoparietal-related activation and connectivity in obsessive-compulsive disorder. Human Brain Mapping, 44(16), 5460–5470. 10.1002/hbm.26457

López-Giménez, J. F., & González-Maeso, J. (2018). Hallucinogens and Serotonin 5-HT2A Receptor-Mediated Signaling Pathways. Current Topics in Behavioral Neurosciences, 36, 45–73. 10.1007/7854_2017_478

Luan, L., Eckernäs, E., Ashton, M., Rosas, F., Uthaug, M., Bartha, A., Jagger, S., Gascon-Perai, K., Gomes, L., Nutt, D., Erritzoe, D., Carhart-Harris, R., & Timmermann, C. (2023). Psychological and physiological effects of extended DMT. PsyArXiv. 10.31234/osf.io/vg4dp

Luppi, A. I., Carhart-Harris, R. L., Roseman, L., Pappas, I., Menon, D. K., & Stamatakis, E. A. (2021). LSD alters dynamic integration and segregation in the human brain. NeuroImage, 227, 117653. 10.1016/j.neuroimage.2020.117653

Lyon, A. (2023). Attention. In A. Lyon (Ed.), Psychedelic Experience: Revealing the Mind (p. 0). Oxford University Press. 10.1093/oso/9780198843757.003.0005

Madsen, M. K., Stenbæk, D. S., Arvidsson, A., Armand, S., Marstrand-Joergensen, M. R., Johansen, S. S., Linnet, K., Ozenne, B., Knudsen, G. M., & Fisher, P. M. (2021). Psilocybin-induced changes in brain network integrity and segregation correlate with plasma psilocin level and psychedelic experience. European Neuropsychopharmacology, 50, 121–132. 10.1016/j.euroneuro.2021.06.001

Marek, S., & Dosenbach, N. U. F. (2018). The frontoparietal network: Function, electrophysiology, and importance of individual precision mapping. Dialogues in Clinical Neuroscience, 20(2), 133–140.

Martín-Signes, M., Cano-Melle, C., & Chica, A. B. (2021). Fronto-parietal networks underlie the interaction between executive control and conscious perception: Evidence from TMS and DWI. Cortex, 134, 1–15. 10.1016/j.cortex.2020.09.027

Mertens, L. J., Wall, M. B., Roseman, L., Demetriou, L., Nutt, D. J., & Carhart-Harris, R. L. (2020). Therapeutic mechanisms of psilocybin: Changes in amygdala and prefrontal functional connectivity during emotional processing after psilocybin for treatment-resistant depression. Journal of Psychopharmacology (Oxford, England), 34(2), 167–180. 10.1177/0269881119895520

Molnar-Szakacs, I., & Uddin, L. (2013). Self-Processing and the Default Mode Network: Interactions with the Mirror Neuron System. Frontiers in Human Neuroscience, 7. https://www.frontiersin.org/article/10.3389/fnhum.2013.00571

Moran, J. M., Kelley, W. M., & Heatherton, T. F. (2013). What Can the Organization of the Brain’s Default Mode Network Tell us About Self-Knowledge? Frontiers in Human Neuroscience, 7, 391. 10.3389/fnhum.2013.00391

Moreton, S. G., Szalla, L., Menzies, R. E., & Arena, A. F. (2020). Embedding existential psychology within psychedelic science: Reduced death anxiety as a mediator of the therapeutic effects of psychedelics. Psychopharmacology, 237(1), 21–32. 10.1007/s00213-019-05391-0

Müller, F., Dolder, P. C., Schmidt, A., Liechti, M. E., & Borgwardt, S. (2018). Altered network hub connectivity after acute LSD administration. NeuroImage. Clinical, 18, 694–701. 10.1016/j.nicl.2018.03.005

Müller, F., Liechti, M. E., Lang, U. E., & Borgwardt, S. (2018). Chapter 6 - Advances and challenges in neuroimaging studies on the effects of serotonergic hallucinogens: Contributions of the resting brain. In T. Calvey (Ed.), Progress in Brain Research (Vol. 242, pp. 159–177). Elsevier. 10.1016/bs.pbr.2018.08.004

Muttoni, S., Ardissino, M., & John, C. (2019). Classical psychedelics for the treatment of depression and anxiety: A systematic review. Journal of Affective Disorders, 258, 11–24. 10.1016/j.jad.2019.07.076

Nichols, D. E. (2016). Psychedelics. Pharmacological Reviews, 68(2), 264–355. 10.1124/pr.115.011478

Nour, M. M., Evans, L., Nutt, D., & Carhart-Harris, R. L. (2016). Ego-Dissolution and Psychedelics: Validation of the Ego-Dissolution Inventory (EDI). Frontiers in Human Neuroscience, 10. https://www.frontiersin.org/article/10.3389/fnhum.2016.00269

Nutt, D., Erritzoe, D., & Carhart-Harris, R. (2020). Psychedelic Psychiatry’s Brave New World. Cell, 181(1), 24–28. 10.1016/j.cell.2020.03.020

Pahnke, W. N. (1969). The Psychedelic Mystical Experience in the Human Encounter with Death*. Harvard Theological Review, 62(1), 1–21. 10.1017/S0017816000027577

Palhano-Fontes, F., Andrade, K. C., Tofoli, L. F., Santos, A. C., Crippa, J. A. S., Hallak, J. E. C., Ribeiro, S., & Araujo, D. B. de. (2015). The Psychedelic State Induced by Ayahuasca Modulates the Activity and Connectivity of the Default Mode Network. PLOS ONE, 10(2), e0118143. 10.1371/journal.pone.0118143

Pasquini, L., Palhano-Fontes, F., & Araujo, D. B. (2020). Subacute effects of the psychedelic ayahuasca on the salience and default mode networks. Journal of Psychopharmacology, 34(6), 623–635. 10.1177/0269881120909409

Pierce, P. A., & Peroutka, S. J. (1989). Hallucinogenic drug interactions with neurotransmitter receptor binding sites in human cortex. Psychopharmacology, 97(1), 118–122. 10.1007/BF00443425

Pokorny, T., Preller, K. H., Kraehenmann, R., & Vollenweider, F. X. (2016). Modulatory effect of the 5-HT1A agonist buspirone and the mixed non-hallucinogenic 5-HT1A/2A agonist ergotamine on psilocybin-induced psychedelic experience. European Neuropsychopharmacology, 26(4), 756–766. 10.1016/j.euroneuro.2016.01.005

Pottie, E., Cannaert, A., & Stove, C. P. (2020). In vitro structure-activity relationship determination of 30 psychedelic new psychoactive substances by means of β-arrestin 2 recruitment to the serotonin 2A receptor. Archives of Toxicology, 94(10), 3449–3460. 10.1007/s00204-020-02836-w

Pottie, E., Dedecker, P., & Stove, C. P. (2020). Identification of psychedelic new psychoactive substances (NPS) showing biased agonism at the 5-HT2AR through simultaneous use of β-arrestin 2 and miniGαq bioassays. Biochemical Pharmacology, 182, 114251. 10.1016/j.bcp.2020.114251

Preller, K. H., Herdener, M., Pokorny, T., Planzer, A., Kraehenmann, R., Stämpfli, P., Liechti, M. E., Seifritz, E., & Vollenweider, F. X. (2017). The Fabric of Meaning and Subjective Effects in LSD-Induced States Depend on Serotonin 2A Receptor Activation. Current Biology: CB, 27(3), 451–457. 10.1016/j.cub.2016.12.030

Qiu, T. (Tim), & Minda, J. P. (2021). Recreational Psychedelic Users Frequently Encounter Complete Mystical Experiences: Trip Content and Implications for Wellbeing. PsyArXiv. 10.31234/osf.io/xrbzs

Quednow, B. B., Kometer, M., Geyer, M. A., & Vollenweider, F. X. (2012). Psilocybin-Induced Deficits in Automatic and Controlled Inhibition are Attenuated by Ketanserin in Healthy Human Volunteers. Neuropsychopharmacology, 37(3), Article 3. 10.1038/npp.2011.228

Rai, S., Griffiths, K. R., Breukelaar, I. A., Barreiros, A. R., Chen, W., Boyce, P., Hazell, P., Foster, S. L., Malhi, G. S., Harris, A. W. F., & Korgaonkar, M. S. (2021). Default-mode and fronto-parietal network connectivity during rest distinguishes asymptomatic patients with bipolar disorder and major depressive disorder. Translational Psychiatry, 11(1), 1–8. 10.1038/s41398-021-01660-9

Raison, C. L., Sanacora, G., Woolley, J., Heinzerling, K., Dunlop, B. W., Brown, R. T., Kakar, R., Hassman, M., Trivedi, R. P., Robison, R., Gukasyan, N., Nayak, S. M., Hu, X., O’Donnell, K. C., Kelmendi, B., Sloshower, J., Penn, A. D., Bradley, E., Kelly, D. F., … Griffiths, R. R. (2023). Single-Dose Psilocybin Treatment for Major Depressive Disorder: A Randomized Clinical Trial. JAMA, 330(9), 843–853. 10.1001/jama.2023.14530

Rickli, A., Moning, O. D., Hoener, M. C., & Liechti, M. E. (2016). Receptor interaction profiles of novel psychoactive tryptamines compared with classic hallucinogens. European Neuropsychopharmacology: The Journal of the European College of Neuropsychopharmacology, 26(8), 1327–1337. 10.1016/j.euroneuro.2016.05.001

Rodriguiz, R. M., Nadkarni, V., Means, C. R., Pogorelov, V. M., Chiu, Y.-T., Roth, B. L., & Wetsel, W. C. (2021). LSD-stimulated behaviors in mice require β-arrestin 2 but not β-arrestin 1. Scientific Reports, 11(1), Article 1. 10.1038/s41598-021-96736-3

Roseman, L., Demetriou, L., Wall, M. B., Nutt, D. J., & Carhart-Harris, R. L. (2018). Increased amygdala responses to emotional faces after psilocybin for treatment-resistant depression. Neuropharmacology, 142, 263–269. 10.1016/j.neuropharm.2017.12.041

Roseman, L., Leech, R., Feilding, A., Nutt, D. J., & Carhart-Harris, R. L. (2014). The effects of psilocybin and MDMA on between-network resting state functional connectivity in healthy volunteers. Frontiers in Human Neuroscience, 8. https://www.frontiersin.org/article/10.3389/fnhum.2014.00204

Roseman, L., Nutt, D. J., & Carhart-Harris, R. L. (2018). Quality of Acute Psychedelic Experience Predicts Therapeutic Efficacy of Psilocybin for Treatment-Resistant Depression. Frontiers in Pharmacology, 8. https://www.frontiersin.org/articles/10.3389/fphar.2017.00974

Sanz, C., Zamberlan, F., Erowid, E., Erowid, F., & Tagliazucchi, E. (2018). The Experience Elicited by Hallucinogens Presents the Highest Similarity to Dreaming within a Large Database of Psychoactive Substance Reports. Frontiers in Neuroscience, 12. 10.3389/fnins.2018.00007

Schmitz, G. P., Jain, M. K., Slocum, S. T., & Roth, B. L. (2022). 5-HT2A SNPs Alter the Pharmacological Signaling of Potentially Therapeutic Psychedelics. ACS Chemical Neuroscience, 13(16), 2386–2398. 10.1021/acschemneuro.1c00815

Schrantee, A., Ferguson, B., Stoffers, D., Booij, J., Rombouts, S., & Reneman, L. (2016). Effects of dexamphetamine-induced dopamine release on resting-state network connectivity in recreational amphetamine users and healthy controls. Brain Imaging and Behavior, 10(2), 548–558. 10.1007/s11682-015-9419-z

Shah, U., Pincas, H., Sealfon, S. C., & González-Maeso, J. (2020). Chapter 11—Structure and function of serotonin GPCR heteromers. In C. P. Müller & K. A. Cunningham (Eds.), Handbook of Behavioral Neuroscience (Vol. 31, pp. 217–238). Elsevier. 10.1016/B978-0-444-64125-0.00011-6

Siegel, J. S., Subramanian, S., Perry, D., Kay, B. P., Gordon, E. M., Laumann, T. O., Reneau, T. R., Metcalf, N. V., Chacko, R. V., Gratton, C., Horan, C., Krimmel, S. R., Shimony, J. S., Schweiger, J. A., Wong, D. F., Bender, D. A., Scheidter, K. M., Whiting, F. I., Padawer-Curry, J. A., … Dosenbach, N. U. F. (2024). Psilocybin desynchronizes the human brain. Nature, 632(8023), 131–138. 10.1038/s41586-024-07624-5

Smigielski, L., Scheidegger, M., Kometer, M., & Vollenweider, F. X. (2019). Psilocybin-assisted mindfulness training modulates self-consciousness and brain default mode network connectivity with lasting effects. NeuroImage, 196, 207–215. 10.1016/j.neuroimage.2019.04.009

Studerus, E., Gamma, A., & Vollenweider, F. X. (2010). Psychometric Evaluation of the Altered States of Consciousness Rating Scale (OAV). PLOS ONE, 5(8), e12412. 10.1371/journal.pone.0012412

Sykes, D. A., Stoddart, L. A., Kilpatrick, L. E., & Hill, S. J. (2019). Binding kinetics of ligands acting at GPCRs. Molecular and Cellular Endocrinology, 485, 9–19. 10.1016/j.mce.2019.01.018

Tabaac, B. J., Shinozuka, K., Arenas, A., Beutler, B. D., Cherian, K., Evans, V. D., Fasano, C., & Muir, O. S. (2024). Psychedelic Therapy: A Primer for Primary Care Clinicians-Psilocybin. American Journal of Therapeutics, 31(2), e121–e132. 10.1097/MJT.0000000000001724

Tagliazucchi, E., Carhart-Harris, R., Leech, R., Nutt, D., & Chialvo, D. R. (2014). Enhanced repertoire of brain dynamical states during the psychedelic experience. Human Brain Mapping, 35(11), 5442–5456. 10.1002/hbm.22562

Tagliazucchi, E., Roseman, L., Kaelen, M., Orban, C., Muthukumaraswamy, S. D., Murphy, K., Laufs, H., Leech, R., McGonigle, J., Crossley, N., Bullmore, E., Williams, T., Bolstridge, M., Feilding, A., Nutt, D. J., & Carhart-Harris, R. (2016). Increased Global Functional Connectivity Correlates with LSD-Induced Ego Dissolution. Current Biology: CB, 26(8), 1043–1050. 10.1016/j.cub.2016.02.010

Thomas Yeo, B. T., Krienen, F. M., Sepulcre, J., Sabuncu, M. R., Lashkari, D., Hollinshead, M., Roffman, J. L., Smoller, J. W., Zöllei, L., Polimeni, J. R., Fischl, B., Liu, H., & Buckner, R. L. (2011). The organization of the human cerebral cortex estimated by intrinsic functional connectivity. Journal of Neurophysiology, 106(3), 1125–1165. 10.1152/jn.00338.2011

Timmermann, C., Roseman, L., Haridas, S., Rosas, F. E., Luan, L., Kettner, H., Martell, J., Erritzoe, D., Tagliazucchi, E., Pallavicini, C., Girn, M., Alamia, A., Leech, R., Nutt, D. J., & Carhart-Harris, R. L. (2023). Human brain effects of DMT assessed via EEG-fMRI. Proceedings of the National Academy of Sciences of the United States of America, 120(13), e2218949120. 10.1073/pnas.2218949120

Timmermann, C., Spriggs, M. J., Kaelen, M., Leech, R., Nutt, D. J., Moran, R. J., Carhart-Harris, R. L., & Muthukumaraswamy, S. D. (2018). LSD modulates effective connectivity and neural adaptation mechanisms in an auditory oddball paradigm. Neuropharmacology, 142, 251–262. 10.1016/j.neuropharm.2017.10.039

Tófoli, L. F., & de Araujo, D. B. (2016). Treating Addiction: Perspectives from EEG and Imaging Studies on Psychedelics. International Review of Neurobiology, 129, 157–185. 10.1016/bs.irn.2016.06.005

Tupper, K. W., Wood, E., Yensen, R., & Johnson, M. W. (2015). Psychedelic medicine: A re-emerging therapeutic paradigm. CMAJ : Canadian Medical Association Journal, 187(14), 1054–1059. 10.1503/cmaj.141124

Tuteja, N. (2009). Signaling through G protein coupled receptors. Plant Signaling & Behavior, 4(10), 942–947.

van Buuren, M., Gladwin, T. E., Zandbelt, B. B., Kahn, R. S., & Vink, M. (2010). Reduced functional coupling in the default-mode network during self-referential processing. Human Brain Mapping, 31(8), 1117–1127. 10.1002/hbm.20920

Van Essen, D. C., Smith, S. M., Barch, D. M., Behrens, T. E. J., Yacoub, E., & Ugurbil, K. (2013). The WU-Minn Human Connectome Project: An overview. NeuroImage, 80, 62–79. 10.1016/j.neuroimage.2013.05.041

Vogt, S. B., Ley, L., Erne, L., Straumann, I., Becker, A. M., Klaiber, A., Holze, F., Vandersmissen, A., Mueller, L., Duthaler, U., Rudin, D., Luethi, D., Varghese, N., Eckert, A., & Liechti, M. E. (2023). Acute effects of intravenous DMT in a randomized placebo-controlled study in healthy participants. Translational Psychiatry, 13(1), 172. 10.1038/s41398-023-02477-4

Vollenweider, F. X., Csomor, P. A., Knappe, B., Geyer, M. A., & Quednow, B. B. (2007). The effects of the preferential 5-HT2A agonist psilocybin on prepulse inhibition of startle in healthy human volunteers depend on interstimulus interval. Neuropsychopharmacology: Official Publication of the American College of Neuropsychopharmacology, 32(9), 1876–1887. 10.1038/sj.npp.1301324

Vollenweider, F. X., Vollenweider-Scherpenhuyzen, M. F., Bäbler, A., Vogel, H., & Hell, D. (1998). Psilocybin induces schizophrenia-like psychosis in humans via a serotonin-2 agonist action. Neuroreport, 9(17), 3897–3902. 10.1097/00001756-199812010-00024

Von Rotz, R., Schindowski, E. M., Jungwirth, J., Schuldt, A., Rieser, N. M., Zahoranszky, K., Seifritz, E., Nowak, A., Nowak, P., Jäncke, L., Preller, K. H., & Vollenweider, F. X. (2023). Single-dose psilocybin-assisted therapy in major depressive disorder: A placebo-controlled, double-blind, randomised clinical trial. eClinicalMedicine, 56, 101809. 10.1016/j.eclinm.2022.101809

Vossel, S., Geng, J. J., & Fink, G. R. (2014). Dorsal and Ventral Attention Systems. The Neuroscientist, 20(2), 150–159. 10.1177/1073858413494269

Wall, M. B., Lam, C., Ertl, N., Kaelen, M., Roseman, L., Nutt, D. J., & Carhart-Harris, R. L. (2022). Increased low-frequency brain responses to music after psilocybin therapy for depression (p. 2022.02.13.480302). bioRxiv. 10.1101/2022.02.13.480302

Wallach, J., Cao, A. B., Calkins, M. M., Heim, A. J., Lanham, J. K., Bonniwell, E. M., Hennessey, J. J., Bock, H. A., Anderson, E. I., Sherwood, A. M., Morris, H., Klein, R. de, Klein, A. K., Cuccurazzu, B., Gamrat, J., Fannana, T., Zauhar, R., Halberstadt, A. L., & McCorvy, J. D. (2023). Identification of 5-HT2A Receptor Signaling Pathways Responsible for Psychedelic Potential (p. 2023.07.29.551106). bioRxiv. 10.1101/2023.07.29.551106

Watts, V. J., Lawler, C. P., Fox, D. R., Neve, K. A., Nichols, D. E., & Mailman, R. B. (1995). LSD and structural analogs: Pharmacological evaluation at D1 dopamine receptors. Psychopharmacology, 118(4), 401–409. 10.1007/BF02245940

Weber, A. M., Soreni, N., & Noseworthy, M. D. (2014). A preliminary study on the effects of acute ethanol ingestion on default mode network and temporal fractal properties of the brain. Magma (New York, N.Y.), 27(4), 291–301. 10.1007/s10334-013-0420-5

Xia, M., Wang, J., & He, Y. (2013). BrainNet Viewer: A Network Visualization Tool for Human Brain Connectomics. PLOS ONE, 8(7), e68910. 10.1371/journal.pone.0068910

Yaden, D. B., Johnson, M. W., Griffiths, R. R., Doss, M. K., Garcia-Romeu, A., Nayak, S., Gukasyan, N., Mathur, B. N., & Barrett, F. S. (2021). Psychedelics and consciousness: Distinctions, demarcations, and opportunities. International Journal of Neuropsychopharmacology, 24(8), 615–623. 10.1093/ijnp/pyab026

Zamberlan, F., Sanz, C., Martínez Vivot, R., Pallavicini, C., Erowid, F., Erowid, E., & Tagliazucchi, E. (2018). The Varieties of the Psychedelic Experience: A Preliminary Study of the Association Between the Reported Subjective Effects and the Binding Affinity Profiles of Substituted Phenethylamines and Tryptamines. Frontiers in Integrative Neuroscience, 12, 54. 10.3389/fnint.2018.00054

Zhang, R., & Volkow, N. D. (2019). Brain default-mode network dysfunction in addiction. NeuroImage, 200, 313–331. 10.1016/j.neuroimage.2019.06.036

Zhou, H.-X., Chen, X., Shen, Y.-Q., Li, L., Chen, N.-X., Zhu, Z.-C., Castellanos, F. X., & Yan, C.-G. (2020). Rumination and the default mode network: Meta-analysis of brain imaging studies and implications for depression. NeuroImage, 206, 116287. 10.1016/j.neuroimage.2019.116287

