## Supplementary Materials for "Synergistic, Multi-level Understanding of Psychedelics: Three Systematic Reviews and Meta-analyses of Their Pharmacology, Neuroimaging and Phenomenology"

### Section S1. Literature Search

We performed systematic reviews and meta-analyses of three different levels of psychedelic effects: phenomenology, neuroimaging, and pharmacology. Each review and meta-analysis examined three classical psychedelics: DMT, LSD, and psilocybin. The phenomenology meta-analyses separately assessed two versions (5-dimensional and 11-dimensional) of the Altered States of Consciousness (ASC) scale. We also qualitatively reviewed studies that directly compared the phenomenology of all three classical psychedelics. For the functional neuroimaging (fMRI) literature, we conducted a GingerALE analysis on the BOLD activation data and ran an in-house algorithm to analyse the functional connectivity (FC) data. (Note that we did not ultimately report the results of the BOLD activation meta-analysis because we discovered that there was too much heterogeneity in the experimental procedures.) We also performed a qualitative review of the studies on entropy under psychedelics. Finally, we carried out two meta-analyses on the pharmacology literature: one on binding affinity data, and another on functional activity data, as measured by assays on GPCR-mediated signaling. Therefore, in total, we conducted six quantitative meta-analyses and two qualitative reviews.

We conducted all of our literature searches in accordance with the most recent standards of PRISMA (Preferred Reporting Items for Systematic Reviews and Meta-Analyses), a set of guidelines for performing meta-analyses (Page et al., 2021). Our **PRISMA 2020 Checklist** is included as a supplementary document. For each literature search, one reviewer screened each record and each report received, identifying whether reports met any of the exclusion criteria by reading their title and sometimes their abstract. One reviewer also collected data from each report (specific methods described below).

##### *Section S1.1. Phenomenology*

We firstly performed a quantitative meta-analysis of studies that measured the subjective effects of DMT, LSD, and psilocybin using the 5-dimensional and 11-dimensional versions of the Altered States of Consciousness (ASC) scale. We focused on these scales because they are the most common questionnaires in the neuroimaging literature on psychedelics. We also qualitatively reviewed the studies that directly compared the phenomenology of DMT/ayahuasca, LSD, and psilocybin (*Section S3*).

We conducted our literature search of the ASC data in August 2023. We examined four databases: SCOPUS, Web of Science, the Altered States Database, and Google Scholar. There were two studies that we independently discovered from citations in the included papers; these studies did not appear on any of the databases. The Altered States Database (alteredstatesdb.org) is a publicly available website that displays data from several different questionnaires about a variety of altered states of consciousness (Prugger et al., 2022; Schmidt & Berkemeyer, 2018). It is not exhaustive, however. Search terms followed the format "<drug name> AND <scale name>", where "drug name" was "LSD", "psilocybin", "DMT", or "N,N-dimethyltryptamine" and "scale name" was "ASC", "5D-ASC", or "11D-ASC". We excluded papers based on the following criteria:

1. Work that did not report any original experiments or analyses, such as literature reviews, commentaries and editorials.

2. Studies that did not examine one or more of three classical psychedelics: DMT, LSD, psilocybin. Note that we did not include studies on ayahuasca for this meta-analysis.
3. Studies that did not use the ASC scale.
4. Studies that did not report original ASC data.
5. Studies that were conducted in a clinical population of participants, such as depressed participants. Only studies on healthy human participants were included.
6. Studies in which the participants were meditators or learning meditation, as these participants may experience psychedelics in significantly different ways than non-meditators do.
7. Studies on extended DMT, a recently-developed experimental technique that prolongs the duration of the DMT trip (Luan et al., 2023).
8. Studies on microdosing, since the subjective effects of microdosing are often indistinguishable from those of placebo (Bershad et al., 2020).
9. Studies that reported data in unclear units.
10. Original studies that administered the ASC questionnaire to participants but did not publish any ASC data. This includes studies that measured and reported correlations between ASC results and other metrics, but did not actually report the ASC scores themselves.
11. Studies that administered LSD or psilocybin intravenously. Most phenomenological studies on LSD and psilocybin administer the drugs orally, and IV administration yields very different pharmacokinetics and hence subjective effects.

A PRISMA flowchart of the literature search is shown in **Figure S1a.** Our search yielded ***n* = 44 studies in total** (Becker et al., 2023; Bernasconi et al., 2014; Carbonaro et al., 2018; Carter, Burr, et al., 2005; Carter, Pettigrew, et al., 2005; Carter et al., 2007; Daumann et al., 2008; Duerler et al., 2020; Family et al., 2022; Gouzoulis-Mayfrank et al., 2005; Hasler et al., 2004; Holze et al., 2020, 2021, 2022; Kometer et al., 2012; Kraehenmann, Preller, et al., 2015; Kraehenmann et al., 2017; Lewis et al., 2017; Ley et al., 2023; Madsen et al., 2021; Mallaroni et al., 2023; Mason et al., 2020; McAlpine et al., 2023; Müller, Lenz, Dolder, Lang, et al., 2017; Ort et al., 2023; Pallavicini et al., 2021; Pokorny et al., 2016, 2017, 2020; Preller et al., 2016, 2017, 2020; Quednow et al., 2012; Schmid et al., 2015; A. Schmidt et al., 2012, 2018; Smigielski et al., 2020; Straumann et al., 2023; Timmermann et al., 2023; Umbricht et al., 2002; Vogt et al., 2023; Vollenweider et al., 2007; Wießner et al., 2023; Wittmann et al., 2007). These included ***n* = 21 studies on the 5D-ASC** and ***n* = 31 studies on the 11D-ASC**; there were *n* = 11 studies that administered both versions of the questionnaire. The 5D-ASC literature included *n* = 3 studies on DMT, *n* = 9 studies on LSD, and *n* = 12 studies on psilocybin. The 11D-ASC literature contained *n* = 3 studies on DMT, *n* = 12 studies on LSD, and *n* = 17 studies on psilocybin. Two studies gave both LSD and psilocybin to participants (Holze et al., 2022; Ley et al., 2023). Data from a total of 1,037 participants were examined in the meta-analyses (DMT: 111 participants; LSD: 394 participants; psilocybin: 592 participants). Two of the 5D-ASC studies on DMT measured subjective effects for only three of the dimensions, so the remaining two dimensions (auditory alterations and reduction of vigilance) were only examined by a single study (Vogt et al.,

2023). Additionally, two of the 11D-ASC studies on DMT did not report standard errors (Pallavicini et al., 2021; Timmermann et al., 2023). In our meta-analyses, we pooled ASC scores by weighting each one by its inverse variance (squared standard error), so the standard errors were essential. We reached out to the authors for the standard errors but did not hear back. Since there was only one 11D-ASC study on DMT that did publish standard errors, we decided not to perform a meta-analysis of the 11D-ASC data on DMT.

Many studies reported numerical 5D- and 11D-ASC data. However, others merely displayed plots of the data. We chose not to email the authors for the numerical data, since it could be very time-consuming to wait for responses. Instead, we translated graphical data into numerical data by counting the number of pixels that corresponded to ASC scores. For instance, in a bar plot of ASC scores, the length of each bar in pixels would indicate the ASC score. By dividing this length by the length of the relevant axis, we obtained the numerical ASC data. While this technique is imprecise, we found that it generally succeeded when we tested it on studies for which numerical data existed.

In order to determine the neural correlates of the subjective effects of each psychedelic, which we did for the purpose of comparing phenomenology to neuroimaging and pharmacology, we conducted a separate literature search in SCOPUS and the Altered States Database. We identified three new studies from this search (Mueller et al., 2018; Northoff et al., 2005; Scheidegger et al., 2016).

##### *Section S1.2. Neuroimaging*

We conducted our literature search of the fMRI studies in May 2023. We searched four large databases: SCOPUS, Web of Science, psycINFO, and bioRxiv. (Note that, even though we searched bioRxiv, which is a pre-print server, we did not ultimately include any pre-prints in the meta-analysis; we only included pre-prints that were later published by the time that we performed our search.) Our search terms followed the structure: "<drug name> AND <neuroimaging term>", where <drug name> was "psychedelics", "LSD", "psilocybin", "DMT", or "ayahuasca", and <neuroimaging term> was "fMRI", "MRI", "functional magnetic resonance imaging", or "neuroimaging". We then excluded any reports that met the following criteria:

1. Works that did not report any original research, such as literature reviews, commentaries, and editorials.
2. Studies in which the experimental drug was not a classical psychedelic. We define a "classical psychedelic" as any drug whose primary mechanism of action is mediated by the $5\text{-HT}_{2A}$ receptor. This criterion excluded studies that investigated MDMA, cannabis/THC, and/or ketamine. We included studies on the following classical psychedelics: ayahuasca, DMT, LSD, and psilocybin.
3. Studies that did not collect any new fMRI data or run a novel analysis on existing fMRI data. This excluded studies that measured brain activity under psychedelics using MEG, EEG, PET, structural MRI, or other techniques, as well as studies that gathered only behavioural or subjective data. Additionally, we excluded studies that conducted new measurements of correlations between subjective data and existing neuroimaging data, but did not analyse the neuroimaging data itself in an original way (Speth et al., 2016).

4. Studies that were conducted on non-human animals or on *in vitro* cell populations.
5. Studies whose results were not specific to psychedelics, including studies examining the neural correlates of hallucinogen, poly-drug, or general drug use/addiction (e.g. Butcher et al. (2021)).
6. Studies that investigated disorders caused by psychedelic use, such as palinopsia (Kawasaki & Purvin, 1996), cortical blindness (Bernhard & Ulrich, 2009), persistent visual hallucinations (Iaria et al., 2009), and more. Psychedelics rarely induce long-term neurological damage (Studerus et al., 2011), so we did not incorporate the neural correlates of these disorders into our meta-analysis.
7. Studies that were performed on participants from a clinical population, such as patients suffering from depression. Psychedelics may affect brain activity in depressed patients very differently than in healthy participants (Carhart-Harris et al., 2017).
8. Studies that primarily simulated brain activity under the influence of psychedelics, rather than measuring or analysing empirical neuroimaging data on psychedelics (Burt et al., 2021; Deco, Cruzat, et al., 2018; Kringelbach et al., 2020).
9. Studies that measured post-acute effects of psychedelics, i.e. brain activity after the psychedelic trip had already ended.
10. Studies on microdosing. Since the subjective effects of microdoses are often imperceptible, there is good reason to think that brain activity in response to a microdose would be significantly different than brain activity on a macrodose.

The flowchart in **Figure S1b** summarises our literature search, which led us to identify ***n*** **= 58 studies in total**. We further divided the studies into three sections: studies on BOLD activation, on connectivity, and on entropy. We included ***n*** **= 17 studies on BOLD activation** (Barrett, Doss, et al., 2020; Barrett et al., 2018; Carhart-Harris, Erritzoe, et al., 2012; Carhart-Harris, Leech, et al., 2012; Daumann et al., 2008, 2010; de Araujo et al., 2011; Duerler et al., 2020, 2021; Kaelen et al., 2017; Kraehenmann, Preller, et al., 2015; Müller, Lenz, Dolder, Harder, et al., 2017; Palhano-Fontes et al., 2015; Preller et al., 2016, 2017; Preller, Schilbach, et al., 2018; Schmidt et al., 2018), ***n*** **= 34 studies on connectivity** (Avram et al., 2022; Barrett, Doss, et al., 2020; Barrett, Krimmel, et al., 2020; Bedford et al., 2023; Carhart-Harris et al., 2013, 2016; Dai et al., 2023; de Araujo et al., 2011; Delli Pizzi et al., 2023; Gaddis et al., 2022; Girn et al., 2022; Grimm et al., 2018; Jobst et al., 2021; Kaelen et al., 2016; Kraehenmann, Schmidt, et al., 2015; Lebedev et al., 2015; Lord et al., 2019; Luppi et al., 2021; Madsen et al., 2021; Mason et al., 2020, 2021; Müller et al., 2018; Müller, Lenz, Dolder, Lang, et al., 2017; Olsen et al., 2022; Palhano-Fontes et al., 2015; Petri et al., 2014; Preller, Burt, et al., 2018; Preller et al., 2019, 2020; Roseman et al., 2014, 2016; Tagliazucchi et al., 2014, 2016; Timmermann et al., 2023), **and** ***n*** **= 12 studies on entropy/criticality** (Atasoy et al., 2017; Carhart-Harris et al., 2014; Lebedev et al., 2015, 2016; Lord et al., 2019; Luppi et al., 2023; Ruffini et al., 2023; Singleton et al., 2022; Tagliazucchi et al., 2014; Varley et al., 2020; Viol et al., 2017, 2019, 2023). There was one study that did not fall neatly into any of the three categories (Viol et al., 2023).

We then sought the relevant data for each section. We did not require any data on entropy as we performed **a qualitative review of all** ***n*** **= 12 studies on entropy/criticality**. To conduct a meta-analysis on studies of BOLD activation, we used the stereotaxic coordinates in Montreal Neurological Institute (MNI) space of the brain regions that displayed significant changes in BOLD activation under psychedelics. We conducted **a quantitative meta-analysis on all *n* = 17 studies of BOLD activation**. For the FC meta-analysis, we obtained either the coordinates or the spatial maps (in MNI space, at $2mm^3$ resolution) of the brain areas that exhibited significant changes in connectivity (see **Table S5** for a comprehensive account of how we incorporated the data from each study). Our FC meta-analysis algorithm could only accommodate data from studies on pairwise FC, i.e. correlations between pairs of regions. However, there were 11 studies on connectivity that did not measure pairwise FC; instead, they examined global, structural indices of connectivity, such as modularity and GBC, or effective connectivity. We also excluded eight more studies of pairwise FC from the quantitative meta-analysis because they were re-analyses of datasets that we had already included (see *Section S2.2.2*). Finally, we were unable to obtain data from three studies on pairwise FC, which were otherwise eligible for the quantitative meta-analysis. Thus, we conducted a **quantitative meta-analysis on *n* = 12 studies of FC** (Barrett, Krimmel, et al., 2020; Bedford et al., 2023; Dai et al., 2023; de Araujo et al., 2011; Gaddis et al., 2022; Grimm et al., 2018; Kaelen et al., 2016; Madsen et al., 2021; Mason et al., 2020; Palhano-Fontes et al., 2015; Roseman et al., 2014; Timmermann et al., 2023), and we performed a **qualitative review of *n* = 22 studies on connectivity** (Avram et al., 2022; Barrett, Doss, et al., 2020; Carhart-Harris et al., 2013, 2016; Delli Pizzi et al., 2023; Girn et al., 2022; Jobst et al., 2021; Kraehenmann, Schmidt, et al., 2015; Lebedev et al., 2015; Lord et al., 2019; Luppi et al., 2021; Mason et al., 2021; Müller et al., 2018; Müller, Lenz, Dolder, Lang, et al., 2017; Olsen et al., 2022; Petri et al., 2014; Preller, Burt, et al., 2018; Preller et al., 2019, 2020; Roseman et al., 2016; Tagliazucchi et al., 2014, 2016).

Out of the 58 fMRI studies, only 22 produced primary datasets, so over half of the studies are secondary analyses (**Figure S2b**). There are disproportionately more secondary analyses of the primary datasets in Carhart-Harris, Erritzoe, et al. (2012) and Carhart-Harris et al. (2016) (10 of the former and 15 of the latter, among the studies that we included from our literature search) than of any other datasets. Across all primary datasets that we included in our qualitative reviews and quantitative meta-analyses, there were a total of 391 participants (DMT/ayahuasca: 52 participants; LSD: 148 participants; psilocybin: 191 participants). The meta-analysis of the BOLD activation literature included data from 258 participants, and the meta-analysis of the FC literature was performed on data from 237 participants.

##### *Section S1.3. Pharmacology*

We conducted two separate meta-analyses on the pharmacology of psychedelics: one on the selective affinities of psychedelics for three G protein-coupled receptors (GPCRs) – $5\text{-HT}_{2A}$, $5\text{-HT}_{2C}$, and $D_2$, relative to $5\text{-HT}_{1A}$ – and another on the functional activity of psychedelics, as measured by assays on GPCR-mediated signaling.

We performed our literature searches in May-August 2023. We searched the following four databases: SCOPUS; Web of Science; the National Institute of Mental Health's Psychoactive Drug Screening Program (PDSP) $K_i$ Database; and ChEMBL, the "chemogenomic" database of the European Bioinformatics Institute, which is part of the European Molecular Biology Laboratory

(EMBL). The PDSP $K_i$ Database (pdsp.unc.edu/databases/pdsp.php) is an online resource that contains nearly 100,000 published affinity ($K_i$) values for a very large number of chemicals and receptors (Roth et al., 2000). ChEMBL contains a wide variety of data on the chemistry, bioactivity, and genomics of more than 2.2 million compounds, including not only affinity ($K_i$ and $K_d$) and functional activity ($EC_{50}$) values but also information on chemical structures, molecular properties, absorption, metabolism, toxicity, and more (Davies et al., 2015; Mendez et al., 2019). Neither the PDSP $K_i$ Database nor the ChEMBL Database is comprehensive, as we found many studies on affinity and functional activity that were not included in the databases. Finally, there were three studies that we found through citations in articles that we had already included, but were not listed in any of the above databases.

We applied the following search terms for the literature search on binding affinities: <drug name> and "receptor affinity", <drug name> AND "inhibitory constant" OR Ki, <drug name> AND "binding affinity", <drug name> AND <receptor name> and affinity, where <drug name> was "DMT", "LSD", or "psilocin", and <receptor name> was "5-HT1A", "5-HT1B", "5-HT2A", "5-HT6", "D1", or "D2". For the literature search on functional activity, we used a single search term: <drug name> AND EC50 OR potency OR "functional assay", where <drug name> was "DMT", "LSD AND 'lysergic acid diethylamide'", or "psilocin". We excluded studies that met the following criteria:

1. Works that did not report any original research, such as literature reviews, commentaries, and editorials.
2. Studies that did not measure the binding affinities of DMT, LSD, or psilocin. Note that we excluded data on ayahuasca and psilocybin. A metabolite of psilocybin, psilocin is the main pharmacologically active substance in magic mushrooms (Passie et al., 2002).
3. Studies that did not examine the following receptors: $5\text{-HT}_{1A}$ and $5\text{-HT}_{2A}$, $5\text{-HT}_{2C}$, or $D_2$. We selected these receptors because we knew in advance that there are multiple studies that analyse the affinity of psychedelics for each of them. They are also relevant for the effects of psychedelics (de Vos et al., 2021; Nichols, 2016).
4. Affinity studies that did not measure $K_i$ or functional activity studies that did not measure GPCR-mediated signaling. While affinities are often measured as $K_d$ (dissociation constant) in pharmacology, studies on $K_d$ typically require a radionuclide to be attached to the drug of interest, in order to radioactively label the drug's interaction with the relevant receptor. However, to our knowledge, there are no studies that have attached radionuclides to psilocybin or DMT, only to LSD. Therefore, we only included $K_i$ data in the binding affinity meta-analysis. For the functional activity literature, we focused on GPCR functional assays because GPCR signalling appears to be a key mechanism underlying the hallucinogenic effects of psychedelics (López-Giménez & González-Maeso, 2018). The datapoint of interest in functional GPCR assays was $\Delta\log(E_{max}/EC_{50})$ (see *Section 2.3*).
5. Studies that did not report original $K_i$, $E_{max}$, or $EC_{50}$ data.
6. Studies that performed original measurements of $K_i$, $E_{max}$, or $EC_{50}$, but did not publish numerical values of the results.

7. Studies that did not specify the receptor subtype, e.g. studies on 5-$HT_1$ binding affinity that did not indicate whether they were measuring affinity for 5-$HT_{1A}$, 5-$HT_{1B}$, or one of the other subtypes of the 5-$HT_1$ family. We separated data from different receptor subtypes, so we did not wish to conflate them. Some older studies were published at a time when the receptor subtypes had not been discovered yet. However, we did include binding affinity studies on 5-$HT_1$ and 5-$HT_2$ if the radioligand, i.e. the radioactive chemical that competes with the drug for the receptor, used in the study has been shown to selectively bind to specific receptor subtypes. For instance, the radioligand [$^3$H]-8-OH-DPAT is quite selective for the 5-$HT_{1A}$ subtype (Hjorth et al., 1982), whereas the radioligand [$^3$H]-5-HT is not (Davis et al., 2002). Therefore, for studies on 5-$HT_1$ affinity that used [$^3$H]-8-OH-DPAT, we included their results in our analysis of affinity for 5-$HT_{1A}$.
8. Studies that did not examine human receptors.
9. Studies that did not specify the source of receptor (e.g. whether the receptor was cloned or extracted from brain homogenates) or the species of the animal from which the receptor was derived.

A flowchart of the pharmacology literature search is shown in **Figure S1c.** We identified ***n* = 28 studies in total**. These included ***n* = 14 studies on affinity** (Chadeayne et al., 2020; Erkizia-Santamaría et al., 2022; Eshleman et al., 2014, 2018; Glatfelter et al., 2022; Halberstadt et al., 2020; Janowsky et al., 2014; Keiser et al., 2009; Kozell et al., 2023; Luethi et al., 2018, 2019; Pierce & Peroutka, 1989; Rickli et al., 2015, 2016), as well as ***n* = 18 studies on functional activity** (Braden & Nichols, 2007; Cimadevila et al., 2020; Cussac et al., 2008; Eshleman et al., 2014, 2018; Gatch et al., 2011; Glatfelter et al., 2022; Halberstadt et al., 2020; Janowsky et al., 2014; Keiser et al., 2009; Klein et al., 2021; Kozell et al., 2023; Newton et al., 1996; Porter et al., 1999; Pottie, Cannaert, et al., 2020; Pottie, Dedecker, et al., 2020; Schmitz et al., 2022; Sherwood et al., 2020).

Note that we first conducted a meta-analysis on affinity relative to the 5-$HT_{1A}$ receptor. In the Supplementary Materials, we also report the results of a meta-analysis on affinity relative to the 5-$HT_{2A}$ receptor, for which we included an additional three studies (Blair et al., 1999; Cussac et al., 2008; Knight et al., 2004).

While the majority of studies reported standard errors (SEM), some published standard deviations (SD) instead but did not explicitly state the number of corresponding experiments, which is needed in order to transform SD into SEM. In these cases, we inferred the number of experiments from other assays performed in the study.

### Section S2. Methods

For the pharmacology and phenomenology data, we performed random-effects meta-analyses, using the dmetar package in R (Harrer et al., 2021). We applied the GingerALE algorithm to conduct the meta-analysis of the BOLD activation studies, and we created a novel method to perform a meta-analysis on the FC studies, given the lack of adequate methods for synthesising FC data.

In order to relate the different levels of the hierarchy to one another, we determined the "profile" of the pharmacology, neuroimaging (BOLD activation and FC), and pharmacology of each psychedelic: DMT, LSD, and psilocybin. Each profile described the alignment between the seven resting-state Yeo networks (Thomas Yeo et al., 2011) and the neural correlates of each level of the hierarchy. These networks are the visual network, somatomotor network (SMN), dorsal attention network (DAN), ventral attention network (VAN), limbic network, frontoparietal network (FPN), and default mode network (DMN).

##### *Section S2.1. Phenomenology*

There are two versions of the Altered States of Consciousness: one that measures five dimensions (5D) of consciousness and another that measures eleven dimensions (11D). While the eleven dimensions are subscales of the five dimensions, the two versions of the questionnaire do not have an equal number of questions, so the 11D data cannot be averaged across subscales to obtain 5D data. Therefore, we had to perform separate meta-analyses of the 5D- and 11D-ASC data.

In addition to measuring multiple subjective dimensions, ASC studies administered multiple doses of a psychedelic to the same group of participants, and a few also gave different psychedelics to the same group. Therefore, the ASC scores are not statistically independent from one another, which violates a core assumption of a traditional meta-analysis. A three-level meta-analysis accounts for these statistical dependencies (Harrer et al., 2021). The three levels are: (1) the effect sizes, or ASC scores, of individual participants; (2) the pooled effect size, or overall ASC scores, of each dimension, dose, and drug; and (3) the pooled effect size of the whole study. Thus, in this model, multiple effect sizes can be nested within one study. The three-level model fits the data to the following formula (Assink & Wibbelink, 2016; Cheung, 2014):

$$\hat{\theta}_{ij} = \mu + \beta x_i + \zeta_{(2)ij} + \zeta_{(3)ij} + \epsilon_{ij}$$

where $\hat{\theta}_{ij}$ is an estimate of the true effect size $i$ (ASC score) nested in study $j$, $\beta$ is the regression weight of a predictor $x$, $\mu$ is the true (population) effect size, $\zeta_{(2)ij}$ is the within-study heterogeneity, $\zeta_{(3)ij}$ is the between-study heterogeneity, and finally $\epsilon_{ij}$ is the sampling error.

To account for the fact that not only effect sizes but also sampling errors within studies are correlated with each other, we specifically applied an extension of the three-level model known as the Correlated and Hierarchical Effects (CHE) model with Robust Variance Estimation (Hedges et al., 2010; Pustejovsky & Tipton, 2022; Tipton & Pustejovsky, 2015). Essentially, this technique estimates the true dependence structure (variance-covariance matrix) between effect sizes within the same study, given a known correlation coefficient $\varrho$ between those effect sizes. We assumed $\varrho = 0.6$, as a previous meta-analysis recommended this correlation coefficient (Harrer et al., 2021); however, the choice of $\varrho$ was apparently arbitrary in this meta-analysis, and we did not run any sensitivity analyses to confirm that this $\varrho$ was appropriate. When the number of studies in a meta-analysis is small (under 40, which is the case for this meta-analysis), traditional variance estimators tend to be biased. We reduced the risk of bias by adjusting the estimated variance with bias-reduced linearisation (CR2 method) (Tipton & Pustejovsky, 2015).

In our model, effect sizes were pooled using the generic inverse variance method (see *Section S2.3*). Initially, the only predictor $x$ was the interaction between drug, dose, and dimension. Within- and between-study heterogeneity were estimated with the restricted maximum-likelihood procedure. We applied the Knapp-Hartung method to adjust the standard errors of the $\beta$ weights in order to provide more conservative estimates of the $p$-value (Knapp & Hartung, 2003).

DMT was administered intravenously in the phenomenological studies whereas LSD and psilocybin were administered orally (we excluded phenomenological studies that used IV LSD and psilocybin). IV administration yields different pharmacokinetics, which are known to influence the subjective experience of the drug (Luan et al., 2023; Vogt et al., 2023). Therefore, while we conducted meta-analyses of all three drugs, we only examined significant differences between ASC ratings of LSD and psilocybin. Additionally, we only measured significance for similar doses of LSD and psilocybin. We assumed 20 mg psilocybin is equivalent to 0.01 mg LSD (Ley et al., 2023), and we defined low, medium, and high doses based on the literature (Carbonaro et al., 2018; Hasler et al., 2004; Vollenweider et al., 2007). A low dose is 8-14 mg psilocybin or 0.05-0.074 mg LSD, a medium dose is 15-21 mg psilocybin or 0.075-0.109 mg LSD, and a high dose is ≥22 mg psilocybin or ≥0.110 mg LSD. (Note that 1 microgram = 0.001 mg.) Any findings that corresponded to quantities of LSD or psilocybin that were below a low dose were excluded from the study, as many (though not all) of these doses were microdoses. Many phenomenological studies on psilocybin adjusted dosing based on the weight of the participant, whereas phenomenological studies on LSD tended to administer the same dose to all participants. To make the data consistent, we assumed an average weight of 70 kg for participants in the psilocybin studies. It is worth noting that, in the literature, a medium dose of LSD was almost always 0.010 mg, whereas a medium dose of psilocybin tended to be 0.215 mg/kg, which we considered to be equivalent to 15.05 mg. Therefore, the medium dose of LSD tended to be higher than the medium dose of psilocybin.

To model the influence of methodological variables that may confound the results, we produced a third model in which we included timing (i.e. the length of time between the administration of the drug and of the questionnaire), the presence or absence of a task in the experiment, and the prior psychedelic use of the participants as extra covariates in addition to the interaction between dose, scale, and drug. To compare the fit of this "full" model and one of the "reduced" models, in which we did not include these covariates, we examined the corrected Akaike's information criterion (Akaike, 1998).

We measured publication bias by assessing small-study effects. That is, small studies tend to have large standard errors, so only small studies with disproportionately high effect sizes produce significant results (Borenstein et al., 2009; Harrer et al., 2021). Specifically, we performed Egger's regression test, which determines whether the ratio between the observed effect sizes and standard errors is close to zero when the standard errors are infinitely large (Egger et al., 1997). If the ratio is significantly different from zero, then small-study effects are present. We did not measure other sources of publication bias; we deemed that it was unlikely for there to be unpublished studies with insignificant results or for studies to perform *p*-hacking, due to the marked perceptual effects of psychedelics relative to placebo.

Publication bias pertains to the entire body of literature included in the meta-analysis. We assessed risk of bias for individual studies by considering several domains of bias covered in the Risk Of Bias in Non-randomized Studies of Interventions (ROBINS-I) tool: bias due to confounding, bias in selection of participants, bias in classification of interventions, bias due to deviations from intended interventions, bias due to missing data, bias in measurement of outcomes, and bias in selection of the reported result (Sterne et al., 2019). Two authors independently examined risk of bias in each study. We used the GRADE approach (Grading of Recommendations Assessment, Development and Evaluation) to assess our level of certainty in the body of evidence (Schünemann et al., 2019).

We compared the different levels of the hierarchy – phenomenology, neuroimaging, and pharmacology – by measuring the "profile" of each level, or the neural correlates of each level in the Yeo networks. In order to obtain the phenomenological profile of each psychedelic, we needed to define the neural correlates of their subjective effects. In general, we believe that subjective experience is encoded in a dynamic and flexible fashion across the whole brain, rather than in a static and specific subset of regions (Barttfeld et al., 2015; Boly et al., 2013; Dehaene & Changeux, 2011; Oizumi et al., 2014). Therefore, it is unlikely that the subjective experiences of psychedelics, which are extraordinarily rich and complex, correlate with a single configuration of Yeo networks. Nevertheless, the subjective profile that we have created in this paper provides a starting point for comparing the phenomenology with the neuroimaging and pharmacology.

We identified $n = 14$ studies that examined correlations between subjective effects and either fMRI activity or pairwise connectivity. We assigned a Yeo network to each region that was significantly correlated with an ASC dimension, based on the coordinates of the centroid of the region. We then averaged correlations within the same Yeo network. This resulted in a series of five 7x1 vectors, one for each ASC dimension, in which the components of the vector correspond to the correlations between that dimension and the corresponding Yeo network. The phenomenology profile of each psychedelic was a weighted sum of the five vectors, in which the weights were the pooled 5D-ASC scores for the corresponding dimension.

$$\mathbf{s}^{(p)} = \sum_{c=1}^{5} w_d^{(p)} \mathbf{r}^{(d)}$$

where $\mathbf{s}^{(p)}$ is a 7x1 vector representing the phenomenology profile of each psychedelic $p$, $w_d$ is the pooled 5D-ASC rating of each psychedelic for each subjective dimension $d$, and $\mathbf{r}^{(d)}$ is a 7x1 vector containing the correlations between the Yeo networks and the neural correlates of each dimension.

##### *Section S2.2. Neuroimaging*

The fMRI studies on psychedelics measure three different characteristics of brain activity: BOLD activation, functional connectivity, and entropy. We used the GingerALE software to determine common clusters of BOLD activation across studies, quantified overall changes in functional connectivity between the Yeo networks under psychedelics, and conducted a qualitative review of the research on entropy.

*Section S2.2.1. BOLD activation*

As discussed in *Section 2.2* of the main text, we did not report the results of our BOLD activation meta-analysis because we discovered that the experimental procedures likely confounded the results. Nevertheless, we briefly report our method for this meta-analysis here.

We conducted our meta-analysis of BOLD activity by using a technique called GingerALE, which computes common clusters of BOLD activation across different studies (Eickhoff et al., 2009, 2012; Turkeltaub et al., 2011). GingerALE takes in a list of the coordinates (in either MNI or Talaraich space) of peak activations, or foci, from each study, and then it outputs the common clusters of activation. Our inputs into GingerALE were the MNI coordinates of peak BOLD activation for any contrast comparing brain activity in the placebo/baseline condition with brain activity under psychedelics. We applied a cluster-forming threshold of $p < 0.05$ and a cluster-level inference threshold of $p < 0.05$.

*Section S2.2.2. Functional connectivity*

In the FC literature, a wide range of parcellations are used to define regions of interest (ROIs). To perform a meta-analysis on studies of FC, we first needed to "re-parcellate" all of the data into the same reference parcellation. The reference parcellation that we chose was the seven resting-state Yeo networks, since we determined the profiles, or neural fingerprints, of the other hierarchical levels of analysis (phenomenology and pharmacology) based on these networks. Additionally, the Yeo networks are perhaps the most coarse-grained parcellation of the human brain. We conducted what amounted to a "vote-counting" meta-analysis (Bushman & Wang, 2009). Our algorithm determined the pair of Yeo networks that contained every pair of functionally connected ROIs in the literature. That is, we identified the Yeo networks that overlapped with the largest number of voxels in each ROI. If the pair of ROIs became more connected under psychedelics, then a "vote" was cast in favour of the corresponding Yeo networks; that is, the vote was counted as positive. If not, then a vote was cast against those Yeo networks; that is, the vote was counted as negative.

Studies that use parcellations with many seeds bias vote-counting FC meta-analyses because the number of significant connections is much higher, and hence these studies contribute much more to the aggregate FC. To avoid this bias, we weighted each study's vote for a pair of networks by the total number of *possible* connections between those networks in the study. For instance, if a study used a parcellation with 11 regions in the DMN and 8 regions in the FPN, then there are 11 * 8 = 88 possible connections between the DMN and FPN. If the study only found 22 significant connections between the DMN and the FPN, then the study would contribute a vote of 22/88 = 0.25 to the aggregate DMN-FPN FC. This method substantially reduces the contribution of studies with large parcellations to the aggregate FC.

Our quantitative meta-analysis included any studies on pairwise FC, namely studies that performed one of the following analyses: seed-to-seed analysis; seed-to-voxel analysis; or component-to-component analysis, in which the components are identified through Independent Component

Analysis (ICA). We qualitatively reviewed studies that conducted other types of FC analyses in *Section S5*.

Because most studies did not publish standard errors or *p*-values, we could not perform the quantitative analyses of publication bias that we conducted in the pharmacology and phenomenology meta-analyses. Our qualitative review revealed one major source of publication bias: many of the FC studies were secondary analyses of preexisting datasets, and some datasets, such as those of Carhart-Harris, Erritzoe, et al. (2012) and Carhart-Harris et al. (2016), were analysed many more times than others. To prevent bias from primary datasets that received more than one analysis, we selected the most informative analysis of each unique dataset, unless the secondary analysis addressed a part of the dataset that was not reported in the original publication. (For instance, Kaelen et al. (2016) is an analysis of the music condition in the primary dataset collected by Carhart-Harris et al. (2016), which reported analyses of only the resting-state condition. Therefore, we included Kaelen et al. (2016) despite the fact that it was a secondary analysis.) We defined the most informative analysis as the most spatially extensive, fine-grained one, i.e. the analysis that examined FC across the largest portion of the brain, using the parcellation with the highest number of ROIs. We further excluded secondary analyses in which the number of significant connections did not scale as expected with the size of the parcellation. In general, we expect the number of significant between-ROI connections to scale roughly quadratically with the parcellation size, since the number of possible connections is approximately the square of the number of regions in the parcellation. However, in Lebedev et al. (2015) and Luppi et al. (2021), the number of significant connections is smaller than the parcellation size. This effect likely occurred because both studies were the only ones in the literature to use the network-based statistic (NBS) to determine statistical significance. Unlike other techniques, NBS measures the size of interconnected "components" in brain networks, rather than assessing each possible connection individually (Zalesky et al., 2010). Since there are fewer components than pairwise connections, the number of significant connections determined by NBS is likely to be much lower than the number of significant connections identified by other techniques, such as mass-univariate *t*-tests. Thus, even though Lebedev et al. (2015) and Luppi et al. (2021) were both whole-brain secondary analyses that used more fine-grained parcellations than other analyses did, we excluded them because their method underestimated the number of significant connections, relative to the other techniques in the literature. **Table S4** lists all the primary datasets in the literature on pairwise FC, the secondary analyses, and the studies that we included in the FC meta-analysis.

What were the inputs into the meta-analysis? **Table S5** specifies the data that we inputted from each of the 12 studies. For seed-to-seed analyses, we obtained spatial maps of the seeds from the relevant parcellation or from the authors of the study, when special techniques were used to extract the seed (Barrett, Krimmel, et al., 2020). One seed-to-seed analysis measured FC between pairs of networks by averaging across the FC of 10mm-radius spheres constructed around the centroids of the constituent ROIs in those networks (Madsen et al., 2021). To incorporate the data from this study into the meta-analysis, we used the MarsBar toolkit (Brett et al., 2002) to generate 10mm-radius spheres around the coordinates of each ROI centroid. For each network, we determined the Yeo network that contained each of the constituent ROIs and then identified the mode, i.e. the Yeo network that overlapped with more of the constituent ROIs than any other Yeo network. FC between mode networks then contributed towards the aggregate FC matrix.

For seed-to-voxel analyses, we reached out to the authors for spatial maps of the voxels that displayed significant changes in FC with the corresponding seeds. When these maps were unavailable (Grimm et al., 2018), we created spheres around the reported peak coordinates of each significant cluster of voxels. The radius of each sphere corresponded to the size of the corresponding cluster. We then assigned each cluster to a Yeo network.

Finally, the three ICA-based analyses that we included in the meta-analysis (Gaddis et al., 2022; Mason et al., 2020; Roseman et al., 2014) found strong correlations between independent components and resting-state networks (RSNs) derived by Smith et al. (2009), or they directly back-projected those RSNs onto their participants' data through dual regression. In order to help unify the three studies, our inputs for all three studies were the RSNs defined by Smith et al. (2009), not the group ICs.

To compute the significance of our results, we compared our results to a surrogate dataset formed from the resting-state BOLD timeseries of 100 unrelated participants in the Human Connectome Project (HCP) (Van Essen et al., 2013). We parcellated this timeseries with the Schaefer-100 parcellation (Schaefer et al., 2018). To do this, we randomly sampled $N$ subjects from the HCP data, and two length-$T$ windows from each subject's BOLD data, where $N$ and $T$ are the average number of subjects and the average number of TRs, respectively, in the studies of FC under psychedelics. In each window, we calculated the Pearson correlation between the ROIs of the Schaefer-100 parcellation. The pairs of ROIs that were significantly correlated constituted the results of one surrogate study. (Note that we did not perform multiple comparisons on the correlations, since otherwise there would have been no significant edges.) Since there are 12 studies of FC in our meta-analysis, we repeated the FC calculation 12 times, resulting in a "surrogate study set" of significantly connected ROIs. We then computed the aggregate FC matrix of this surrogate study set. We repeated this process 500 times, which produced a null distribution of values for each element of the aggregate FC matrix. If an element of the aggregate matrix for the psychedelic data was below the 2.5$^{th}$ percentile or above the 97.5$^{th}$ percentile of the null distribution, then it was deemed to be significant. Finally, the aggregate FC matrix was corrected for multiple comparisons with the Benjamini-Hochberg FDR procedure.

We stress that our method for performing an FC meta-analysis is novel and has not been validated elsewhere. Indeed, there are reasons to think that the surrogate method does not accurately assess statistical significance. When measuring resting-state FC in each surrogate study, we calculate the correlations between all pairs of ROIs in the Schaefer-100 parcellation. On the other hand, in the literature, some studies analyse connectivity between only a subset of the ROIs from the chosen parcellation.

While the FC meta-analysis was initially performed on the cortical Yeo networks, we later expanded the analysis to include eight subcortical regions from the Tian atlas – the hippocampus, amygdala, posterior thalamus, anterior thalamus, nucleus accumbens, globus pallidus, putamen, and caudate – plus cerebellum (Tian et al., 2020). Since the Schaefer-100 atlas only contains cortical regions, we used the Desikan-Killiany (DK-80) atlas, which includes 62 cortical and 18 subcortical regions, to parcellate the null distribution for the cortical + subcortical analysis (Deco et al., 2021; Desikan et al., 2006).

As we did in the phenomenology and pharmacology meta-analyses, we tried to identify methodological covariates that were significantly associated with the results. We identified eight potential covariates *a priori*: (1) the method used to regress out white matter from BOLD signal (in particular, whether anatomical Component-Based Noise Correction (aCompCor) was used); (2) whether or not the study performed scrubbing in order to correct for head motion; (3) whether or not the study preprocessed fMRI data with FSL; (4) whether or not the study used SPM for preprocessing; (5) whether or not the study administered psilocybin; (6) the route of administration (either oral or IV); (7) the width of the Gaussian kernel that was used to smooth the data; and (8) the time that the drug was administered, relative to the duration of the drug's acute effects (7.5 hours for ayahuasca, 20 minutes for DMT, 10 hours for LSD, 5 hours for psilocybin). (1)-(5) are binary covariates, whereas (6)-(8) are continuous.

For binary covariates, we compared the aggregate FC matrices of studies that contained the covariate in question to those that did not, by taking the absolute value of the difference between each entry of the corresponding matrices. Then, we scrambled the labels of the covariates and re-compared the score matrices for the permuted data. We repeated this process approximately 500 times. Our analysis was hampered by the small number of studies; there were fewer than 495 unique permutations when fewer than four studies exhibited a particular covariate. Thus, we could not determine whether the FC of psilocybin and LSD were significantly different from each other, as we only analysed three studies on the FC of LSD. We could only assess whether psilocybin's FC was significantly different from the rest of the FC dataset, as we included six studies on the FC of psilocybin. For continuous covariates, we assessed the significance of the correlation between each entry of each study's marginal contribution to the aggregate FC matrix and each study's value for the covariate in question. We adjusted for multiple comparisons with FDR for both types of covariates.

In addition to correcting for publication bias stemming from the large number of secondary analyses in the literature, as discussed above, we assessed risk of bias in individual studies using the ROBINS-I tool. Two authors independently performed the assessment.

Finally, we obtained the FC profiles by first determining the aggregate FC matrix of each psychedelic. Then, we summed across the rows of the matrix (note that the matrix is symmetric, so the choice of row or column does not matter) and divided by the sum of all values in the matrix.

*Section S2.2.3. Entropy*

See *Section S4*.

##### *Section S2.3. Pharmacology*

We performed two meta-analyses of the pharmacology literature. To synthesise the data on binding affinity, we analysed selectivity, measured as the ratio of the $K_i$ for the three receptors of interest (5-$HT_{2A}$, 5-$HT_{2C}$, and $D_2$) relative to the $K_i$ for 5-$HT_{1A}$. We also performed a supplementary analysis on selectivity relative to 5-$HT_{2A}$. To analyse the data on functional activity, we pooled relative activity values, defined as $\Delta\log(E_{max}/EC_{50})$ (see *Section S6.3* for justification).

All GPCR functional assays, at least in humans, were performed on cloned cell lines. The majority of affinity studies were also conducted on cloned cell lines, and a handful of affinity studies carried out their experiments on brain homogenates. While some of the latter studies reported multiple data points, since they measured affinity for multiple drugs or receptors, we treated the data points in these studies as statistically independent from one another, as it was not apparent that the homogenates were extracted from the same brain. Thus, we chose not to perform a multilevel meta-analysis as we did for the phenomenology literature.

Instead, for both the affinity and activity meta-analyses, we constructed random-effects models, which, unlike fixed-effects models, assume that true effect sizes ($K_i$ and $EC_{50}$) are not uniform across studies once sampling errors are accounted for (Hedges & Vevea, 1998). Rather, the true effect sizes exist along a distribution with an overall mean $\mu$. The true effect size of a particular study $k$ deviates from the mean $\mu$ of all true effect sizes by an error $\zeta_k$, and the actual effect size of one study deviates from its true effect size by the sampling error $\epsilon_k$. Therefore, the random-effects model is:

$$\hat{\theta}_k = \mu + \beta x_k + \zeta_k + \epsilon_k$$

where $\hat{\theta}_k$ is the actual effect size of study $k$, $x$ is a predictor, and $\beta$ is its weight.

We chose random-effects models over fixed-effects models due to the large heterogeneity that we observed in the data. In order to account for $\zeta_k$, we must estimate the variance of the distribution of true effect sizes, known as $\tau^2$ (Harrer et al., 2021). There are several well-validated estimators of $\tau^2$; we chose the restricted maximum likelihood procedure, which has been shown to outperform other estimators for continuous data like ours (Veroniki et al., 2016). Once $\tau^2$ has been estimated, we can calculate a weight for each data point:

$$w_k^* = \frac{1}{s_k^2 + \tau^2}$$

where $s$ is the square of the standard error. Then, the pooled effect size is calculated as follows, in what is known as the generic inverse variance method:

$$\hat{\theta} = \frac{\sum_{k=1}^{K} \hat{\theta}_k w_k^*}{\sum_{k=1}^{K} w_k^*}$$

For both the selectivity and functional activity analyses, we initially modeled the drugs as the sole covariates. We applied the Knapp-Hartung adjustment to produce more conservative estimates of variance in the $\beta$-weight of the predictor (Knapp & Hartung, 2003). However, we stress that the range of variances was still quite large, and the number of studies on some receptors and functional assays was small. (For instance, there are only four studies that we included on selectivity for the $D_2$ receptor.) Therefore, our estimates of $p$-values may not be reliable.

For just the selectivity analysis, we created another model in which we included the radioligand as an additional covariate, since the radioligand is known to influence the $K_i$ estimate. To compare the fit of this "full" model to the fit of the "reduced" models (i.e. the model where only the choice of radioligand was not modelled), we examined the corrected Akaike's information criterion (Akaike, 1998). Because relative activity already "cancels out" differences in assay protocols, we did not model any additional covariates for the functional activity analysis.

We measured publication bias with the same method that we used for the phenomenology meta-analysis (*Section S2.1*). Note that omission of studies due to insignificant *p*-values is not a concern in the pharmacology literature. The studies either did not measure significant between-drug differences in binding affinity or functional activity, or they used one of the classical psychedelics as the control drug in these comparisons. ROBINS-I, the tool that we used to assess risk of bias for individual studies in the phenomenology meta-analysis, does not pertain to the pharmacology literature, since ROBINS-I assesses bias for interventional trials. Instead, we discuss some potential confounders in the literature in *Section S6.3*. As we did in the phenomenology meta-analysis, we used the GRADE tool to assess our level of certainty in the body of pharmacological evidence.

In order to obtain the pharmacology profile of psychedelics, we needed to identify the neural substrate of their pharmacology in the human brain. We based the pharmacology profile on the binding affinity data, rather than the functional activity data, because it was unclear how to combine data about the three different cellular signaling pathways. We defined the pharmacology profile as a weighted combination of the overall receptor density in each Yeo network, weighted by the selectivity of the psychedelics:

$$\mathbf{h}^{(p)} = \sum_{r=1}^{2} S_r^{(p)} \mathbf{l}^{(r)}$$

where $S_r^{(p)}$ is the selectivity of a psychedelic *p* for a receptor *r*, relative to the 5-$HT_{1A}$ receptor; $\mathbf{l}^{(r)}$ is a 7x1 vector containing the total expression level of each receptor in each Yeo network; and $\mathbf{h}^{(p)}$ is a 7x1 vector representing the pharmacology profile of each psychedelic.

To determine $\mathbf{l}^{(r)}$, we took human PET maps of 5-$HT_{2A}$ and $D_2$ from Hansen et al. (2022), which itself obtained and in some cases averaged maps from previous literature. (Unfortunately, there was no map of the 5-$HT_{2C}$ receptor in this atlas.) The 5-$HT_{2A}$ map was taken from Beliveau et al. (2017) and the $D_2$ map was averaged from data by Sandiego et al. (2015) and C. T. Smith et al. (2019). The 5-$HT_{2A}$ study used the [$^{11}$C]-CIMBI-5 PET radiotracer, and both $D_2$ studies used the [$^{11}$C]-FLB457 PET radiotracer. Hansen and colleagues parcellated these maps with the Schaefer-200 atlas. We then performed min-max normalisation on the maps, which turned the lowest value of receptor expression into 0 and the highest into 1. To obtain $\mathbf{l}^{(r)}$, we then summed the normalised values of receptor expression across all regions belonging to each Yeo network.

We also constructed subcortical pharmacology profiles based on subcortical maps of 5-$HT_{2A}$ (Beliveau et al., 2017) and $D_2$ receptor expression (Malén et al., 2022). Because the subcortical $D_2$ map used a different radiotracer, [$^{11}$C]-raclopride, than the cortical map, we kept the cortical and subcortical pharmacology profiles separate.

#### Section S3. Summary of comparative phenomenological studies

To complement our meta-analysis of the 5D- and 11D-ASC scores, we qualitatively reviewed seven studies that directly compared the phenomenology of ayahuasca/DMT, LSD, and psilocybin. Zamberlan et al. (2018) conducted a latent semantic analysis (LSA), followed by a principal components analysis (PCA), on trip reports about 29 psychoactive substances, including LSD, DMT and psilocybin. Trip reports were taken from Erowid, a large, online, publicly available collection of "trip reports" written by individuals about their psychedelic experiences (Erowid et al., 2022). The aim of the study was to identify key themes that differentiate psychedelics from one another. Their findings showed that five principal components, or themes, cumulatively accounted for 58% of the variance between drugs: perception, body load, preparation (i.e. how the drugs are prepared in order to be consumed), dependence (e.g. addiction), and therapeutic. LSD ranked high on perception but relatively low on the other components. The DMT scores showed a similar pattern, except for the therapeutic component, which was the 2nd highest out of all the psychoactive substances (the top was ibogaine). The scores for psilocybin were not displayed in the paper.

Ballentine et al. (2022) applied LSA to 6850 Erowid testimonials about 27 drugs. They extracted the binding affinities of the drugs to 40 neurotransmitter receptor subclasses. A canonical correlation analysis was used to delineate factors relating themes of subjective experience to the receptor affinities of the associated drugs. Each factor is defined by a spectrum of phenomenological themes, such that the two ends of the spectrum tend to be anti-correlated with each other (for instance, the leading factor spanned from "visuals" to "reality"). Results of the first eight factors, representing the highest systematic variation between receptor affinity profiles and phenomenological themes, seemed to suggest a separation between DMT in one thematic extreme of 7/8 factors and LSD and psilocin in the other.

Sanz et al. (2018) also ran LSA on the Erowid dataset; however, their work examined the similarities between these reports and those of individuals who had experienced high and low lucidity dreams. All four compounds (reports of Ayahuasca and DMT use were studied separately) were found to be experientially similar to both dream states. LSD was ranked the most similar out of the four in both cases. Out of 161 hallucinogenic compounds, LSD, psilocybin, DMT and ayahuasca were respectively ranked 1st, 4th, 15th, 11th in similarity to high lucidity dreams.

Qiu & Minda (2021) carried out a sentiment analysis using a sentiment dictionary that assigns an emotional valence score to almost 2500 words, thereby quantifying how happy or sad a word is. Words in Erowid trip reports were matched with reference words in the AFINN lexicon and were subsequently assigned an emotional valence. Their results showed that the DMT group had significantly higher (i.e. happier) sentiment scores than either the psilocybin or LSD groups. The authors then analysed a dataset that consisted of responses from individuals who had completed the MEQ, along with a variety of wellbeing psychometrics. In this dataset, they found that LSD and DMT were significantly associated with a greater likelihood of occasioning a complete mystical experience, unlike psilocybin.

Coyle et al. (2012) used 1000 reports from the Erowid dataset across 10 drugs. They used the frequency of individual words in each report to predict which drug was taken, using a machine learning technique known as random-forest classification. They were able to reach high accuracy for DMT and LSD reports: 52% and 40%, respectively. However, the accuracy dropped to 25% for psilocybin reports.

Griffiths et al. (2019) conducted a large-scale survey using a 76-item questionnaire examining the subjective experience of an individual's single most memorable "god encounter" moment. The participants of the survey included not only people who had taken psychedelics before, but also anyone who had ever had a profound religious experience. The psychedelic participants were separated into four groups (with ayahuasca and DMT being considered separately). Analysis of the survey revealed little to no difference between the reports of LSD and psilocybin. These two substances were, however, found to have significantly different rankings than DMT and ayahuasca on several statements. For example, those who had taken ayahuasca or DMT were more likely to support the statement that the encounter was not initiated by them, but rather by the encountered being, as well as the statement that they communicated with the encountered being. Additionally, a significantly higher percentage of people within the DMT group achieved a complete mystical experience. Those in the ayahuasca condition had the highest scores for the vividness of their memories of the "god encounter" moment, while those in the DMT group had the lowest.

Finally, Hase et al. (2022) employed LSA to quantify the semantic similarity between each report in another Erowid dataset and two reference texts, which were scales for measuring psychedelic (an Altered States of Consciousness Scale) and mystical (Hood's Mysticism Scale) experiences, respectively. The ayahuasca + DMT condition had significantly higher similarity to the ASC Scale and Hood's Mysticism Scale than both LSD and psilocybin. LSD reports had significantly higher similarity to Hood's Mysticism Scale than psilocybin reports. Hase et al. (2022) also used Linguistic Inquiry and Word Count to calculate the percentage of words in a given experiential report which fit one or more pre-defined psychologically meaningful categories. The word categories analysed in this study included: positive emotion, negative emotion, sadness, anxiety, cognitive processes, social processes, biological processes, vision-related, health-related, drive-related, affiliation-related, reward-related, and space-related words. The subjective scores for each psychedelic appear to be very similar to one another; the range of scores within a category (never greater than 1.42) is much smaller than the range of scores across categories (11.92). Nevertheless, the ayahuasca+DMT reports diverged significantly from LSD and psilocybin for seven word categories (negative emotion, sadness, social processes, vision-related, drive-related, affiliation-related, reward-related). On the other hand, psilocybin and LSD reports were significantly different across only two categories: biological processes and reward-related words. The two categories with the largest range of scores were social processing and "drive-related" experience (i.e. motivation), with LSD ranking nearly 1.4 points higher than ayahuasca+DMT on both counts. The sadness and anxiety categories exhibited the smallest ranges: 0.05 and 0.07, respectively. While we expected that cognitive processing would score very highly in the trip reports for all three psychedelics (score range: 11.77-12.06), it was surprising that vision-related words were much less common (score range: 1.65-1.96), given the strongly distortionary effects that each of these psychedelics is known to have on visual perception (Kraehenmann, 2017).

### Section S4. Qualitative review of entropy studies

##### *Section S4.1. Entropic Brain Hypothesis*

The $n$ = 12 fMRI studies on entropy and criticality in the psychedelic literature are very heterogeneous, employing a wide variety of measures on dissimilar variables of interest. A quantitative meta-analysis was extremely challenging, so we opted instead to perform a qualitative review of this literature.

A seminal theory known as the Entropic Brain Hypothesis proposes that the subjective effects of psychedelics, such as ego dissolution, are associated with increased levels of entropy in the brain (Carhart-Harris et al., 2014). While entropy is typically associated with the amount of disorder in a system, it is more accurate to think of entropy as a measure of variability or unpredictability. In statistical mechanics, the entropy of a system of particles is proportional to the number of possible microscopic states, i.e. states of each particle, that are consistent with the macroscopic state of the system. (A box in which all the particles are clustered together in one corner is less entropic than a box in which the particles are scattered.) Similarly, according to EBH, the brain would be more entropic if, for instance, individual regions were to exhibit a greater number of possible connectivity motifs. Tagliazucchi et al. (2014) found that psychedelics raise entropy in this manner, and since then, ten other fMRI studies and several MEG and EEG studies have all shown that entropy increases during psychedelic use (Alonso et al., 2015; Atasoy et al., 2017; Lebedev et al., 2015, 2016; Lord et al., 2019; Luppi et al., 2023; Mediano et al., 2020; Pallavicini et al., 2021; Schartner et al. 2017; Singleton et al., 2022; Timmermann et al., 2019, 2023; Varley et al., 2020; Viol et al., 2017, 2019).

Entropy was formally defined as a measure of informational uncertainty by Claude Shannon (Shannon, 1948). Shannon entropy is equivalent to the *breadth* of possible outcomes for a random variable. If only a single outcome ever occurs with a probability of 1, then Shannon entropy is minimised at zero. Otherwise, if a broad range of outcomes can occur, each with low probability, then Shannon entropy will increase. Thus, in neuroscience, greater (Shannon) entropy refers broadly to an increase in the range of possible neural configurations. However, there are many different ways to define a neural configuration, and each study in the psychedelic literature measures a different configuration of interest. Some assess the variance of intra-network synchrony, whereas others consider the diversity of connections to clusters of brain nodes. Therefore, while most studies demonstrate an increase in Shannon entropy under psychedelics (Carhart-Harris et al., 2014; Luppi et al., 2023; Tagliazucchi et al., 2014; Viol et al., 2017, 2019), none of them are measuring changes in entropy *of the same variable*.

Shannon entropy is closely related to compressibility, which three of the other studies examine (Ruffini et al., 2023; Singleton et al., 2022; Varley et al., 2020). Indeed, a fundamental theorem in information theory, the Source Coding Theorem, states that entropy is a lower bound on the length of the optimal code for compressing data into a given set of symbols (e.g. a binary code of 0s and 1s) (Shannon, 1948). In the psychedelic literature, compressibility is measured as Lempel-Ziv complexity (LZc), which converges to the entropy rate when normalised (Ziv & Lempel, 1978). LZc determines the minimum number of unique substrings that are needed in order to decompose a string (Lempel & Ziv, 1976; Ziv & Lempel, 1977, 1978). For instance, if the string in question is a binary series of zeros and ones, then the LZ algorithm will compute the "building blocks" of binary patterns (e.g. "00101" and "1011") that comprise the entire string. If the string is a repeating sequence (e.g. "010101…"), then LZc will be minimised, as the string can be constructed by writing the same substring (in this case, "01") over and over again. If the string is completely random (e.g., independent flips of a fair coin), then LZc will be maximised, as the set of substrings that make up the entire string is infinitely large. All three fMRI studies on compressibility conclude that psychedelics increase LZc (that being said, Varley et al. (2020) only find significant differences for LSD, but not for psilocybin). This result is consistent with the findings of MEG studies on LZc under psychedelics (Mediano et al., 2020; Schartner et al., 2017; Timmermann et al., 2019).

The last of the three categories of entropy measures is sample entropy (Lebedev et al., 2016). Given that two sequences (e.g. BOLD timeseries) of length *m* are separated by a distance of less than *r* (distance may be measured by Pearson correlation or some other metric), sample entropy quantifies the probability that the sequences will still be less than *r* units apart when they are extended by one sample, i.e. when their lengths are increased to *m* + 1. Lebedev et al. (2016) found a distributed pattern of LSD-induced increases in sample entropy across both unimodal and transmodal cortices.

As discussed in *Section 3.2.3*, the Entropic Brain Hypothesis claims that increasing entropy brings the brain closer to criticality. Three studies (Atasoy et al., 2017; Lord et al., 2019; Ruffini et al., 2023) have measured the effect of psychedelics on criticality. However, the relationship between entropy and criticality is far from straightforward. Entropy and criticality are two different statistical phenomena; while entropy describes the amount of variability or disorder in a system, criticality refers to behaviour exhibited when the so-called "order parameter" of a system undergoes a discontinuous "phase transition" as some other ambient property is modulated (Hesse & Gross, 2014). For example, the order parameter of water is its density, which decreases sharply as the temperature of the water (the ambient property) increases beyond 100° C. This results in a transition between the liquid and the gaseous phases of water. The "critical point" of a system can be roughly described as the threshold between order and disorder, as it is characterised in the Entropic Brain Hypothesis, but in complex, non-equilibrium systems like the brain, states above the critical point may not always be more disordered. Canonical models of criticality in physics, such as the Ising model, describe equilibrium systems, in which all microstates of a system are equally likely in the absence of perturbations. However, in the brain, some microstates may intrinsically be more likely than others; for instance, it may be inherently more probable for a neuron to be active than to be quiescent (Beggs & Timme, 2012). This may consequently predispose the system towards macrostates of greater order even past the critical point. Nevertheless, Ruffini et al. (2023) fit fMRI data on psychedelics to an Ising model and found that the fitted model surprisingly resembled empirical data. Unlike the other two studies on criticality under psychedelics, Ruffini et al. (2023) showed that psychedelics bring the brain away from criticality, since in the sober state, the brain is already above the critical point according to their model.

Critical systems are also characterised by power-law scaling, in which the size, duration, and other parameters $L$ of events $k$ are related by the power law $L(k) \propto k^{-\beta}$, where β is a constant corresponding to the "critical exponent" of the system. To show that psychedelics push the brain towards criticality, one study in the literature showed that psychedelics improve the goodness of fit of brain activity to a power-law relationship (Atasoy et al., 2017). Brain activity was defined here in a unique manner: the "harmonics" of the connectome, or patterns of oscillations determined by the topology of structural connections in the brain. However, the technique used by the study – linear regression – has been shown to yield inaccurate estimates of power-law distributions, which usually disobey the assumptions of linear regression (Clauset et al., 2009).

The third study on criticality (Lord et al., 2019) does not assess power-law relationships in the brain, but instead measures the metastability of brain activity. Metastability is closely related to criticality; it describes a system in which states are locally stable yet are vulnerable to perturbations, resulting in frequent transitions between states. In particular, a metastable system that fluctuates between states of synchronisation and desynchronisation is poised at criticality (Shanahan, 2010). These fluctuations are typically measured with the Kuramoto order parameter, which quantifies the degree of synchrony between the phases of oscillators (Shanahan, 2010). The higher the variance of the Kuramoto order parameter, the more metastable the brain will be, since the oscillating BOLD timeseries will more strongly vary between states of synchronisation and desynchronisation.

What can we take away from this review, given that the entropy measures are so different? We argue that the aspect of the data that entropy is computed on – that is, the variable of interest – matters just as much as, if not more than, the measure of entropy that is used. No two studies in the literature estimated Shannon entropy on the same random variable, which ranged from graph-theoretic properties to the variance of intra-network synchrony. Similarly, none of the studies computed criticality on the same property of brain activity/structure. In this literature, the results were always consistent – psychedelics increased entropy – despite the huge variation in the variable of interest. However, the statement that psychedelics increase entropy does not specify very much, as it does not indicate *which* feature of brain activity is undergoing these changes.

Another problem is that ten of the twelve studies measured entropy on the same two primary datasets (Carhart-Harris, Erritzoe, et al., 2012; Carhart-Harris et al., 2016), which could explain why they all arrived at similar conclusions despite differences in methods. A recent study applied all twelve of the entropy measures in the literature to an independent cohort of 28 participants who were administered psilocybin (McCulloch et al., 2023). (The study is still a pre-print, so we did not officially include it in our review.) Psilocybin did not have a significant effect on seven of these measures, suggesting that the results from the literature may not generalise to other datasets. Additionally, the measures did not correlate well with each other. The study highlights the need for more primary data and replication studies in the field of psychedelic neuroimaging.

##### *Section S4.2. Relaxed Beliefs Under Psychedelics*

A subsequent theory called Relaxed Beliefs Under Psychedelics (REBUS) integrated the Entropic Brain Hypothesis with the Bayesian brain hypothesis (Carhart-Harris & Friston, 2019). Broadly speaking, the Bayesian brain framework states that the brain generates models that make top-down predictions about the world based on certain priors (or "assumptions"). When those predictions are proven wrong by bottom-up information from the environment, the model updates its priors (Holmes

& Nolte, 2019; Knill & Pouget, 2004; Seth & Friston, 2016). REBUS hypothesises that psychedelics relax the precision weighting of the priors, or, in other words, the confidence in the assumptions that shape the model. In other words, psychedelics loosen the suppressive influence of top-down predictions on the processing of bottom-up sensory information, such that strongly held beliefs will exert less control over the interpretation of "sense-data" from the environment. A similar idea called "neural annealing" compares psychedelics to metallurgy; in the same way that heating and then cooling down a metal enables it to recrystallise into new patterns, psychedelics also energise the brain in such a way that it can subsequently self-organise into new equilibria (Gómez-Emilsson, 2021; Johnson, 2019). Under REBUS, these new equilibria would correspond to novel beliefs.

REBUS make several concrete predictions about the effects of psychedelics on the brain, including some that can be tested with fMRI. Hypothesis **(1)**: Psychedelics flatten the functional hierarchy of the brain by increasing bottom-up and decreasing top-down information flow. (There are other hypotheses concerned with changes in connectivity within and between resting-state networks, but we already addressed these in previous sections.) Hypothesis **(2)**: Psychedelics disrupt the integrity of large-scale brain networks, especially those that are situated at the highest layers of the functional hierarchy, such as the DMN. Hypothesis **(3)**: Psychedelics break down the modular structure of the brain, resulting in greater global connectivity (Carhart-Harris & Friston, 2019).

In the literature, some support for hypothesis (1) comes from Girn et al. (2022), which found that psychedelics flatten the principal gradient of functional connectivity from unimodal (i.e. sensory) to transmodal (i.e. associative) cortices. In other words, unimodal regions are less functionally segregated from transmodal regions, suggesting that top-down and bottom-up information flow become less distinguishable from each other. On the other hand, unimodal regions become more segregated from each other. Our meta-analysis of the FC dataset substantiates Girn's findings; it revealed that psychedelics elevate connectivity between unimodal and transmodal networks, while reducing connectivity between unimodal networks. However, because our quantitative meta-analysis only included studies on undirected FC (which constituted the majority of the literature), our results are unable to speak to the direction of information flow under psychedelics. In other words, it is unclear from our results whether psychedelics make information flow more top-down or bottom-up. The evidence for hypothesis (1) from studies on (directed) effective connectivity is more conclusive. For instance, Preller et al. (2019) models the EC of the cortico-striato-thalamo-cortical (CTSC) loop under LSD. If the orientation of information flow is defined by the direction of the loop, then LSD heightens bottom-up information flow, e.g. EC from the thalamus up to the cortex.

Our meta-analysis of the FC data generally upholds hypothesis (2); we demonstrated that psychedelics significantly decrease within-network connectivity in some transmodal networks, including the DMN. However, we did find a significant, albeit small, increase in within-network FC in another transmodal network, the FPN. Our quantitative meta-analysis showed that psychedelics generally elevate between-network connectivity, lending some support to hypothesis (3). Additionally, as discussed in our qualitative review (*Section S5*), five independent studies concluded that psychedelics enhance global brain connectivity (Madsen et al., 2021; Müller, Lenz, Dolder, Lang, et al., 2017; Preller, Burt, et al., 2018; Preller et al., 2020; Tagliazucchi et al., 2016), which is measured as the average correlation between voxels and the whole-brain timeseries. Two graph-theoretic analyses directly revealed that psychedelics diminish the modularity of the brain, implying reduced connectivity within modules and heightened connectivity between modules (Barrett, Krimmel, et al., 2020; Tagliazucchi et al., 2016).

### Section S5. Qualitative review of other connectivity studies

The algorithm that we developed to analyse the FC literature could only incorporate data from studies on pairwise FC. There are other measures in the literature that do not compute pairwise FC, such as global brain connectivity (GBC), which defines the FC between voxels and the whole-brain timeseries, or graph-theoretic metrics (e.g. modularity), which quantify global, structural properties of FC. Additionally, our algorithm only analyses undirected connectivity in the brain, so it cannot accommodate studies of effective connectivity.

In this subsection, we will first summarise the findings of the studies that *did* measure pairwise, undirected connectivity but either did not make their data available or were excluded from the meta-analysis in order to mitigate bias. Then, we will discuss the studies that did not compute pairwise, undirected connectivity.

#### *Section S5.1. Other studies on pairwise, undirected connectivity*

Three of these studies would have been included in the meta-analysis had they provided the necessary data, i.e., coordinates or spatial maps of the regions whose connectivity significantly changed under psychedelics. Carhart-Harris et al. (2013) used ICA to identify 11 RSNs, including the anterior DMN (aDMN). Through linear regression, they calculated the FC between the aDMN and each other RSN. They discovered that psilocybin increased FC between the aDMN and the salience network, right frontoparietal network (FPN), the auditory network, and the dorsal attention network (DAN). They also performed a seed-to-voxel analysis in which they measured the FC between a seed in the ventromedial prefrontal cortex (vmPFC) and every other voxel in the brain. They defined the set of voxels that had positive correlations with the vmPFC as the DMN and the negatively correlated voxels as the task-positive network (TPN). Linear regression revealed that the FC between the DMN and the TPN grew larger under psilocybin.

Roseman et al. (2016) identified patches in the visual cortex (specifically, V1 and V3) that were "retinotopically specific," that is, patches that were uniquely responsive to stimuli presented along the horizontal or vertical meridians. Then, through linear regression, they determined the "retinotopic coordination" under LSD: the difference between the FC of patches with congruent retinotopic specificity (e.g. horizontally-activated patches in V1 and horizontally-activated patches in V3) and the FC of patches with incongruent retinotopic specificity (e.g. horizontally-activated patches in V1 and vertically-activated patches in V3). Retinotopic coordination was greater under LSD than under placebo.

Finally, Barrett, Doss, et al. (2020) computed the Pearson correlation coefficient between the resting-state timeseries of each ROI in the 268-node Shen parcellation (Shen et al., 2013) and found that FC increased overall under psilocybin, although changes in FC did not exhibit any discernible patterns at the network level.

In our meta-analysis, we showed that psychedelics increased FC between the DMN & both the dorsal attention network (VAN) and ventral attention network (VAN), two networks that are commonly associated with the TPN (Seeley et al., 2007), therefore corroborating the findings of Carhart-Harris et al. (2013). The meta-analysis does not examine any data about retinotopic coordination; we found that FC within the visual network decreased under psychedelics, but there is likely no direct correspondence between retinotopic coordination and FC within the visual network. In agreement with Barrett, Doss, et al. (2020), we demonstrated that psilocybin, as well as other psychedelics, generally increased FC.

Our results were generally consistent with the findings of the other eight studies that would have been eligible for the meta-analysis. Of the five that focused on cortical FC, two measured FC with ICA and reported widespread disintegration and desegregation, i.e. reduced within-network FC and elevated between-network FC (Carhart-Harris et al., 2016; Müller et al., 2018). We also showed significant desegregation for most pairs of non-identical brain networks and significant disintegration for all networks except the FPN and SMN. Lebedev et al. (2015) reported a correlation between ego death and salience network disintegration on psilocybin; according to our meta-analysis, the VAN, which is often referred to interchangeably with the salience network, became less connected to itself under psychedelics. In agreement with our findings, Luppi et al. (2021) determined that time-averaged FC globally increased under LSD. (Finally, Mason et al. (2021) reported a nearly identical analysis to the one in Mason et al. (2020), which was included in our quantitative meta-analysis.)

##### *Section S5.2. Studies on non-pairwise or directed connectivity*

The remaining studies used the following methods: graph-theoretic analyses (7 studies), global brain connectivity (6), dynamic causal modelling (4), gradient-based analyses (2), Leading Eigenvector Dynamical Analysis (2), Perturbational Integration Latency Index (1), and local correlation (1).

Seven studies measure the effects of psychedelics on FC with graph-theoretic metrics. A graph is a network structure consisting of nodes and the edges connecting them. When applied to neuroscience, graphs typically describe patterns of FC, in which nodes are ROIs and edges represent the strength (or the mere presence) of connection between ROIs. There are five graph-theoretic properties or structures that are measured in the literature.

The first and most common one is the participation coefficient, which is used in four of the seven studies. The participation coefficient algorithm first divides the brain into modules, which are more connected within themselves than they are with regions outside themselves (Rubinov & Sporns, 2010). The participation coefficient then quantifies the extent to which regions are more connected with other regions in the same module than they are with regions in other modules. Two studies find that psilocybin and LSD reduce modularity in the FPN and in frontal and midline regions, respectively (Barrett, Krimmel, et al., 2020; Tagliazucchi et al., 2016). The third study divided windows of the BOLD timeseries into predominantly integrated and segregated sub-states, as determined by the degree of modularity in each window (Luppi et al., 2021). Intriguingly, the study only revealed significant differences under LSD in the *segregated* rather than the integrated sub-state. Lebedev et al. (2015) quantified the diversity of connections between brain regions and modules, which, in the anterior parahippocampal cortex, negatively correlated with subjective ratings of ego death on LSD.

(Note that this metric of diversity is actually a measure of Shannon entropy, as discussed in *Section 2.2.3.*)

Lebedev et al. (2015) also measured the effect of LSD on another graph-theoretic measure known as the within-network clustering coefficient. The within-network clustering coefficient reflects the proportion of fully connected triplets (sets of three nodes in which each node is connected to both of the others) in the set of all possible triplets in a network (Masuda et al., 2018). The study showed that the decrease in the clustering coefficient of the salience network under LSD correlated with the subjective experience of ego death. On the other hand, Viol et al. (2017) discovered an increase in clustering coefficients under ayahuasca; the divergent findings may be attributable to differences in the thresholding techniques that were applied to define edges.

Petri et al. (2014) identified persistence homological scaffolds in the graph structure of the partial correlation matrix of the brain. These scaffolds (Edelsbrunner et al., 2002) consist of cycles or "holes" that appear and disappear in the graph as edges are added one-by-one; additionally, cycles are weighted based on their persistence, or the length of time between their appearance and disappearance. Under psilocybin, edges in the graph participate in much more persistent cycles, suggesting more stable connectivity between different cortical regions. Furthermore, the homological scaffolds revealed that connections between modules became stronger and much more pervasive.

Viol et al. (2017) measured changes in several graph-theoretic metrics under ayahuasca, including the Shannon entropy of the node degrees, geodesic distance, global and local efficiency, and clustering coefficients. The degree refers to the number of other nodes to which a particular node is connected, geodesic distance to the shortest path between two nodes, and efficiency to the inverse of the harmonic mean of geodesic distances, either across the entire graph (global) or within subgraphs (local). Shannon entropy of node degree, geodesic distance, and local efficiency all grew larger under ayahuasca, whereas global efficiency grew smaller. (Viol et al. (2019) subsequently found that ayahuasca also elevates the Shannon entropy of geodesic distance.) These results suggest that regions within individual subnetworks become more directly connected on ayahuasca, whereas regions in separate subnetworks become less directly connected. This appears to contradict the aforementioned findings of decreased modularity under psilocybin.

To the extent that decreases in modularity correspond to increases in between-network FC, the results of our meta-analysis do agree with the findings of diminished modularity in the literature. In agreement with the findings of decreased clustering coefficients within the salience network in Lebedev et al. (2015), the meta-analysis revealed that the within-network connectivity of the VAN, which is often equated with the salience network, decreases under psychedelics. It is difficult to relate our findings of increased between-network FC and decreased within-network FC on ayahuasca/DMT to the results of Viol et al. (2017, 2019), since changes in FC do not necessarily imply alterations to the geodesic distances between nodes. We cannot know for sure whether our meta-analysis substantiates or negates any of the above findings until we conduct a graph-theoretic analysis on our results, which we leave as a task for a future project.

The six studies on global brain connectivity (GBC), also referred to in the literature as "global correlation" (GCOR), measured the averaged Fisher-transformed Pearson correlation coefficient either between each voxel and a 64-component whole-brain time course (Madsen et al., 2021; Müller, Lenz, Dolder, Lang, et al., 2017) or between each grayordinate and every other grayordinate in the brain (Preller, Burt, et al., 2018; Preller et al., 2020; Tagliazucchi et al., 2016). (A grayordinate is a gray matter coordinate that is specified by a surface vertex in Connectome Workbench.) Madsen et al. (2021) demonstrated increases in GCOR in regions within the DAN and ECN under psilocybin, while Müller, Lenz, Dolder, Lang, et al. (2017) found that LSD amplified GCOR in subcortical areas spanning the thalamus, caudate, putamen, and pallidum. According to Tagliazucchi et al. (2016), LSD heightened GBC in association cortices. Preller, Burt, et al. (2018) reported opposite findings: if GSR was applied, GBC was higher in sensory and somatomotor areas under LSD, yet lower in subcortical areas and regions involved in associative networks. Preller et al. (2020) reported similar results for psilocybin. Finally, Delli Pizzi et al. (2023) measured the intrinsic connectivity contrast (ICC), which is the root-mean square of GBC, and correlated the results with maps of receptor density. They determined that ICC increased in areas rich in 5-$HT_{2A}$ receptors and poor in 5-$HT_{1A}$ receptors.

In agreement with Madsen et al. (2021), we showed that between-network connectivity of the FPN (which is often called the ECN) and DAN both grew larger under psychedelics. While we did not report our analysis on subcortical FC, we also determined that psychedelics elevated FC between subcortical regions, as defined in the Tian atlas (Tian et al., 2020), and between subcortical & cortical networks. This finding agrees with that of Müller, Lenz, Dolder, Lang, et al. (2017). Unlike in Preller, Burt, et al. (2018) and in Preller et al. (2020), associative areas (the VAN, DAN, FPN, and DMN) become more connected to other networks in our analysis, though we do find that the visual network and SMN increase in connectivity with other networks. That being said, while their results may overlap, pairwise brain-network analysis cannot be equated with global brain connectivity analysis.

The results of Delli Pizzi et al. (2023) are especially pertinent to this meta-analysis because they sought to integrate the neuroimaging and pharmacology of psychedelics. According to existing PET maps (Hansen et al., 2022), the 5-$HT_{1A}$ receptor is expressed most in the DAN, whereas the 5-$HT_{2A}$ receptor is densest in the VAN and FPN. Our meta-analysis lends some support to Delli Pizzi et al. (2023), as we found that the total increase in FC was greater for the VAN and FPN than for the DAN, although only slightly.

Studies on dynamic causal modelling (DCM) (Bedford et al., 2023; Kaelen et al., 2016; Kraehenmann, Schmidt, et al., 2015; Preller et al., 2019), unlike all the others, measure effective connectivity (EC) rather than functional connectivity. FC is fundamentally different from EC; the former analyses correlations between the activity of brain regions, whereas the latter models the simplest brain circuit that explains the data (Stephan & Friston, 2010). The circuit includes intrinsic connections (between an ROI and itself), bilinear extrinsic connections (between ROIs), the driving inputs that affect the intrinsic connections, and the modulatory effects of the experiment on the strength of the extrinsic connections (Friston et al., 2003). Once they have been specified, these causal factors entail a model space, in which each model is defined by a particular set of driving inputs and modulatory effects. Bayesian model selection (BMS) is then used to identify the model that best explains the data.

Kaelen et al. (2016) measured brain activity in response to music while participants were on LSD. The ROIs in the DCM of that study are the parahippocampus (PH), which is involved in regulating emotional responses to music (Baumgartner et al., 2006; Gosselin et al., 2006; Koelsch, 2014), and the visual cortex (VC), which is activated when the PH is stimulated externally (Barbeau et al., 2005). BMS revealed that, in the model with the highest evidence, music and LSD weakened the intrinsic connections of the two ROIs, whereas the interaction effect between them positively modulated the connection from the PH to the VC.

Kraehenmann, Schmidt, et al. (2015) sought to determine the changes in connectivity that mediate the amygdala's decreased response to emotionally charged images under psilocybin. Their ROIs are the amygdala; visual cortex 1 (V1), which is driven by the images; and the lateral prefrontal cortex (lPFC), which regulates the amygdala's reaction to emotionally salient stimuli (Aznar & Klein, 2013; Hahn et al., 2011). They found that, in the optimal model, threatening stimuli modulated both bottom-up (V1 → amygdala → lPFC) and top-down (lPFC → amygdala → V1) connections, and furthermore that the interaction between threatening stimuli and psilocybin negatively modulated the connectivity from the amygdala to V1.

Preller et al. (2019) ran a spectral DCM (Friston et al., 2014), which, unlike the DCM used in the other two studies, is advantageous for modelling resting-state fMRI data. In the spectral DCM, the intrinsic and extrinsic connectivity between ROIs is modulated not by the experiment (e.g. tasks or stimuli) but rather by stochastic fluctuations. A Parametric Empirical Bayes (PEB) model then estimates the effect of between-subject differences (in this case, the placebo vs. LSD condition) on within-subject connections. Preller et al. (2019) measured the effect of LSD on the cortico-straiatal-thalamo-cortical (CSTC) loop, which is comprised of ROIs in the PCC, thalamus, ventral striatum (VS), and the superior temporal gyrus (STG). The study found that, relative to placebo, LSD increased connectivity from the thalamus to the PCC from the thalamus to the VS, from the VS to the STG, and from the STG to itself, while decreasing connectivity from the PCC to the thalamus.

Finally, Bedford et al. (2023) applied another type of DCM known as regression DCM (rDCM), which is also useful for modeling resting-state fMRI. Whereas normal DCM requires the *a priori* selection of a handful of brain regions in which to measure connectivity, rDCM enables connectivity estimates across the entire brain by recasting DCM as a linear regression in the frequency domain (Bedford et al., 2023). Connections between individual regions and themselves are assumed to be inhibitory. Bedford et al. (2023) showed that LSD elevated EC between regions and reduced self-inhibition, except in occipital and subcortical regions.

Because the DCM-based studies all have different aims and they do not model the same modulatory effects and driving inputs, it is difficult to perform a meta-analysis on them. Even if their methods were consistent with one another, our meta-analysis algorithm does not incorporate any information about modulatory effects and driving effects. Additionally, the aggregate FC matrix computed in our meta-analysis is symmetric and therefore does not contain any information about the directionality of connections between brain networks, whereas the EC measured in DCMs is inherently directional. In Preller et al. (2019), the sign (positive or negative) of PCC-VS and thalamus-VS connectivity depends on the direction of connectivity, so it is meaningless to compare our undirected results to theirs. However, we can corroborate the findings of Bedford et al. (2023), as we also found that psychedelics elevate between-network FC.

Fundamentally different than FC approaches that assess correlations between discrete ROIs or networks, the gradient-based methods in Girn et al. (2022) and Atasoy et al. (2017) identify axes of connectivity that span multiple regions of the brain. Our meta-analysis algorithm cannot accommodate these studies either, as the algorithm spatially localises patterns of FC to the Yeo networks.

Girn et al. (2022) computed the similarity between the global FC pattern (as measured by the Pearson correlation coefficient) of each of the 10,000 vertices in each cortical surface. They then applied diffusion map embedding to a similarity (covariance) matrix, resulting in a "gradient space"; vertices with lower (higher) gradient values had fewer (more) global connections. Unimodal and transmodal regions of cortex were defined as the set of vertices with lower-percentile and higher-percentile gradient values, respectively. On both LSD and psilocybin, unimodal and transmodal regions became more correlated with each other, whereas within-unimodal and within-transmodal connectivity declined. In the meta-analysis, we also found higher FC between unimodal and transmodal networks under psychedelics, as well as a reduction in FC within and between unimodal networks.

On the other hand, Atasoy et al. (2017) defined gradients of connectivity based on the harmonics of the connectome, which are patterns of coactivation that each have a characteristic spatial frequency and are constrained by the graph structure of white matter tracts in the brain. Each such pattern is associated with a certain amount of power (strength of activation) and energy (strength of activation scaled by the characteristic frequency). LSD elevated the power and energy of the harmonics, especially the energy of high-frequency harmonics; expanded the repertoire of connectome harmonics that contribute to the overall state of brain activity; and reduced cross-frequency correlations, especially among low-frequency harmonics. It is difficult to relate these findings to the results of the meta-analysis due to the spectral content of the data; there is no information about the *frequency* of brain network activation in the meta-analysis.

Leading Eigenvector Dynamical Analysis (LEIDA) identifies the leading eigenvector of the BOLD phase-locking matrix, which reflects a dominant global pattern of connectivity in the brain. Each region's BOLD phase is then projected onto the leading eigenvector. *K*-means clustering on these projections reveal brain states, e.g. resting-state networks, that are recurringly visited over time. Lord et al. (2019) discovered that psilocybin decreased the probability of a phase-locking state that strongly resembled the FPN and increased the probability of a phase-locking state that reflects global coherence. Olsen et al. (2022) showed that psilocybin reduced the occurrence and dwell time of a brain state characterised by connectivity between the lateral frontoparietal (part of the FPN) and medial parietal-cingulate (part of the DMN) cortices. To the extent that phase-locking reflects FC as defined by most other studies in the literature, our meta-analysis support neither of the above findings. According to our results, both within-network FC for the FPN and FPN-DMN FC increased, though between-network FC generally grew larger.

The last two analysis methods were each performed by only one study, so there was not enough data to execute the meta-analysis algorithm on them. (The algorithm would not have been able to accommodate the results from these studies anyway.)

Jobst et al. (2021) computed changes in the Perturbational Integration Latency Index (PILI) under LSD. They used a whole-brain model to perturb the supercritical Hopf bifurcation parameter of brain regions (Deco, Cabral, et al., 2018), resulting in patterns of global synchronisation and desynchronisation. Then, they compared the integration of brain activity, which was quantified as the overall amount of phase-locking, before and after perturbation. Changes in integration therefore served as a measure of the recovery of brain dynamics to the baseline state. Under LSD, brain dynamics, especially in the limbic network, visual network, and DMN, were perturbed farther away from their baseline state and also took longer to recover than under placebo. While there is no corollary to this finding in the results of our meta-analysis, they do substantiate another conclusion in the study, which is that the baseline state of brain dynamics is more integrated under LSD than under placebo. In other words, the brain exhibits greater global FC, as we show through increased between-network FC in the meta-analysis.

The local correlation (LCOR) analysis in Madsen et al. (2021) measures a weighted average of the Pearson correlation coefficient between the timeseries of each voxel and neighbouring voxels. Psilocybin decreased LCOR in regions within the DMN, ECN, and visual cortex. If LCOR can be likened to a measure of within-network connectivity, then our meta-analysis supports these results, although within-network connectivity for the FPN, which is equated with the ECN, does increase.

### Section S6. Sources of heterogeneity and bias

*Section S6.1. Phenomenology*

Heterogeneity indicates that the true effect size is not a single, fixed value but rather exists along a distribution. In our phenomenology meta-analysis, we determined that residual heterogeneity was very high, suggesting that the observed effect sizes deviated strongly from the estimated true effect size. (Unfortunately, we were unable to determine the amount of heterogeneity that was attributed to within-study and between-study variances.) However, we were unable to identify any methodological covariates that significantly contributed to these deviations. The large degree of heterogeneity in the effect sizes may be attributable to the strong inter-individual variability in phenomenological reports. This is an issue that pertains to subjective measurements in general and especially affects psychedelic research, given that these drugs can have widely varying effects on different individuals (Knudsen, 2023; Moujaes et al., 2023).

We also found that most phenomenology studies were either at moderate (52% of studies) or serious (48%) risk of overall bias. While all studies received low ratings of bias in five of the domains, most studies were either moderately or seriously biased in two other domains: confounding and measurement of outcomes. We deemed that studies were confounded by the participants' levels of prior psychedelic use. Compared to those with no experience, participants who have had multiple trips may have a stronger frame of reference for the subjective effects of psychedelics. In other words, they can judge the subjective effects relative to their past experiences. This may or may not have a significant effect on ASC ratings; our own analysis revealed that prior psychedelic use did not significantly confound the pooled effect sizes. Therefore, while none of the studies explicitly adjusted for prior psychedelic use in their analyses, we rated risk of bias from confounding as moderate, so long as the studies reported the amount of prior use. If the latter was not the case, then we rated risk of bias as serious. One study recruited only participants with no prior psychedelic experiences (Hasler et al., 2004), so this was the only study that had low risk of bias in the confounding domain.

One of the most prominent sources of bias in psychedelic research is unblinding, which occurs because participants realise that they received a psychedelic even when the study is intended to be double-blinded (Butler et al., 2022; Goodwin et al., 2023; Wen et al., 2023). In one clinical trial, more than 90% of participants correctly guessed whether they were administered psilocybin or placebo (Bogenschutz et al., 2022). Arguably, unblinding is not an issue in phenomenological studies, where positive ratings of subjective effects require the participant to be aware that they are in an altered state of consciousness. However, as long as the participant knows for certain that they were administered a psychedelic, their confidence in their ratings will increase, which may subsequently influence the ratings themselves. Thus, we attributed at least a moderate risk of bias in measurement of outcome to any study in which it was relatively likely for participants to correctly guess which intervention (drug and dose level, if applicable) they received. Studies that used a within-subjects, cross-over design with only two drugs – psychedelic and inactive placebo – were deemed to have a serious risk of bias. Participants who received a psychedelic in the first session were likely to know with certainty that they would get a placebo in the second session; thus, they may respond that they experienced zero effects along all dimensions of the ASC scale, whereas participants who received placebo in the first session may give low but non-zero responses for some dimensions. The only studies with low risk of bias in measurement of outcome either took extra steps to reduce expectancy effects (Carbonaro et al., 2018), or used experimental conditions that made it difficult for participants to guess the intervention they received; for instance, one study applied either sham or real brain stimulation in addition to administering either psilocybin or placebo (Ort et al., 2023), while another administered three phenomenologically similar psychedelics with no placebo condition (Ley et al., 2023).

Finally, we also found evidence for publication bias in addition to bias in individual studies. In particular, small studies in the phenomenology literature appeared to report disproportionately high effect sizes.

*Section S6.2. Neuroimaging*

Effect sizes in the FC literature were not directly comparable between studies, due to the wide range of parcellations. Therefore, we were unable to apply traditional measures of heterogeneity. However, we did attempt to measure the effect of various methodological covariates, such as preprocessing techniques, on the results of our FC meta-analysis; we did not find a significant effect for any of the six covariates.

For the same reason that we could not measure heterogeneity, we could not quantify publication bias as we did for the phenomenology and pharmacology meta-analyses. However, our qualitative assessment revealed that the most concerning source of publication bias in the neuroimaging literature is the large number of secondary analyses of the same primary datasets. (The reason for the lack of primary data may be attributable to the illegal status of psychedelic drugs in many countries.) Of the 58 studies that we included in the neuroimaging meta-analysis, fewer than half (22) collected original data, and over a third are secondary analyses of the same two primary datasets (Carhart-Harris, Erritzoe, et al., 2012; Carhart-Harris et al., 2016) (**Figure S2a**). Ten of the 12 studies on entropy analyse data from these primary datasets, so it is not surprising that all of them conclude that psychedelics increase entropy, even though all the measures of entropy are different. If the studies had measured entropy on different primary datasets, then the results may have been different. In fact, a recent study applied each of the entropy measures used in the literature to a new, independent cohort of 28 participants and found that psilocybin had no significant effect on seven of the 12 measures (McCulloch et al., 2023). (Note that this study is a pre-print still, so we did not include it in our review). We acknowledge this bias in our qualitative review of the entropy literature, and we also account for it in our quantitative meta-analysis of the functional connectivity (FC) literature, where we only included one study on each unique dataset.

We also performed a risk-of-bias assessment for individual FC studies. Ten of the 12 studies that we included in our quantitative meta-analysis were deemed to have moderate risk of bias, while the remaining two had serious risk of bias. In particular, most studies were judged to be at moderate risk of bias due to confounding. Psychedelics are known to be vasoconstrictors (Dyer & Gant, 1973) which may influence the fMRI signal by affecting the concentration of oxygenated hemoglobin (Hillman, 2014; Özbay et al., 2018, 2019). None of the studies explicitly controlled for vasoconstriction, but it is also unclear how to feasibly and directly measure neural vasoconstriction in a study with humans, so most studies were assigned a moderate risk-of-bias rating in the confounding domain. Timmermann et al. (2023) came the closest to receiving a low risk-of-bias rating in this domain because it measured EEG in addition to fMRI, and EEG signal is less susceptible to vascular changes than fMRI (Kumral et al., 2020). Studies that did not report the results of head motion analyses, such as the number of participants excluded due to excessive motion or the presence of significant between-group differences in head motion, were given a serious risk-of-bias rating in the confounding domain; head motion is known to strongly confound fMRI measurements (Makowski et al., 2019). One study that used a between-subjects, parallel group design received a moderate risk-of-bias rating in the missing data domain, since more subjects were excluded in the psilocybin group than in the placebo group for head motion (Mason et al., 2020). Studies that were not double-blind were deemed to have moderate risk of bias due to measurement of outcomes. In this case, the researchers were aware of whether participants received a psychedelic or placebo, but this likely did not have a significant effect on the measurement outcomes.

*Section S6.3. Pharmacology*

We observed very high levels of heterogeneity for both the affinity and functional activity literature. For the affinity studies, we were able to identify reasons for this heterogeneity: the choice of radioligand significantly influenced our pooled estimates of selectivity for 5-$HT_{2A}$ and 5-$HT_{2C}$. This is not surprising, given that differences in radioligand properties, including selectivity, the degree of cooperativity between the radioligand and ligand of interest, and association and dissociation rates, are known to influence estimates of $K_i$ values (Bosma et al., 2019; Hulme & Trevethick, 2010). Unfortunately, the variability in radioligands was too high for us to conduct a separate meta-analysis for each radioligand type.

Furthermore, affinity and activity are fundamentally linked quantities, for two reasons (Strange, 2008). Firstly, since serotonin and dopamine receptors are both GPCRs, GPCR agonists, such as psychedelics, bind not only to the receptor but also to G proteins; hence, measurements of affinity will be biased by the extent to which the agonist activates G proteins (De Lean et al., 1980). Secondly, binding induces a conformational change to the receptor, which in turn stabilises a partially active state of the receptor without even coupling to G proteins (Colquhoun, 1998; Strange, 1999).

Hence, in addition to performing a meta-analysis on affinity, we also analysed the functional activity of psychedelics. We sought a measure of activity that was minimally biased: one that was not very susceptible to confounding influences. Classical receptor theory indicates that activity ratios ($\Delta EC_{50}$, where $EC_{50}$ is the concentration of drug that is needed to bring about 50% of the maximal response) are unbiased by experimental conditions, such as the density of the receptor of interest in the corresponding assay (Kenakin et al., 2012). However, when a drug is a partial, rather than full, agonist at a receptor, there is a nonlinear relationship between receptor density and activity ratios (Kenakin, 2017). One solution is to compare the relative activity, defined as $\Delta\log(E_{max}/EC_{50})$ (Kenakin et al., 2011), where $E_{max}$ is the maximal response elicited by the drug of interest relative to a reference ligand. Normalising the activity ratio by $E_{max}$ essentially "cancels out" the effect of receptor density, as well as the sensitivity of the assay in general, on activity estimates, resulting in a minimally biased measure of receptor agonism (Kenakin, 2017).

### Supplementary References

Akaike, H. (1998). Information Theory and an Extension of the Maximum Likelihood Principle. In E. Parzen, K. Tanabe, & G. Kitagawa (Eds.), *Selected Papers of Hirotugu Akaike* (pp. 199–213). Springer. https://doi.org/10.1007/978-1-4612-1694-0_15

Assink, M., & Wibbelink, C. (2016). Fitting three-level meta-analytic models in R: A step-by-step tutorial. *The Quantitative Methods for Psychology*, *12*. https://doi.org/10.20982/tqmp.12.3.p154

Atasoy, S., Roseman, L., Kaelen, M., Kringelbach, M. L., Deco, G., & Carhart-Harris, R. L. (2017). Connectome-harmonic decomposition of human brain activity reveals dynamical repertoire re-organization under LSD. *Scientific Reports*, *7*(1), Article 1. https://doi.org/10.1038/s41598-017-17546-0

Avram, M., Müller, F., Rogg, H., Korda, A., Andreou, C., Holze, F., Vizeli, P., Ley, L., Liechti, M. E., & Borgwardt, S. (2022). Characterizing Thalamocortical (Dys)connectivity Following D-Amphetamine, LSD, and MDMA Administration. *Biological Psychiatry. Cognitive Neuroscience and Neuroimaging*, *7*(9), 885–894. https://doi.org/10.1016/j.bpsc.2022.04.003

Aznar, S., & Klein, A. B. (2013). Regulating prefrontal cortex activation: An emerging role for the 5-$HT_2A$ serotonin receptor in the modulation of emotion-based actions? *Molecular Neurobiology*, *48*(3), 841–853. https://doi.org/10.1007/s12035-013-8472-0

Ballentine, G., Friedman, S. F., & Bzdok, D. (2022). Trips and neurotransmitters: Discovering principled patterns across 6850 hallucinogenic experiences. *Science Advances*, *8*(11), eabl6989. https://doi.org/10.1126/sciadv.abl6989

Barbeau, E., Wendling, F., Régis, J., Duncan, R., Poncet, M., Chauvel, P., & Bartolomei, F. (2005). Recollection of vivid memories after perirhinal region stimulations: Synchronization in the theta range of spatially distributed brain areas. *Neuropsychologia*, *43*(9), 1329–1337. https://doi.org/10.1016/j.neuropsychologia.2004.11.025

Barrett, F. S., Doss, M. K., Sepeda, N. D., Pekar, J. J., & Griffiths, R. R. (2020). Emotions and brain function are altered up to one month after a single high dose of psilocybin. *Scientific Reports*, *10*(1), 2214. https://doi.org/10.1038/s41598-020-59282-y

Barrett, F. S., Krimmel, S. R., Griffiths, R. R., Seminowicz, D. A., & Mathur, B. N. (2020). Psilocybin acutely alters the functional connectivity of the claustrum with brain networks that support perception, memory, and attention. *NeuroImage*, *218*, 116980. https://doi.org/10.1016/j.neuroimage.2020.116980

Barrett, F. S., Preller, K. H., Herdener, M., Janata, P., & Vollenweider, F. X. (2018). Serotonin 2A Receptor Signaling Underlies LSD-induced Alteration of the Neural Response to Dynamic Changes in Music. *Cerebral Cortex (New York, N.Y.: 1991)*, *28*(11), 3939–3950. https://doi.org/10.1093/cercor/bhx257

Barttfeld, P., Uhrig, L., Sitt, J. D., Sigman, M., Jarraya, B., & Dehaene, S. (2015). Signature of consciousness in the dynamics of resting-state brain activity. *Proceedings of the National Academy of Sciences*, *112*(3), 887–892. https://doi.org/10.1073/pnas.1418031112

Baumgartner, T., Lutz, K., Schmidt, C. F., & Jäncke, L. (2006). The emotional power of music: How music enhances the feeling of affective pictures. *Brain Research*, *1075*(1), 151–164. https://doi.org/10.1016/j.brainres.2005.12.065

Becker, A. M., Klaiber, A., Holze, F., Istampoulouoglou, I., Duthaler, U., Varghese, N., Eckert, A., & Liechti, M. E. (2023). Ketanserin Reverses the Acute Response to LSD in a Randomized, Double-Blind, Placebo-Controlled, Crossover Study in Healthy Participants. *International Journal of Neuropsychopharmacology*, *26*(2), 97–106. https://doi.org/10.1093/ijnp/pyac075

Bedford, P., Hauke, D. J., Wang, Z., Roth, V., Nagy-Huber, M., Holze, F., Ley, L., Vizeli, P., Liechti, M. E., Borgwardt, S., Müller, F., & Diaconescu, A. O. (2023). The effect of lysergic acid diethylamide (LSD) on whole-brain functional and effective connectivity. *Neuropsychopharmacology*, *48*(8), Article 8. https://doi.org/10.1038/s41386-023-01574-8

Beliveau, V., Ganz, M., Feng, L., Ozenne, B., Højgaard, L., Fisher, P. M., Svarer, C., Greve, D. N., & Knudsen, G. M. (2017). A High-Resolution In Vivo Atlas of the Human Brain's Serotonin System. *The Journal of Neuroscience: The Official Journal of the Society for Neuroscience*, *37*(1), 120–128. https://doi.org/10.1523/JNEUROSCI.2830-16.2016

Bernasconi, F., Schmidt, A., Pokorny, T., Kometer, M., Seifritz, E., & Vollenweider, F. X. (2014). Spatiotemporal brain dynamics of emotional face processing modulations induced by the serotonin 1A/2A receptor agonist psilocybin. *Cerebral Cortex (New York, N.Y.: 1991)*, *24*(12), 3221–3231. https://doi.org/10.1093/cercor/bht178

Bernhard, M. K., & Ulrich, K. (2009). [Recurrent cortical blindness after LSD-intake]. *Fortschritte Der Neurologie-Psychiatrie*, *77*(2), 102–104. https://doi.org/10.1055/s-0028-1109114

Bershad, A. K., Preller, K. H., Lee, R., Keedy, S., Wren-Jarvis, J., Bremmer, M. P., & de Wit, H. (2020). Preliminary Report on the Effects of a Low Dose of LSD on Resting-State Amygdala Functional Connectivity. *Biological Psychiatry. Cognitive Neuroscience and Neuroimaging*, *5*(4), 461–467. https://doi.org/10.1016/j.bpsc.2019.12.007

Blair, J. B., Marona-Lewicka, D., Kanthasamy, A., Lucaites, V. L., Nelson, D. L., & Nichols, D. E. (1999). Thieno[3,2-b]- and thieno[2,3-b]pyrrole bioisosteric analogues of the hallucinogen and serotonin agonist N,N-dimethyltryptamine. *Journal of Medicinal Chemistry*, *42*(6), 1106–1111. https://doi.org/10.1021/jm980692q

Bogenschutz, M. P., Ross, S., Bhatt, S., Baron, T., Forcehimes, A. A., Laska, E., Mennenga, S. E., O'Donnell, K., Owens, L. T., Podrebarac, S., Rotrosen, J., Tonigan, J. S., & Worth, L. (2022). Percentage of Heavy Drinking Days Following Psilocybin-Assisted Psychotherapy vs Placebo in the Treatment of Adult Patients With Alcohol Use Disorder: A Randomized Clinical Trial. *JAMA Psychiatry*, *79*(10), 953–962. https://doi.org/10.1001/jamapsychiatry.2022.2096

Boly, M., Seth, A. K., Wilke, M., Ingmundson, P., Baars, B., Laureys, S., Edelman, D. B., & Tsuchiya, N. (2013). Consciousness in humans and non-human animals: Recent advances and future directions. *Frontiers in Psychology*, *4*, 625. https://doi.org/10.3389/fpsyg.2013.00625

Borenstein, M., Hedges, L. V., Higgins, J. P. T., & Rothstein, H. R. (2009). Publication Bias. In *Introduction to Meta-Analysis* (pp. 277–292). John Wiley & Sons, Ltd. https://doi.org/10.1002/9780470743386.ch30

Bosma, R., Stoddart, L. A., Georgi, V., Bouzo-Lorenzo, M., Bushby, N., Inkoom, L., Waring, M. J., Briddon, S. J., Vischer, H. F., Sheppard, R. J., Fernández-Montalván, A., Hill, S. J., & Leurs, R. (2019). Probe dependency in the determination of ligand binding kinetics at a prototypical G protein-coupled receptor. *Scientific Reports*, *9*(1), Article 1. https://doi.org/10.1038/s41598-019-44025-5

Braden, M. R., & Nichols, D. E. (2007). Assessment of the roles of serines 5.43(239) and 5.46(242) for binding and potency of agonist ligands at the human serotonin 5-HT2A receptor. *Molecular Pharmacology*, *72*(5), 1200–1209. https://doi.org/10.1124/mol.107.039255

Brett, M., Anton, J.-L., Valabregue, R., & Poline, J.-B. (2002). Region of Interest Analysis Using an SPM Toolbox [Abstract]. *Neuroimage*, *16*. https://doi.org/10.1016/S1053-8119(02)90013-3

Burt, J. B., Preller, K. H., Demirtas, M., Ji, J. L., Krystal, J. H., Vollenweider, F. X., Anticevic, A., & Murray, J. D. (2021). Transcriptomics-informed large-scale cortical model captures topography of pharmacological neuroimaging effects of LSD. *eLife*, *10*, e69320. https://doi.org/10.7554/eLife.69320

Bushman, B. J., & Wang, M. C. (2009). Vote-counting procedures in meta-analysis. In *The handbook of research synthesis and meta-analysis, 2nd ed* (pp. 207–220). Russell Sage Foundation.

Butcher, T. J., Dzemidzic, M., Harezlak, J., Hulvershorn, L. A., & Oberlin, B. G. (2021). Brain responses during delay discounting in youth at high-risk for substance use disorders. *NeuroImage: Clinical*, *32*, 102772. https://doi.org/10.1016/j.nicl.2021.102772

Butler, M., Jelen, L., & Rucker, J. (2022). Expectancy in placebo-controlled trials of psychedelics: If so, so what? *Psychopharmacology*, *239*(10), 3047–3055. https://doi.org/10.1007/s00213-022-06221-6

Carbonaro, T. M., Johnson, M. W., Hurwitz, E., & Griffiths, R. R. (2018). Double-blind comparison of the two hallucinogens psilocybin and dextromethorphan: Similarities and differences in subjective experiences. *Psychopharmacology*, *235*(2), 521–534. https://doi.org/10.1007/s00213-017-4769-4

Carhart-Harris, R. L., Erritzoe, D., Williams, T., Stone, J. M., Reed, L. J., Colasanti, A., Tyacke, R. J., Leech, R., Malizia, A. L., Murphy, K., Hobden, P., Evans, J., Feilding, A., Wise, R. G., & Nutt, D. J. (2012). Neural correlates of the psychedelic state as determined by fMRI studies with psilocybin. *Proceedings of the National Academy of Sciences*, *109*(6), 2138–2143. https://doi.org/10.1073/pnas.1119598109

Carhart-Harris, R. L., & Friston, K. J. (2019). REBUS and the Anarchic Brain: Toward a Unified Model of the Brain Action of Psychedelics. *Pharmacological Reviews*, *71*(3), 316–344. https://doi.org/10.1124/pr.118.017160

Carhart-Harris, R. L., Leech, R., Erritzoe, D., Williams, T. M., Stone, J. M., Evans, J., Sharp, D. J., Feilding, A., Wise, R. G., & Nutt, D. J. (2013). Functional connectivity measures after psilocybin inform a novel hypothesis of early psychosis. *Schizophrenia Bulletin*, *39*(6), 1343–1351. https://doi.org/10.1093/schbul/sbs117

Carhart-Harris, R. L., Leech, R., Williams, T. M., Erritzoe, D., Abbasi, N., Bargiotas, T., Hobden, P., Sharp, D. J., Evans, J., Feilding, A., Wise, R. G., & Nutt, D. J. (2012). Implications for psychedelic-assisted psychotherapy: Functional magnetic resonance imaging study with psilocybin. *The British Journal of Psychiatry: The Journal of Mental Science*, *200*(3), 238–244. https://doi.org/10.1192/bjp.bp.111.103309

Carhart-Harris, R. L., Muthukumaraswamy, S., Roseman, L., Kaelen, M., Droog, W., Murphy, K., Tagliazucchi, E., Schenberg, E. E., Nest, T., Orban, C., Leech, R., Williams, L. T., Williams, T. M., Bolstridge, M., Sessa, B., McGonigle, J., Sereno, M. I., Nichols, D., Hellyer, P. J., … Nutt, D. J. (2016). Neural correlates of the LSD experience revealed by multimodal neuroimaging. *Proceedings of the National Academy of Sciences of the United States of America*, *113*(17), 4853–4858. https://doi.org/10.1073/pnas.1518377113

Carhart-Harris, R. L., Roseman, L., Bolstridge, M., Demetriou, L., Pannekoek, J. N., Wall, M. B., Tanner, M., Kaelen, M., McGonigle, J., Murphy, K., Leech, R., Curran, H. V., & Nutt, D. J. (2017). Psilocybin for treatment-resistant depression: fMRI-measured brain mechanisms. *Scientific Reports*, *7*(1), Article 1. https://doi.org/10.1038/s41598-017-13282-7

Carhart-Harris, R., Leech, R., Hellyer, P., Shanahan, M., Feilding, A., Tagliazucchi, E., Chialvo, D., & Nutt, D. (2014). The entropic brain: A theory of conscious states informed by neuroimaging research with psychedelic drugs. *Frontiers in Human Neuroscience*, *8*. https://www.frontiersin.org/article/10.3389/fnhum.2014.00020

Carter, O. L., Burr, D. C., Pettigrew, J. D., Wallis, G. M., Hasler, F., & Vollenweider, F. X. (2005). Using psilocybin to investigate the relationship between attention, working memory, and the serotonin 1A and 2A receptors. *Journal of Cognitive Neuroscience*, *17*(10), 1497–1508. https://doi.org/10.1162/089892905774597191

Carter, O. L., Hasler, F., Pettigrew, J. D., Wallis, G. M., Liu, G. B., & Vollenweider, F. X. (2007). Psilocybin links binocular rivalry switch rate to attention and subjective arousal levels in humans. *Psychopharmacology*, *195*(3), 415–424. https://doi.org/10.1007/s00213-007-0930-9

Carter, O. L., Pettigrew, J. D., Hasler, F., Wallis, G. M., Liu, G. B., Hell, D., & Vollenweider, F. X. (2005). Modulating the rate and rhythmicity of perceptual rivalry alternations with the mixed 5-HT2A and 5-HT1A agonist psilocybin. *Neuropsychopharmacology: Official Publication of the American College of Neuropsychopharmacology*, *30*(6), 1154–1162. https://doi.org/10.1038/sj.npp.1300621

Chadeayne, A. R., Pham, D. N. K., Reid, B. G., Golen, J. A., & Manke, D. R. (2020). Active Metabolite of Aeruginascin (4-Hydroxy-N,N,N-trimethyltryptamine): Synthesis, Structure, and Serotonergic Binding Affinity. *ACS Omega*, *5*(27), 16940–16943. https://doi.org/10.1021/acsomega.0c02208

Cheung, M. W.-L. (2014). Modeling dependent effect sizes with three-level meta-analyses: A structural equation modeling approach. *Psychological Methods*, *19*(2), 211–229. https://doi.org/10.1037/a0032968

Cimadevila, M., Gómez-García, L., Martínez, A. L., Iglesias, A., López-Giménez, J., Castro, M., Cadavid, M. I., Loza, M. I., & Brea, J. (2020). Essential role of the C148-C227 disulphide bridge in the human 5-HT2A homodimeric receptor. *Biochemical Pharmacology*, *177*, 113985. https://doi.org/10.1016/j.bcp.2020.113985

Clauset, A., Shalizi, C. R., & Newman, M. E. J. (2009). Power-Law Distributions in Empirical Data. *SIAM Review*, *51*(4), 661–703. https://doi.org/10.1137/070710111

Colquhoun, D. (1998). Binding, gating, affinity and efficacy: The interpretation of structure-activity relationships for agonists and of the effects of mutating receptors. *British Journal of Pharmacology*, *125*(5), 923–947. https://doi.org/10.1038/sj.bjp.0702164

Coyle, J. R., Presti, D. E., & Baggott, M. J. (2012). *Quantitative Analysis of Narrative Reports of Psychedelic Drugs* (arXiv:1206.0312). arXiv. https://doi.org/10.48550/arXiv.1206.0312

Cussac, D., Boutet-Robinet, E., Ailhaud, M.-C., Newman-Tancredi, A., Martel, J.-C., Danty, N., & Rauly-Lestienne, I. (2008). Agonist-directed trafficking of signalling at serotonin 5-HT2A, 5-HT2B and 5-HT2C-VSV receptors mediated Gq/11 activation and calcium mobilisation in CHO cells. *European Journal of Pharmacology*, *594*(1–3), 32–38. https://doi.org/10.1016/j.ejphar.2008.07.040

Dai, R., Larkin, T. E., Huang, Z., Tarnal, V., Picton, P., Vlisides, P. E., Janke, E., McKinney, A., Hudetz, A. G., Harris, R. E., & Mashour, G. A. (2023). Classical and non-classical psychedelic drugs induce common network changes in human cortex. *NeuroImage*, *273*, 120097. https://doi.org/10.1016/j.neuroimage.2023.120097

Daumann, J., Heekeren, K., Neukirch, A., Thiel, C. M., Möller-Hartmann, W., & Gouzoulis-Mayfrank, E. (2008). Pharmacological modulation of the neural basis underlying inhibition of return (IOR) in the human 5-HT2A agonist and NMDA antagonist model of psychosis. *Psychopharmacology*, *200*(4), 573–583. https://doi.org/10.1007/s00213-008-1237-1

Daumann, J., Wagner, D., Heekeren, K., Neukirch, A., Thiel, C. M., & Gouzoulis-Mayfrank, E. (2010). Neuronal correlates of visual and auditory alertness in the DMT and ketamine model of psychosis. *Journal of Psychopharmacology (Oxford, England)*, *24*(10), 1515–1524. https://doi.org/10.1177/0269881109103227

Davies, M., Nowotka, M., Papadatos, G., Dedman, N., Gaulton, A., Atkinson, F., Bellis, L., & Overington, J. P. (2015). ChEMBL web services: Streamlining access to drug discovery data and utilities. *Nucleic Acids Research*, *43*(Web Server issue), W612–W620. https://doi.org/10.1093/nar/gkv352

Davis, K. L., Charney, D., Coyle, J. T., & Nemeroff, C. B. (2002). *Neuropsychopharmacology: The Fifth Generation of Progress : An Official Publication of the American College of Neuropsychopharmacology*. Lippincott Williams & Wilkins. http://ebookcentral.proquest.com/lib/oxford/detail.action?docID=3418686

de Araujo, D. B., Ribeiro, S., Cecchi, G. A., Carvalho, F. M., Sanchez, T. A., Pinto, J. P., de Martinis, B. S., Crippa, J. A., Hallak, J. E. C., & Santos, A. C. (2011). Seeing with the eyes shut: Neural basis of enhanced imagery following ayahuasca ingestion. *Human Brain Mapping*, *33*(11), 2550–2560. https://doi.org/10.1002/hbm.21381

De Lean, A., Stadel, J. M., & Lefkowitz, R. J. (1980). A ternary complex model explains the agonist-specific binding properties of the adenylate cyclase-coupled beta-adrenergic receptor. *The Journal of Biological Chemistry*, *255*(15), 7108–7117.

de Vos, C. M. H., Mason, N. L., & Kuypers, K. P. C. (2021). Psychedelics and Neuroplasticity: A Systematic Review Unraveling the Biological Underpinnings of Psychedelics. *Frontiers in Psychiatry*, *12*. https://www.frontiersin.org/article/10.3389/fpsyt.2021.724606

Deco, G., Cabral, J., Saenger, V. M., Boly, M., Tagliazucchi, E., Laufs, H., Van Someren, E., Jobst, B., Stevner, A., & Kringelbach, M. L. (2018). Perturbation of whole-brain dynamics in silico reveals mechanistic differences between brain states. *NeuroImage*, *169*, 46–56. https://doi.org/10.1016/j.neuroimage.2017.12.009

Deco, G., Cruzat, J., Cabral, J., Knudsen, G. M., Carhart-Harris, R. L., Whybrow, P. C., Logothetis, N. K., & Kringelbach, M. L. (2018). Whole-Brain Multimodal Neuroimaging Model Using Serotonin Receptor Maps Explains Non-linear Functional Effects of LSD. *Current Biology*, *28*(19), 3065-3074.e6. https://doi.org/10.1016/j.cub.2018.07.083

Deco, G., Vidaurre, D., & Kringelbach, M. L. (2021). Revisiting the global workspace orchestrating the hierarchical organization of the human brain. *Nature Human Behaviour*, *5*(4), 497–511. https://doi.org/10.1038/s41562-020-01003-6

Dehaene, S., & Changeux, J.-P. (2011). Experimental and theoretical approaches to conscious processing. *Neuron*, *70*(2), 200–227. https://doi.org/10.1016/j.neuron.2011.03.018

Delli Pizzi, S., Chiacchiaretta, P., Sestieri, C., Ferretti, A., Onofrj, M., Della Penna, S., Roseman, L., Timmermann, C., Nutt, D. J., Carhart-Harris, R. L., & Sensi, S. L. (2023). Spatial Correspondence of LSD-Induced Variations on Brain Functioning at Rest With Serotonin Receptor Expression. *Biological Psychiatry. Cognitive Neuroscience and Neuroimaging*, *8*(7), 768–776. https://doi.org/10.1016/j.bpsc.2023.03.009

Desikan, R. S., Ségonne, F., Fischl, B., Quinn, B. T., Dickerson, B. C., Blacker, D., Buckner, R. L., Dale, A. M., Maguire, R. P., Hyman, B. T., Albert, M. S., & Killiany, R. J. (2006). An automated labeling system for subdividing the human cerebral cortex on MRI scans into gyral based regions of interest. *NeuroImage*, *31*(3), 968–980. https://doi.org/10.1016/j.neuroimage.2006.01.021

Duerler, P., Brem, S., Fraga-González, G., Neef, T., Allen, M., Zeidman, P., Stämpfli, P., Vollenweider, F. X., & Preller, K. H. (2021). Psilocybin Induces Aberrant Prediction Error Processing of Tactile Mismatch Responses-A Simultaneous EEG-FMRI Study. *Cerebral Cortex (New York, N.Y.: 1991)*, *32*(1), 186–196. https://doi.org/10.1093/cercor/bhab202

Duerler, P., Schilbach, L., Stämpfli, P., Vollenweider, F. X., & Preller, K. H. (2020). LSD-induced increases in social adaptation to opinions similar to one's own are associated with stimulation of serotonin receptors. *Scientific Reports*, *10*(1), Article 1. https://doi.org/10.1038/s41598-020-68899-y

Dyer, D. C., & Gant, D. W. (1973). Vasoconstriction Produced by Hallucinogens on Isolated Human and Sheep Umbilical Vasculature. *Journal of Pharmacology and Experimental Therapeutics*, *184*(2), 366–375.

Edelsbrunner, Letscher, & Zomorodian. (2002). Topological Persistence and Simplification. *Discrete & Computational Geometry*, *28*(4), 511–533. https://doi.org/10.1007/s00454-002-2885-2

Egger, M., Davey Smith, G., Schneider, M., & Minder, C. (1997). Bias in meta-analysis detected by a simple, graphical test. *BMJ (Clinical Research Ed.)*, *315*(7109), 629–634. https://doi.org/10.1136/bmj.315.7109.629

Eickhoff, S. B., Bzdok, D., Laird, A. R., Kurth, F., & Fox, P. T. (2012). Activation Likelihood Estimation meta-analysis revisited. *Neuroimage*, *59*(3), 2349–2361. https://doi.org/10.1016/j.neuroimage.2011.09.017

Eickhoff, S. B., Laird, A. R., Grefkes, C., Wang, L. E., Zilles, K., & Fox, P. T. (2009). Coordinate-based activation likelihood estimation meta-analysis of neuroimaging data: A random-effects approach based on empirical estimates of spatial uncertainty. *Human Brain Mapping*, *30*(9), 2907–2926. https://doi.org/10.1002/hbm.20718

Erkizia-Santamaría, I., Alles-Pascual, R., Horrillo, I., Meana, J. J., & Ortega, J. E. (2022). Serotonin 5-HT2A, 5-HT2c and 5-HT1A receptor involvement in the acute effects of psilocybin in mice. In vitro pharmacological profile and modulation of thermoregulation and head-twich response. *Biomedicine & Pharmacotherapy = Biomedecine & Pharmacotherapie*, *154*, 113612. https://doi.org/10.1016/j.biopha.2022.113612

Erowid, E., Erowid, F., & Thyssen, S. (2022). *Index (Front Page): Erowid Experience Vaults*. Erowid Experience Vaults. https://erowid.org/experiences/

Eshleman, A. J., Forster, M. J., Wolfrum, K. M., Johnson, R. A., Janowsky, A., & Gatch, M. B. (2014). Behavioral and neurochemical pharmacology of six psychoactive substituted phenethylamines: Mouse locomotion, rat drug discrimination and in vitro receptor and transporter binding and function. *Psychopharmacology*, *231*(5), 875–888. https://doi.org/10.1007/s00213-013-3303-6

Eshleman, A. J., Wolfrum, K. M., Reed, J. F., Kim, S. O., Johnson, R. A., & Janowsky, A. (2018). Neurochemical pharmacology of psychoactive substituted N-benzylphenethylamines: High potency agonists at 5-HT2A receptors. *Biochemical Pharmacology*, *158*, 27–34. https://doi.org/10.1016/j.bcp.2018.09.024

Family, N., Hendricks, P. S., Williams, L. T., Luke, D., Krediet, E., Maillet, E. L., & Raz, S. (2022). Safety, tolerability, pharmacokinetics, and subjective effects of 50, 75, and 100 μg LSD in healthy participants within a novel intervention paradigm: A proof-of-concept study. *Journal of Psychopharmacology*, *36*(3), 321–336. https://doi.org/10.1177/02698811211069103

Friston, K. J., Harrison, L., & Penny, W. (2003). Dynamic causal modelling. *NeuroImage*, *19*(4), 1273–1302. https://doi.org/10.1016/s1053-8119(03)00202-7

Friston, K. J., Kahan, J., Biswal, B., & Razi, A. (2014). A DCM for resting state fMRI. *Neuroimage*, *94*(100), 396–407. https://doi.org/10.1016/j.neuroimage.2013.12.009

Gaddis, A., Lidstone, D. E., Nebel, M. B., Griffiths, R. R., Mostofsky, S. H., Mejia, A. F., & Barrett, F. S. (2022). Psilocybin induces spatially constrained alterations in thalamic functional organizaton and connectivity. *NeuroImage*, *260*, 119434. https://doi.org/10.1016/j.neuroimage.2022.119434

Gatch, M. B., Forster, M. J., Janowsky, A., & Eshleman, A. J. (2011). Abuse liability profile of three substituted tryptamines. *The Journal of Pharmacology and Experimental Therapeutics*, *338*(1), 280–289. https://doi.org/10.1124/jpet.111.179705

Girn, M., Roseman, L., Bernhardt, B., Smallwood, J., Carhart-Harris, R., & Nathan Spreng, R. (2022). Serotonergic psychedelic drugs LSD and psilocybin reduce the hierarchical differentiation of unimodal and transmodal cortex. *NeuroImage*, *256*, 119220. https://doi.org/10.1016/j.neuroimage.2022.119220

Glatfelter, G. C., Pottie, E., Partilla, J. S., Sherwood, A. M., Kaylo, K., Pham, D. N. K., Naeem, M., Sammeta, V. R., DeBoer, S., Golen, J. A., Hulley, E. B., Stove, C. P., Chadeayne, A. R., Manke, D. R., & Baumann, M. H. (2022). Structure–Activity Relationships for Psilocybin, Baeocystin, Aeruginascin, and Related Analogues to Produce Pharmacological Effects in Mice. *ACS Pharmacology & Translational Science*, *5*(11), 1181–1196. https://doi.org/10.1021/acsptsci.2c00177

Gómez-Emilsson, A. (2021). *Healing Trauma With Neural Annealing*. https://www.qri.org/blog/Neural-Annealing

Goodwin, G. M., Croal, M., Marwood, L., & Malievskaia, E. (2023). Unblinding and demand characteristics in the treatment of depression. *Journal of Affective Disorders*, *328*, 1–5. https://doi.org/10.1016/j.jad.2023.02.030

Gosselin, N., Samson, S., Adolphs, R., Noulhiane, M., Roy, M., Hasboun, D., Baulac, M., & Peretz, I. (2006). Emotional responses to unpleasant music correlates with damage to the parahippocampal cortex. *Brain*, *129*(10), 2585–2592. https://doi.org/10.1093/brain/awl240

Gouzoulis-Mayfrank, E., Heekeren, K., Neukirch, A., Stoll, M., Stock, C., Obradovic, M., & Kovar, K.-A. (2005). Psychological effects of (S)-ketamine and N,N-dimethyltryptamine (DMT): A double-blind, cross-over study in healthy volunteers. *Pharmacopsychiatry*, *38*(6), 301–311. https://doi.org/10.1055/s-2005-916185

Griffiths, R. R., Hurwitz, E. S., Davis, A. K., Johnson, M. W., & Jesse, R. (2019). Survey of subjective 'God encounter experiences': Comparisons among naturally occurring experiences and those occasioned by the classic psychedelics psilocybin, LSD, ayahuasca, or DMT. *PLOS ONE*, *14*(4), e0214377. https://doi.org/10.1371/journal.pone.0214377

Grimm, O., Kraehenmann, R., Preller, K. H., Seifritz, E., & Vollenweider, F. X. (2018). Psilocybin modulates functional connectivity of the amygdala during emotional face discrimination. *European Neuropsychopharmacology: The Journal of the European College of Neuropsychopharmacology*, *28*(6), 691–700. https://doi.org/10.1016/j.euroneuro.2018.03.016

Hahn, A., Stein, P., Windischberger, C., Weissenbacher, A., Spindelegger, C., Moser, E., Kasper, S., & Lanzenberger, R. (2011). Reduced resting-state functional connectivity between amygdala and orbitofrontal cortex in social anxiety disorder. *NeuroImage*, *56*(3), 881–889. https://doi.org/10.1016/j.neuroimage.2011.02.064

Halberstadt, A. L., Chatha, M., Klein, A. K., McCorvy, J. D., Meyer, M. R., Wagmann, L., Stratford, A., & Brandt, S. D. (2020). Pharmacological and biotransformation studies of 1-acyl-substituted derivatives of d-lysergic acid diethylamide (LSD). *Neuropharmacology*, *172*, 107856. https://doi.org/10.1016/j.neuropharm.2019.107856

Hansen, J. Y., Shafiei, G., Markello, R. D., Smart, K., Cox, S. M. L., Nørgaard, M., Beliveau, V., Wu, Y., Gallezot, J.-D., Aumont, É., Servaes, S., Scala, S. G., DuBois, J. M., Wainstein, G., Bezgin, G., Funck, T., Schmitz, T. W., Spreng, R. N., Galovic, M., … Misic, B. (2022). Mapping neurotransmitter systems to the structural and functional organization of the human neocortex. *Nature Neuroscience*, *25*(11), Article 11. https://doi.org/10.1038/s41593-022-01186-3

Harrer, M., Cuijpers, P., A, F. T., & Ebert, D. D. (2021). *Doing Meta-Analysis With R: A Hands-On Guide* (1st ed.). Chapman & Hall/CRC Press.

Hase, A., Erdmann, M., Limbach, V., & Hasler, G. (2022). Analysis of recreational psychedelic substance use experiences classified by substance. *Psychopharmacology*, *239*(2), 643–659. https://doi.org/10.1007/s00213-022-06062-3

Hasler, F., Grimberg, U., Benz, M. A., Huber, T., & Vollenweider, F. X. (2004). Acute psychological and physiological effects of psilocybin in healthy humans: A double-blind, placebo-controlled dose-effect study. *Psychopharmacology*, *172*(2), 145–156. https://doi.org/10.1007/s00213-003-1640-6

Hedges, L. V., Tipton, E., & Johnson, M. C. (2010). Robust variance estimation in meta-regression with dependent effect size estimates. *Research Synthesis Methods*, *1*(1), 39–65. https://doi.org/10.1002/jrsm.5

Hedges, L. V., & Vevea, J. L. (1998). Fixed- and random-effects models in meta-analysis. *Psychological Methods*, *3*(4), 486–504. https://doi.org/10.1037/1082-989X.3.4.486

Hesse, J., & Gross, T. (2014). Self-organized criticality as a fundamental property of neural systems. *Frontiers in Systems Neuroscience*, *8*. https://www.frontiersin.org/articles/10.3389/fnsys.2014.00166

Hillman, E. M. C. (2014). Coupling Mechanism and Significance of the BOLD Signal: A Status Report. *Annual Review of Neuroscience*, *37*, 161–181. https://doi.org/10.1146/annurev-neuro-071013-014111

Hjorth, S., Carlsson, A., Lindberg, P., Sanchez, D., Wikström, H., Arvidsson, L.-E., Hacksell, U., & Nilsson, J. L. G. (1982). 8-hydroxy-2-(di-n-propylamino)tetralin, 8-OH-DPAT, a potent and selective simplified ergot congener with central 5-HT-receptor stimulating activity. *Journal of Neural Transmission*, *55*(3), 169–188. https://doi.org/10.1007/BF01276574

Holmes, J., & Nolte, T. (2019). "Surprise" and the Bayesian Brain: Implications for Psychotherapy Theory and Practice. *Frontiers in Psychology*, *10*. https://www.frontiersin.org/articles/10.3389/fpsyg.2019.00592

Holze, F., Ley, L., Müller, F., Becker, A. M., Straumann, I., Vizeli, P., Kuehne, S. S., Roder, M. A., Duthaler, U., Kolaczynska, K. E., Varghese, N., Eckert, A., & Liechti, M. E. (2022). Direct comparison of the acute effects of lysergic acid diethylamide and psilocybin in a double-blind placebo-controlled study in healthy subjects. *Neuropsychopharmacology*, *47*(6), 1180–1187. https://doi.org/10.1038/s41386-022-01297-2

Holze, F., Vizeli, P., Ley, L., Müller, F., Dolder, P., Stocker, M., Duthaler, U., Varghese, N., Eckert, A., Borgwardt, S., & Liechti, M. E. (2021). Acute dose-dependent effects of lysergic acid diethylamide in a double-blind placebo-controlled study in healthy subjects. *Neuropsychopharmacology: Official Publication of the American College of Neuropsychopharmacology*, *46*(3), 537–544. https://doi.org/10.1038/s41386-020-00883-6

Holze, F., Vizeli, P., Müller, F., Ley, L., Duerig, R., Varghese, N., Eckert, A., Borgwardt, S., & Liechti, M. E. (2020). Distinct acute effects of LSD, MDMA, and D-amphetamine in healthy subjects. *Neuropsychopharmacology*, *45*(3), Article 3. https://doi.org/10.1038/s41386-019-0569-3

Hulme, E. C., & Trevethick, M. A. (2010). Ligand binding assays at equilibrium: Validation and interpretation. *British Journal of Pharmacology*, *161*(6), 1219–1237. https://doi.org/10.1111/j.1476-5381.2009.00604.x

Iaria, G., Fox, C., Scheel, M., Stowe, R., & Barton, J. (2009). A case of persistent visual hallucinations of faces following LSD abuse: A functional Magnetic Resonance Imaging study. *Neurocase*, *16*, 106–118. https://doi.org/10.1080/13554790903329141

Janowsky, A., Eshleman, A. J., Johnson, R. A., Wolfrum, K. M., Hinrichs, D. J., Yang, J., Zabriskie, T. M., Smilkstein, M. J., & Riscoe, M. K. (2014). Mefloquine and Psychotomimetics Share Neurotransmitter Receptor and Transporter Interactions In Vitro. *Psychopharmacology*, *231*(14), 2771–2783. https://doi.org/10.1007/s00213-014-3446-0

Jobst, B. M., Atasoy, S., Ponce-Alvarez, A., Sanjuán, A., Roseman, L., Kaelen, M., Carhart-Harris, R., Kringelbach, M. L., & Deco, G. (2021). Increased sensitivity to strong perturbations in a whole-brain model of LSD. *NeuroImage*, *230*, 117809. https://doi.org/10.1016/j.neuroimage.2021.117809

Johnson, M. (2019, November 28). *Neural Annealing: Toward a Neural Theory of Everything.* https://opentheory.net/2019/11/neural-annealing-toward-a-neural-theory-of-everything/

Kaelen, M., Lorenz, R., Barrett, F., Roseman, L., Orban, C., Santos-Ribeiro, A., Wall, M. B., Feilding, A., Nutt, D., Muthukumaraswamy, S., Carhart-Harris, R., & Leech, R. (2017). *Effects of LSD on music-evoked brain activity* (p. 153031). bioRxiv. https://doi.org/10.1101/153031

Kaelen, M., Roseman, L., Kahan, J., Santos-Ribeiro, A., Orban, C., Lorenz, R., Barrett, F. S., Bolstridge, M., Williams, T., Williams, L., Wall, M. B., Feilding, A., Muthukumaraswamy, S., Nutt, D. J., & Carhart-Harris, R. (2016). LSD modulates music-induced imagery via changes in parahippocampal connectivity. *European Neuropsychopharmacology: The Journal of the European College of Neuropsychopharmacology*, *26*(7), 1099–1109. https://doi.org/10.1016/j.euroneuro.2016.03.018

Kawasaki, A., & Purvin, V. (1996). Persistent palinopsia following ingestion of lysergic acid diethylamide (LSD). *Archives of Ophthalmology (Chicago, Ill.: 1960)*, *114*(1), 47–50. https://doi.org/10.1001/archopht.1996.01100130045007

Keiser, M. J., Setola, V., Irwin, J. J., Laggner, C., Abbas, A., Hufeisen, S. J., Jensen, N. H., Kuijer, M. B., Matos, R. C., Tran, T. B., Whaley, R., Glennon, R. A., Hert, J., Thomas, K. L. H., Edwards, D. D., Shoichet, B. K., & Roth, B. L. (2009). Predicting new molecular targets for known drugs. *Nature*, *462*(7270), 175–181. https://doi.org/10.1038/nature08506

Kenakin, T. (2017). A Scale of Agonism and Allosteric Modulation for Assessment of Selectivity, Bias, and Receptor Mutation. *Molecular Pharmacology*, *92*(4), 414–424. https://doi.org/10.1124/mol.117.108787

Kenakin, T., Watson, C., Muniz-Medina, V., Christopoulos, A., & Novick, S. (2011). A Simple Method for Quantifying Functional Selectivity and Agonist Bias. *ACS Chemical Neuroscience*, *3*(3), 193–203. https://doi.org/10.1021/cn200111m

Kenakin, T., Watson, C., Muniz-Medina, V., Christopoulos, A., & Novick, S. (2012). A Simple Method for Quantifying Functional Selectivity and Agonist Bias. *ACS Chemical Neuroscience*, *3*(3), 193–203. https://doi.org/10.1021/cn200111m

Klein, A. K., Chatha, M., Laskowski, L. J., Anderson, E. I., Brandt, S. D., Chapman, S. J., McCorvy, J. D., & Halberstadt, A. L. (2021). Investigation of the Structure-Activity Relationships of Psilocybin Analogues. *ACS Pharmacology & Translational Science*, *4*(2), 533–542. https://doi.org/10.1021/acsptsci.0c00176

Knapp, G., & Hartung, J. (2003). Improved tests for a random effects meta-regression with a single covariate. *Statistics in Medicine*, *22*(17), 2693–2710. https://doi.org/10.1002/sim.1482

Knight, A. R., Misra, A., Quirk, K., Benwell, K., Revell, D., Kennett, G., & Bickerdike, M. (2004). Pharmacological characterisation of the agonist radioligand binding site of 5-HT(2A), 5-HT(2B) and 5-HT(2C) receptors. *Naunyn-Schmiedeberg's Archives of Pharmacology*, *370*(2), 114–123. https://doi.org/10.1007/s00210-004-0951-4

Knill, D. C., & Pouget, A. (2004). The Bayesian brain: The role of uncertainty in neural coding and computation. *Trends in Neurosciences*, *27*(12), 712–719. https://doi.org/10.1016/j.tins.2004.10.007

Knudsen, G. M. (2023). Sustained effects of single doses of classical psychedelics in humans. *Neuropsychopharmacology*, *48*(1), 145–150. https://doi.org/10.1038/s41386-022-01361-x

Koelsch, S. (2014). Brain correlates of music-evoked emotions. *Nature Reviews Neuroscience*, *15*(3), Article 3. https://doi.org/10.1038/nrn3666

Kometer, M., Schmidt, A., Bachmann, R., Studerus, E., Seifritz, E., & Vollenweider, F. X. (2012). Psilocybin biases facial recognition, goal-directed behavior, and mood state toward positive relative to negative emotions through different serotonergic subreceptors. *Biological Psychiatry*, *72*(11), 898–906. https://doi.org/10.1016/j.biopsych.2012.04.005

Kozell, L. B., Eshleman, A. J., Swanson, T. L., Bloom, S. H., Wolfrum, K. M., Schmachtenberg, J. L., Olson, R. J., Janowsky, A., & Abbas, A. I. (2023). Pharmacologic Activity of Substituted Tryptamines at 5-Hydroxytryptamine (5-HT)2A Receptor (5-HT2AR), 5-HT2CR, 5-HT1AR, and Serotonin Transporter. *The Journal of Pharmacology and Experimental Therapeutics*, *385*(1), 62–75. https://doi.org/10.1124/jpet.122.001454

Kraehenmann, R. (2017). Dreams and Psychedelics: Neurophenomenological Comparison and Therapeutic Implications. *Current Neuropharmacology*, *15*(7), 1032–1042. https://doi.org/10.2174/1573413713666170619092629

Kraehenmann, R., Pokorny, D., Aicher, H., Preller, K. H., Pokorny, T., Bosch, O. G., Seifritz, E., & Vollenweider, F. X. (2017). LSD Increases Primary Process Thinking via Serotonin 2A Receptor Activation. *Frontiers in Pharmacology*, *8*, 814. https://doi.org/10.3389/fphar.2017.00814

Kraehenmann, R., Preller, K. H., Scheidegger, M., Pokorny, T., Bosch, O. G., Seifritz, E., & Vollenweider, F. X. (2015). Psilocybin-Induced Decrease in Amygdala Reactivity Correlates with Enhanced Positive Mood in Healthy Volunteers. *Biological Psychiatry*, *78*(8), 572–581. https://doi.org/10.1016/j.biopsych.2014.04.010

Kraehenmann, R., Schmidt, A., Friston, K., Preller, K. H., Seifritz, E., & Vollenweider, F. X. (2015). The mixed serotonin receptor agonist psilocybin reduces threat-induced modulation of amygdala connectivity. *NeuroImage: Clinical*, *11*, 53–60. https://doi.org/10.1016/j.nicl.2015.08.009

Kringelbach, M. L., Cruzat, J., Cabral, J., Knudsen, G. M., Carhart-Harris, R., Whybrow, P. C., Logothetis, N. K., & Deco, G. (2020). Dynamic coupling of whole-brain neuronal and neurotransmitter systems. *Proceedings of the National Academy of Sciences*, *117*(17), 9566–9576. https://doi.org/10.1073/pnas.1921475117

Kumral, D., Şansal, F., Cesnaite, E., Mahjoory, K., Al, E., Gaebler, M., Nikulin, V. V., & Villringer, A. (2020). BOLD and EEG signal variability at rest differently relate to aging in the human brain. *NeuroImage*, *207*, 116373. https://doi.org/10.1016/j.neuroimage.2019.116373

Lebedev, A. V., Kaelen, M., Lövdén, M., Nilsson, J., Feilding, A., Nutt, D. J., & Carhart-Harris, R. L. (2016). LSD-induced entropic brain activity predicts subsequent personality change. *Human Brain Mapping*, *37*(9), 3203–3213. https://doi.org/10.1002/hbm.23234

Lebedev, A. V., Lövdén, M., Rosenthal, G., Feilding, A., Nutt, D. J., & Carhart-Harris, R. L. (2015). Finding the self by losing the self: Neural correlates of ego-dissolution under psilocybin. *Human Brain Mapping*, *36*(8), 3137–3153. https://doi.org/10.1002/hbm.22833

Lempel, A., & Ziv, J. (1976). On the Complexity of Finite Sequences. *IEEE Transactions on Information Theory*, *22*(1), 75–81. IEEE Transactions on Information Theory. https://doi.org/10.1109/TIT.1976.1055501

Lewis, C. R., Preller, K. H., Kraehenmann, R., Michels, L., Staempfli, P., & Vollenweider, F. X. (2017). Two dose investigation of the 5-HT-agonist psilocybin on relative and global cerebral blood flow. *NeuroImage*, *159*, 70–78. https://doi.org/10.1016/j.neuroimage.2017.07.020

Ley, L., Holze, F., Arikci, D., Becker, A. M., Straumann, I., Klaiber, A., Coviello, F., Dierbach, S., Thomann, J., Duthaler, U., Luethi, D., Varghese, N., Eckert, A., & Liechti, M. E. (2023). Comparative acute effects of mescaline, lysergic acid diethylamide, and psilocybin in a randomized, double-blind, placebo-controlled cross-over study in healthy participants. *Neuropsychopharmacology*, 1–9. https://doi.org/10.1038/s41386-023-01607-2

López-Giménez, J. F., & González-Maeso, J. (2018). Hallucinogens and Serotonin 5-HT2A Receptor-Mediated Signaling Pathways. *Current Topics in Behavioral Neurosciences*, *36*, 45–73. https://doi.org/10.1007/7854_2017_478

Lord, L.-D., Expert, P., Atasoy, S., Roseman, L., Rapuano, K., Lambiotte, R., Nutt, D. J., Deco, G., Carhart-Harris, R. L., Kringelbach, M. L., & Cabral, J. (2019). Dynamical exploration of the repertoire of brain networks at rest is modulated by psilocybin. *NeuroImage*, *199*, 127–142. https://doi.org/10.1016/j.neuroimage.2019.05.060

Luan, L., Eckernäs, E., Ashton, M., Rosas, F., Uthaug, M., Bartha, A., Jagger, S., Gascon-Perai, K., Gomes, L., Nutt, D., Erritzoe, D., Carhart-Harris, R., & Timmermann, C. (2023). *Psychological and physiological effects of extended DMT*. PsyArXiv. https://doi.org/10.31234/osf.io/vg4dp

Luethi, D., Hoener, M. C., Krähenbühl, S., Liechti, M. E., & Duthaler, U. (2019). Cytochrome P450 enzymes contribute to the metabolism of LSD to nor-LSD and 2-oxo-3-hydroxy-LSD: Implications for clinical LSD use. *Biochemical Pharmacology*, *164*, 129–138. https://doi.org/10.1016/j.bcp.2019.04.013

Luethi, D., Trachsel, D., Hoener, M. C., & Liechti, M. E. (2018). Monoamine receptor interaction profiles of 4-thio-substituted phenethylamines (2C-T drugs). *Neuropharmacology*, *134*(Pt A), 141–148. https://doi.org/10.1016/j.neuropharm.2017.07.012

Luppi, A. I., Carhart-Harris, R. L., Roseman, L., Pappas, I., Menon, D. K., & Stamatakis, E. A. (2021). LSD alters dynamic integration and segregation in the human brain. *NeuroImage*, *227*, 117653. https://doi.org/10.1016/j.neuroimage.2020.117653

Luppi, A. I., Vohryzek, J., Kringelbach, M. L., Mediano, P. A. M., Craig, M. M., Adapa, R., Carhart-Harris, R. L., Roseman, L., Pappas, I., Peattie, A. R. D., Manktelow, A. E., Sahakian, B. J., Finoia, P., Williams, G. B., Allanson, J., Pickard, J. D., Menon, D. K., Atasoy, S., & Stamatakis, E. A. (2023). Distributed harmonic patterns of structure-function dependence orchestrate human consciousness. *Communications Biology*, *6*(1), Article 1. https://doi.org/10.1038/s42003-023-04474-1

Madsen, M. K., Stenbæk, D. S., Arvidsson, A., Armand, S., Marstrand-Joergensen, M. R., Johansen, S. S., Linnet, K., Ozenne, B., Knudsen, G. M., & Fisher, P. M. (2021). Psilocybin-induced changes in brain network integrity and segregation correlate with plasma psilocin level and psychedelic experience. *European Neuropsychopharmacology*, *50*, 121–132. https://doi.org/10.1016/j.euroneuro.2021.06.001

Makowski, C., Lepage, M., & Evans, A. C. (2019). Head motion: The dirty little secret of neuroimaging in psychiatry. *Journal of Psychiatry & Neuroscience: JPN*, *44*(1), 62–68. https://doi.org/10.1503/jpn.180022

Malén, T., Karjalainen, T., Isojärvi, J., Vehtari, A., Bürkner, P.-C., Putkinen, V., Kaasinen, V., Hietala, J., Nuutila, P., Rinne, J., & Nummenmaa, L. (2022). Atlas of type 2 dopamine receptors in the human brain: Age and sex dependent variability in a large PET cohort. *NeuroImage*, *255*, 119149. https://doi.org/10.1016/j.neuroimage.2022.119149

Mallaroni, P., Mason, N. L., Reckweg, J. T., Paci, R., Ritscher, S., Toennes, S. W., Theunissen, E. L., Kuypers, K. P. C., & Ramaekers, J. G. (2023). Assessment of the Acute Effects of 2C-B vs. Psilocybin on Subjective Experience, Mood, and Cognition. *Clinical Pharmacology and Therapeutics*, *114*(2), 423–433. https://doi.org/10.1002/cpt.2958

Mason, N. L., Kuypers, K. P. C., Müller, F., Reckweg, J., Tse, D. H. Y., Toennes, S. W., Hutten, N. R. P. W., Jansen, J. F. A., Stiers, P., Feilding, A., & Ramaekers, J. G. (2020). Me, myself, bye: Regional alterations in glutamate and the experience of ego dissolution with psilocybin. *Neuropsychopharmacology*, *45*(12), Article 12. https://doi.org/10.1038/s41386-020-0718-8

Mason, N. L., Kuypers, K. P. C., Reckweg, J. T., Müller, F., Tse, D. H. Y., Da Rios, B., Toennes, S. W., Stiers, P., Feilding, A., & Ramaekers, J. G. (2021). Spontaneous and deliberate creative cognition during and after psilocybin exposure. *Translational Psychiatry*, *11*(1), Article 1. https://doi.org/10.1038/s41398-021-01335-5

Masuda, N., Sakaki, M., Ezaki, T., & Watanabe, T. (2018). Clustering Coefficients for Correlation Networks. *Frontiers in Neuroinformatics*, *12*. https://www.frontiersin.org/articles/10.3389/fninf.2018.00007

McAlpine, R., Blackburne, G., & Kamboj, S. (2023). *Development and psychometric validation of a novel scale for measuring 'psychedelic preparedness'*. PsyArXiv. https://doi.org/10.31234/osf.io/gw9jp

McCulloch, D. E.-W., Olsen, A. S., Ozenne, B., Stenbæk, D. S., Armand, S., Madsen, M. K., Knudsen, G. M., & Fisher, P. M. (2023). *Navigating the chaos of psychedelic neuroimaging: A multi-metric evaluation of acute psilocybin effects on brain entropy* (p. 2023.07.03.23292164). medRxiv. https://doi.org/10.1101/2023.07.03.23292164

Mediano, P. A. M., Rosas, F. E., Timmermann, C., Roseman, L., Nutt, D. J., Feilding, A., Kaelen, M., Kringelbach, M. L., Barrett, A. B., Seth, A. K., Muthukumaraswamy, S., Bor, D., & Carhart-Harris, R. L. (2020). *Effects of external stimulation on psychedelic state neurodynamics* (p. 2020.11.01.356071). bioRxiv. https://doi.org/10.1101/2020.11.01.356071

Mendez, D., Gaulton, A., Bento, A. P., Chambers, J., De Veij, M., Félix, E., Magariños, M. P., Mosquera, J. F., Mutowo, P., Nowotka, M., Gordillo-Marañón, M., Hunter, F., Junco, L., Mugumbate, G., Rodriguez-Lopez, M., Atkinson, F., Bosc, N., Radoux, C. J., Segura-Cabrera, A., … Leach, A. R. (2019). ChEMBL: Towards direct deposition of bioassay data. *Nucleic Acids Research*, *47*(D1), D930–D940. https://doi.org/10.1093/nar/gky1075

Moujaes, F., Preller, K. H., Ji, J. L., Murray, J. D., Berkovitch, L., Vollenweider, F. X., & Anticevic, A. (2023). Toward Mapping Neurobehavioral Heterogeneity of Psychedelic Neurobiology in Humans. *Biological Psychiatry*, *93*(12), 1061–1070. https://doi.org/10.1016/j.biopsych.2022.10.021

Müller, F., Dolder, P. C., Schmidt, A., Liechti, M. E., & Borgwardt, S. (2018). Altered network hub connectivity after acute LSD administration. *NeuroImage. Clinical*, *18*, 694–701. https://doi.org/10.1016/j.nicl.2018.03.005

Müller, F., Lenz, C., Dolder, P. C., Harder, S., Schmid, Y., Lang, U. E., Liechti, M. E., & Borgwardt, S. (2017). Acute effects of LSD on amygdala activity during processing of fearful stimuli in healthy subjects. *Translational Psychiatry*, *7*(4), e1084. https://doi.org/10.1038/tp.2017.54

Müller, F., Lenz, C., Dolder, P., Lang, U., Schmidt, A., Liechti, M., & Borgwardt, S. (2017). Increased thalamic resting-state connectivity as a core driver of LSD-induced hallucinations. *Acta Psychiatrica Scandinavica*, *136*(6), 648–657. https://doi.org/10.1111/acps.12818

Newton, R. A., Phipps, S. L., Flanigan, T. P., Newberry, N. R., Carey, J. E., Kumar, C., McDonald, B., Chen, C., & Elliott, J. M. (1996). Characterisation of Human 5-Hydroxytryptamine2A and 5-Hydroxytryptamine2C Receptors Expressed in the Human Neuroblastoma Cell Line SH-SY5Y: Comparative Stimulation by Hallucinogenic Drugs. *Journal of Neurochemistry*, *67*(6), 2521–2531. https://doi.org/10.1046/j.1471-4159.1996.67062521.x

Nichols, D. E. (2016). Psychedelics. *Pharmacological Reviews*, *68*(2), 264–355. https://doi.org/10.1124/pr.115.011478

Oizumi, M., Albantakis, L., & Tononi, G. (2014). From the Phenomenology to the Mechanisms of Consciousness: Integrated Information Theory 3.0. *PLOS Computational Biology*, *10*(5), e1003588. https://doi.org/10.1371/journal.pcbi.1003588

Olsen, A. S., Lykkebo-Valløe, A., Ozenne, B., Madsen, M. K., Stenbæk, D. S., Armand, S., Mørup, M., Ganz, M., Knudsen, G. M., & Fisher, P. M. (2022). Psilocybin modulation of time-varying functional connectivity is associated with plasma psilocin and subjective effects. *NeuroImage*, *264*, 119716. https://doi.org/10.1016/j.neuroimage.2022.119716

Ort, A., Smallridge, J. W., Sarasso, S., Casarotto, S., von Rotz, R., Casanova, A., Seifritz, E., Preller, K. H., Tononi, G., & Vollenweider, F. X. (2023). TMS-EEG and resting-state EEG applied to altered states of consciousness: Oscillations, complexity, and phenomenology. *iScience*, *26*(5), 106589. https://doi.org/10.1016/j.isci.2023.106589

Özbay, P. S., Chang, C., Picchioni, D., Mandelkow, H., Chappel-Farley, M. G., van Gelderen, P., de Zwart, J. A., & Duyn, J. (2019). Sympathetic activity contributes to the fMRI signal. *Communications Biology*, *2*(1), Article 1. https://doi.org/10.1038/s42003-019-0659-0

Özbay, P. S., Chang, C., Picchioni, D., Mandelkow, H., Moehlman, T. M., Chappel-Farley, M. G., van Gelderen, P., de Zwart, J. A., & Duyn, J. H. (2018). Contribution of systemic vascular effects to fMRI activity in white matter. *NeuroImage*, *176*, 541–549. https://doi.org/10.1016/j.neuroimage.2018.04.045

Page, M. J., McKenzie, J. E., Bossuyt, P. M., Boutron, I., Hoffmann, T. C., Mulrow, C. D., Shamseer, L., Tetzlaff, J. M., Akl, E. A., Brennan, S. E., Chou, R., Glanville, J., Grimshaw, J. M., Hróbjartsson, A., Lalu, M. M., Li, T., Loder, E. W., Mayo-Wilson, E., McDonald, S., … Moher, D. (2021). The PRISMA 2020 statement: An updated guideline for reporting systematic reviews. *BMJ*, *372*, n71. https://doi.org/10.1136/bmj.n71

Palhano-Fontes, F., Andrade, K. C., Tofoli, L. F., Santos, A. C., Crippa, J. A. S., Hallak, J. E. C., Ribeiro, S., & Araujo, D. B. de. (2015). The Psychedelic State Induced by Ayahuasca Modulates the Activity and Connectivity of the Default Mode Network. *PLOS ONE*, *10*(2), e0118143. https://doi.org/10.1371/journal.pone.0118143

Pallavicini, C., Cavanna, F., Zamberlan, F., de la Fuente, L. A., Ilksoy, Y., Perl, Y. S., Arias, M., Romero, C., Carhart-Harris, R., Timmermann, C., & Tagliazucchi, E. (2021). Neural and subjective effects of inhaled N,N-dimethyltryptamine in natural settings. *Journal of Psychopharmacology (Oxford, England)*, *35*(4), 406–420. https://doi.org/10.1177/0269881120981384

Passie, T., Seifert, J., Schneider, U., & Emrich, H. M. (2002). The pharmacology of psilocybin. *Addiction Biology*, *7*(4), 357–364. https://doi.org/10.1080/1355621021000005937

Petri, G., Expert, P., Turkheimer, F., Carhart-Harris, R., Nutt, D., Hellyer, P. J., & Vaccarino, F. (2014). Homological scaffolds of brain functional networks. *Journal of The Royal Society Interface*, *11*(101), 20140873. https://doi.org/10.1098/rsif.2014.0873

Pierce, P. A., & Peroutka, S. J. (1989). Hallucinogenic drug interactions with neurotransmitter receptor binding sites in human cortex. *Psychopharmacology*, *97*(1), 118–122. https://doi.org/10.1007/BF00443425

Pokorny, T., Duerler, P., Seifritz, E., Vollenweider, F. X., & Preller, K. H. (2020). LSD acutely impairs working memory, executive functions, and cognitive flexibility, but not risk-based decision-making. *Psychological Medicine*, *50*(13), 2255–2264. https://doi.org/10.1017/S0033291719002393

Pokorny, T., Preller, K. H., Kometer, M., Dziobek, I., & Vollenweider, F. X. (2017). Effect of Psilocybin on Empathy and Moral Decision-Making. *The International Journal of Neuropsychopharmacology*, *20*(9), 747–757. https://doi.org/10.1093/ijnp/pyx047

Pokorny, T., Preller, K. H., Kraehenmann, R., & Vollenweider, F. X. (2016). Modulatory effect of the 5-HT1A agonist buspirone and the mixed non-hallucinogenic 5-HT1A/2A agonist ergotamine on psilocybin-induced psychedelic experience. *European Neuropsychopharmacology*, *26*(4), 756–766. https://doi.org/10.1016/j.euroneuro.2016.01.005

Porter, R. H., Benwell, K. R., Lamb, H., Malcolm, C. S., Allen, N. H., Revell, D. F., Adams, D. R., & Sheardown, M. J. (1999). Functional characterization of agonists at recombinant human 5-

HT2A, 5-HT2B and 5-HT2C receptors in CHO-K1 cells. *British Journal of Pharmacology*, *128*(1), 13–20. https://doi.org/10.1038/sj.bjp.0702751

Pottie, E., Cannaert, A., & Stove, C. P. (2020). In vitro structure-activity relationship determination of 30 psychedelic new psychoactive substances by means of β-arrestin 2 recruitment to the serotonin 2A receptor. *Archives of Toxicology*, *94*(10), 3449–3460. https://doi.org/10.1007/s00204-020-02836-w

Pottie, E., Dedecker, P., & Stove, C. P. (2020). Identification of psychedelic new psychoactive substances (NPS) showing biased agonism at the 5-HT2AR through simultaneous use of β-arrestin 2 and miniGαq bioassays. *Biochemical Pharmacology*, *182*, 114251. https://doi.org/10.1016/j.bcp.2020.114251

Preller, K. H., Burt, J. B., Ji, J. L., Schleifer, C. H., Adkinson, B. D., Stämpfli, P., Seifritz, E., Repovs, G., Krystal, J. H., Murray, J. D., Vollenweider, F. X., & Anticevic, A. (2018). Changes in global and thalamic brain connectivity in LSD-induced altered states of consciousness are attributable to the 5-HT2A receptor. *eLife*, *7*, e35082. https://doi.org/10.7554/eLife.35082

Preller, K. H., Duerler, P., Burt, J. B., Ji, J. L., Adkinson, B., Stämpfli, P., Seifritz, E., Repovš, G., Krystal, J. H., Murray, J. D., Anticevic, A., & Vollenweider, F. X. (2020). Psilocybin Induces Time-Dependent Changes in Global Functional Connectivity. *Biological Psychiatry*, *88*(2), 197–207. https://doi.org/10.1016/j.biopsych.2019.12.027

Preller, K. H., Herdener, M., Pokorny, T., Planzer, A., Kraehenmann, R., Stämpfli, P., Liechti, M. E., Seifritz, E., & Vollenweider, F. X. (2017). The Fabric of Meaning and Subjective Effects in LSD-Induced States Depend on Serotonin 2A Receptor Activation. *Current Biology: CB*, *27*(3), 451–457. https://doi.org/10.1016/j.cub.2016.12.030

Preller, K. H., Pokorny, T., Hock, A., Kraehenmann, R., Stämpfli, P., Seifritz, E., Scheidegger, M., & Vollenweider, F. X. (2016). Effects of serotonin 2A/1A receptor stimulation on social exclusion processing. *Proceedings of the National Academy of Sciences of the United States of America*, *113*(18), 5119–5124. https://doi.org/10.1073/pnas.1524187113

Preller, K. H., Razi, A., Zeidman, P., Stämpfli, P., Friston, K. J., & Vollenweider, F. X. (2019). Effective connectivity changes in LSD-induced altered states of consciousness in humans. *Proceedings of the National Academy of Sciences*, *116*(7), 2743–2748. https://doi.org/10.1073/pnas.1815129116

Preller, K. H., Schilbach, L., Pokorny, T., Flemming, J., Seifritz, E., & Vollenweider, F. X. (2018). Role of the 5-HT2A Receptor in Self- and Other-Initiated Social Interaction in Lysergic Acid Diethylamide-Induced States: A Pharmacological fMRI Study. *Journal of Neuroscience*, *38*(14), 3603–3611. https://doi.org/10.1523/JNEUROSCI.1939-17.2018

Prugger, J., Derdiyok, E., Dinkelacker, J., Costines, C., & Schmidt, T. T. (2022). The Altered States Database: Psychometric data from a systematic literature review. *Scientific Data*, *9*(1), Article 1. https://doi.org/10.1038/s41597-022-01822-4

Pustejovsky, J. E., & Tipton, E. (2022). Meta-analysis with Robust Variance Estimation: Expanding the Range of Working Models. *Prevention Science*, *23*(3), 425–438. https://doi.org/10.1007/s11121-021-01246-3

Qiu, T. (Tim), & Minda, J. P. (2021). *Recreational Psychedelic Users Frequently Encounter Complete Mystical Experiences: Trip Content and Implications for Wellbeing*. PsyArXiv. https://doi.org/10.31234/osf.io/xrbzs

Quednow, B. B., Kometer, M., Geyer, M. A., & Vollenweider, F. X. (2012). Psilocybin-Induced Deficits in Automatic and Controlled Inhibition are Attenuated by Ketanserin in Healthy Human Volunteers. *Neuropsychopharmacology*, *37*(3), Article 3. https://doi.org/10.1038/npp.2011.228

Ray, T. S. (2010). Psychedelics and the Human Receptorome. *PLoS ONE*, *5*(2), e9019. https://doi.org/10.1371/journal.pone.0009019

Rickli, A., Luethi, D., Reinisch, J., Buchy, D., Hoener, M. C., & Liechti, M. E. (2015). Receptor interaction profiles of novel N-2-methoxybenzyl (NBOMe) derivatives of 2,5-dimethoxy-substituted phenethylamines (2C drugs). *Neuropharmacology*, *99*, 546–553. https://doi.org/10.1016/j.neuropharm.2015.08.034

Rickli, A., Moning, O. D., Hoener, M. C., & Liechti, M. E. (2016). Receptor interaction profiles of novel psychoactive tryptamines compared with classic hallucinogens. *European Neuropsychopharmacology: The Journal of the European College of Neuropsychopharmacology*, *26*(8), 1327–1337. https://doi.org/10.1016/j.euroneuro.2016.05.001

Roseman, L., Leech, R., Feilding, A., Nutt, D. J., & Carhart-Harris, R. L. (2014). The effects of psilocybin and MDMA on between-network resting state functional connectivity in healthy volunteers. *Frontiers in Human Neuroscience*, *8*. https://www.frontiersin.org/article/10.3389/fnhum.2014.00204

Roseman, L., Sereno, M. I., Leech, R., Kaelen, M., Orban, C., McGonigle, J., Feilding, A., Nutt, D. J., & Carhart-Harris, R. L. (2016). LSD alters eyes-closed functional connectivity within the early visual cortex in a retinotopic fashion. *Human Brain Mapping*, *37*(8), 3031–3040. https://doi.org/10.1002/hbm.23224

Roth, B. L., Lopez, E., Patel, S., & Kroeze, W. K. (2000). The Multiplicity of Serotonin Receptors: Uselessly Diverse Molecules or an Embarrassment of Riches? *The Neuroscientist*, *6*(4), 252–262. https://doi.org/10.1177/107385840000600408

Rubinov, M., & Sporns, O. (2010). Complex network measures of brain connectivity: Uses and interpretations. *NeuroImage*, *52*(3), 1059–1069. https://doi.org/10.1016/j.neuroimage.2009.10.003

Ruffini, G., Damiani, G., Lozano-Soldevilla, D., Deco, N., Rosas, F. E., Kiani, N. A., Ponce-Alvarez, A., Kringelbach, M. L., Carhart-Harris, R., & Deco, G. (2023). LSD-induced increase of Ising temperature and algorithmic complexity of brain dynamics. *PLOS Computational Biology*, *19*(2), e1010811. https://doi.org/10.1371/journal.pcbi.1010811

Sandiego, C. M., Gallezot, J.-D., Lim, K., Ropchan, J., Lin, S., Gao, H., Morris, E. D., & Cosgrove, K. P. (2015). Reference region modeling approaches for amphetamine challenge studies with [11C]FLB 457 and PET. *Journal of Cerebral Blood Flow and Metabolism: Official Journal of the International Society of Cerebral Blood Flow and Metabolism*, *35*(4), 623–629. https://doi.org/10.1038/jcbfm.2014.237

Sanz, C., Zamberlan, F., Erowid, E., Erowid, F., & Tagliazucchi, E. (2018). The Experience Elicited by Hallucinogens Presents the Highest Similarity to Dreaming within a Large Database of Psychoactive Substance Reports. *Frontiers in Neuroscience*, *12*. https://doi.org/10.3389/fnins.2018.00007

Schaefer, A., Kong, R., Gordon, E. M., Laumann, T. O., Zuo, X.-N., Holmes, A. J., Eickhoff, S. B., & Yeo, B. T. T. (2018). Local-Global Parcellation of the Human Cerebral Cortex from Intrinsic Functional Connectivity MRI. *Cerebral Cortex (New York, NY)*, *28*(9), 3095–3114. https://doi.org/10.1093/cercor/bhx179

Schartner, M. M., Carhart-Harris, R. L., Barrett, A. B., Seth, A. K., & Muthukumaraswamy, S. D. (2017). Increased spontaneous MEG signal diversity for psychoactive doses of ketamine, LSD and psilocybin. *Scientific Reports*, *7*(1), Article 1. https://doi.org/10.1038/srep46421

Schmid, Y., Enzler, F., Gasser, P., Grouzmann, E., Preller, K. H., Vollenweider, F. X., Brenneisen, R., Müller, F., Borgwardt, S., & Liechti, M. E. (2015). Acute Effects of Lysergic Acid Diethylamide in Healthy Subjects. *Biological Psychiatry*, *78*(8), 544–553. https://doi.org/10.1016/j.biopsych.2014.11.015

Schmidt, A., Bachmann, R., Kometer, M., Csomor, P. A., Stephan, K. E., Seifritz, E., & Vollenweider, F. X. (2012). Mismatch negativity encoding of prediction errors predicts S-ketamine-induced cognitive impairments. *Neuropsychopharmacology: Official Publication of the American College of Neuropsychopharmacology*, *37*(4), 865–875. https://doi.org/10.1038/npp.2011.261

Schmidt, A., Müller, F., Lenz, C., Dolder, P. C., Schmid, Y., Zanchi, D., Lang, U. E., Liechti, M. E., & Borgwardt, S. (2018). Acute LSD effects on response inhibition neural networks. *Psychological Medicine*, *48*(9), 1464–1473. https://doi.org/10.1017/S0033291717002914

Schmidt, T. T., & Berkemeyer, H. (2018). The Altered States Database: Psychometric Data of Altered States of Consciousness. *Frontiers in Psychology*, *9*. https://www.frontiersin.org/articles/10.3389/fpsyg.2018.01028

Schmitz, G. P., Jain, M. K., Slocum, S. T., & Roth, B. L. (2022). 5-HT2A SNPs Alter the Pharmacological Signaling of Potentially Therapeutic Psychedelics. *ACS Chemical Neuroscience*, *13*(16), 2386–2398. https://doi.org/10.1021/acschemneuro.1c00815

Schünemann, H. J., Higgins, J. P., Vist, G. E., Glasziou, P., Akl, E. A., Skoetz, N., Guyatt, G. H., & on behalf of the Cochrane GRADEing Methods Group (formerly Applicability and Recommendations Methods Group) and the Cochrane Statistical Methods Group. (2019). Completing 'Summary of findings' tables and grading the certainty of the evidence. In *Cochrane Handbook for Systematic Reviews of Interventions* (pp. 375–402). John Wiley & Sons, Ltd. https://doi.org/10.1002/9781119536604.ch14

Seeley, W. W., Menon, V., Schatzberg, A. F., Keller, J., Glover, G. H., Kenna, H., Reiss, A. L., & Greicius, M. D. (2007). Dissociable Intrinsic Connectivity Networks for Salience Processing and Executive Control. *Journal of Neuroscience*, *27*(9), 2349–2356. https://doi.org/10.1523/JNEUROSCI.5587-06.2007

Seth, A. K., & Friston, K. J. (2016). Active interoceptive inference and the emotional brain. *Philosophical Transactions of the Royal Society B: Biological Sciences*, *371*(1708), 20160007. https://doi.org/10.1098/rstb.2016.0007

Shannon, C. E. (1948). A Mathematical Theory of Communication. *Bell System Technical Journal*, *27*(3), 379–423. https://doi.org/10.1002/j.1538-7305.1948.tb01338.x

Shen, X., Tokoglu, F., Papademetris, X., & Constable, R. T. (2013). Groupwise whole-brain parcellation from resting-state fMRI data for network node identification. *NeuroImage*, *82*, 403–415. https://doi.org/10.1016/j.neuroimage.2013.05.081

Sherwood, A. M., Halberstadt, A. L., Klein, A. K., McCorvy, J. D., Kaylo, K. W., Kargbo, R. B., & Meisenheimer, P. (2020). Synthesis and Biological Evaluation of Tryptamines Found in Hallucinogenic Mushrooms: Norbaeocystin, Baeocystin, Norpsilocin, and Aeruginascin. *Journal of Natural Products*, *83*(2), 461–467. https://doi.org/10.1021/acs.jnatprod.9b01061

Singleton, S. P., Luppi, A. I., Carhart-Harris, R. L., Cruzat, J., Roseman, L., Nutt, D. J., Deco, G., Kringelbach, M. L., Stamatakis, E. A., & Kuceyeski, A. (2022). *LSD and psilocybin flatten the brain's energy landscape: Insights from receptor-informed network control theory* (p. 2021.05.14.444193). bioRxiv. https://doi.org/10.1101/2021.05.14.444193

Smigielski, L., Kometer, M., Scheidegger, M., Stress, C., Preller, K. H., Koenig, T., & Vollenweider, F. X. (2020). P300-mediated modulations in self–other processing under psychedelic psilocybin are related to connectedness and changed meaning: A window into the self–other overlap. *Human Brain Mapping*, *41*(17), 4982–4996. https://doi.org/10.1002/hbm.25174

Smith, C. T., Crawford, J. L., Dang, L. C., Seaman, K. L., San Juan, M. D., Vijay, A., Katz, D. T., Matuskey, D., Cowan, R. L., Morris, E. D., Zald, D. H., & Samanez-Larkin, G. R. (2019). Partial-volume correction increases estimated dopamine D2-like receptor binding potential and reduces adult age differences. *Journal of Cerebral Blood Flow and Metabolism: Official Journal of* *the International Society of Cerebral Blood Flow and Metabolism*, *39*(5), 822–833. https://doi.org/10.1177/0271678X17737693

Smith, S. M., Fox, P. T., Miller, K. L., Glahn, D. C., Fox, P. M., Mackay, C. E., Filippini, N., Watkins, K. E., Toro, R., Laird, A. R., & Beckmann, C. F. (2009). Correspondence of the brain's functional architecture during activation and rest. *Proceedings of the National Academy of Sciences*, *106*(31), 13040–13045. https://doi.org/10.1073/pnas.0905267106

Speth, J., Speth, C., Kaelen, M., Schloerscheidt, A. M., Feilding, A., Nutt, D. J., & Carhart-Harris, R. L. (2016). Decreased mental time travel to the past correlates with default-mode network disintegration under lysergic acid diethylamide. *Journal of Psychopharmacology (Oxford, England)*, *30*(4), 344–353. https://doi.org/10.1177/0269881116628430

Stephan, K. E., & Friston, K. J. (2010). Analyzing effective connectivity with fMRI. *Wiley Interdisciplinary Reviews. Cognitive Science*, *1*(3), 446–459. https://doi.org/10.1002/wcs.58

Sterne, J. A., Hernán, M. A., McAleenan, A., Reeves, B. C., & Higgins, J. P. (2019). Assessing risk of bias in a non-randomized study. In *Cochrane Handbook for Systematic Reviews of Interventions* (pp. 621–641). John Wiley & Sons, Ltd. https://doi.org/10.1002/9781119536604.ch25

Strange, P. G. (1999). G-protein coupled receptors: Conformations and states. *Biochemical Pharmacology*, *58*(7), 1081–1088. https://doi.org/10.1016/s0006-2952(99)00144-6

Strange, P. G. (2008). Agonist binding, agonist affinity and agonist efficacy at G protein-coupled receptors. *British Journal of Pharmacology*, *153*(7), 1353–1363. https://doi.org/10.1038/sj.bjp.0707672

Straumann, I., Ley, L., Holze, F., Becker, A. M., Klaiber, A., Wey, K., Duthaler, U., Varghese, N., Eckert, A., & Liechti, M. E. (2023). Acute effects of MDMA and LSD co-administration in a double-blind placebo-controlled study in healthy participants. *Neuropsychopharmacology: Official Publication of the American College of Neuropsychopharmacology*. https://doi.org/10.1038/s41386-023-01609-0

Studerus, E., Kometer, M., Hasler, F., & Vollenweider, F. X. (2011). Acute, subacute and long-term subjective effects of psilocybin in healthy humans: A pooled analysis of experimental studies. *Journal of Psychopharmacology (Oxford, England)*, *25*(11), 1434–1452. https://doi.org/10.1177/0269881110382466

Tagliazucchi, E., Carhart-Harris, R., Leech, R., Nutt, D., & Chialvo, D. R. (2014). Enhanced repertoire of brain dynamical states during the psychedelic experience. *Human Brain Mapping*, *35*(11), 5442–5456. https://doi.org/10.1002/hbm.22562

Tagliazucchi, E., Roseman, L., Kaelen, M., Orban, C., Muthukumaraswamy, S. D., Murphy, K., Laufs, H., Leech, R., McGonigle, J., Crossley, N., Bullmore, E., Williams, T., Bolstridge, M., Feilding, A., Nutt, D. J., & Carhart-Harris, R. (2016). Increased Global Functional Connectivity Correlates with LSD-Induced Ego Dissolution. *Current Biology: CB*, *26*(8), 1043–1050. https://doi.org/10.1016/j.cub.2016.02.010

Thomas Yeo, B. T., Krienen, F. M., Sepulcre, J., Sabuncu, M. R., Lashkari, D., Hollinshead, M., Roffman, J. L., Smoller, J. W., Zöllei, L., Polimeni, J. R., Fischl, B., Liu, H., & Buckner, R. L. (2011). The organization of the human cerebral cortex estimated by intrinsic functional connectivity. *Journal of Neurophysiology*, *106*(3), 1125–1165. https://doi.org/10.1152/jn.00338.2011

Tian, Y., Margulies, D. S., Breakspear, M., & Zalesky, A. (2020). Topographic organization of the human subcortex unveiled with functional connectivity gradients. *Nature Neuroscience*, *23*(11), Article 11. https://doi.org/10.1038/s41593-020-00711-6

Timmermann, C., Roseman, L., Haridas, S., Rosas, F. E., Luan, L., Kettner, H., Martell, J., Erritzoe, D., Tagliazucchi, E., Pallavicini, C., Girn, M., Alamia, A., Leech, R., Nutt, D. J., & Carhart-Harris, R. L. (2023). Human brain effects of DMT assessed via EEG-fMRI. *Proceedings of the*

*National Academy of Sciences of the United States of America*, *120*(13), e2218949120. https://doi.org/10.1073/pnas.2218949120

Timmermann, C., Roseman, L., Schartner, M., Milliere, R., Williams, L. T. J., Erritzoe, D., Muthukumaraswamy, S., Ashton, M., Bendrioua, A., Kaur, O., Turton, S., Nour, M. M., Day, C. M., Leech, R., Nutt, D. J., & Carhart-Harris, R. L. (2019). Neural correlates of the DMT experience assessed with multivariate EEG. *Scientific Reports*, *9*(1), Article 1. https://doi.org/10.1038/s41598-019-51974-4

Tipton, E., & Pustejovsky, J. E. (2015). Small-Sample Adjustments for Tests of Moderators and Model Fit Using Robust Variance Estimation in Meta-Regression. *Journal of Educational and Behavioral Statistics*, *40*(6), 604–634. https://doi.org/10.3102/1076998615606099

Turkeltaub, P. E., Eickhoff, S. B., Laird, A. R., Fox, M., Wiener, M., & Fox, P. (2011). Minimizing within-experiment and within-group effects in activation likelihood estimation meta-analyses. *Human Brain Mapping*, *33*(1), 1–13. https://doi.org/10.1002/hbm.21186

Umbricht, D., Koller, R., Vollenweider, F. X., & Schmid, L. (2002). Mismatch negativity predicts psychotic experiences induced by NMDA receptor antagonist in healthy volunteers. *Biological Psychiatry*, *51*(5), 400–406. https://doi.org/10.1016/s0006-3223(01)01242-2

Van Essen, D. C., Smith, S. M., Barch, D. M., Behrens, T. E. J., Yacoub, E., & Ugurbil, K. (2013). The WU-Minn Human Connectome Project: An overview. *NeuroImage*, *80*, 62–79. https://doi.org/10.1016/j.neuroimage.2013.05.041

Varley, T. F., Carhart-Harris, R., Roseman, L., Menon, D. K., & Stamatakis, E. A. (2020). Serotonergic psychedelics LSD & psilocybin increase the fractal dimension of cortical brain activity in spatial and temporal domains. *NeuroImage*, *220*, 117049. https://doi.org/10.1016/j.neuroimage.2020.117049

Veroniki, A. A., Jackson, D., Viechtbauer, W., Bender, R., Bowden, J., Knapp, G., Kuss, O., Higgins, J. P. T., Langan, D., & Salanti, G. (2016). Methods to estimate the between-study variance and its uncertainty in meta-analysis. *Research Synthesis Methods*, *7*(1), 55–79. https://doi.org/10.1002/jrsm.1164

Viol, A., Palhano-Fontes, F., Onias, H., de Araujo, D. B., Hövel, P., & Viswanathan, G. M. (2019). Characterizing Complex Networks Using Entropy-Degree Diagrams: Unveiling Changes in Functional Brain Connectivity Induced by Ayahuasca. *Entropy*, *21*(2), Article 2. https://doi.org/10.3390/e21020128

Viol, A., Palhano-Fontes, F., Onias, H., de Araujo, D. B., & Viswanathan, G. M. (2017). Shannon entropy of brain functional complex networks under the influence of the psychedelic Ayahuasca. *Scientific Reports*, *7*(1), Article 1. https://doi.org/10.1038/s41598-017-06854-0

Viol, A., Viswanathan, G. M., Soldatkina, O., Palhano-Fontes, F., Onias, H., Araujo, D. de, & Hövel, P. (2023). Information parity increases on functional brain networks under influence of a psychedelic substance. *Journal of Physics: Complexity*, *4*(1), 01LT02. https://doi.org/10.1088/2632-072X/acc22b

Vogt, S. B., Ley, L., Erne, L., Straumann, I., Becker, A. M., Klaiber, A., Holze, F., Vandersmissen, A., Mueller, L., Duthaler, U., Rudin, D., Luethi, D., Varghese, N., Eckert, A., & Liechti, M. E. (2023). Acute effects of intravenous DMT in a randomized placebo-controlled study in healthy participants. *Translational Psychiatry*, *13*(1), 172. https://doi.org/10.1038/s41398-023-02477-4

Vollenweider, F. X., Csomor, P. A., Knappe, B., Geyer, M. A., & Quednow, B. B. (2007). The effects of the preferential 5-HT2A agonist psilocybin on prepulse inhibition of startle in healthy human volunteers depend on interstimulus interval. *Neuropsychopharmacology: Official Publication of the American College of Neuropsychopharmacology*, *32*(9), 1876–1887. https://doi.org/10.1038/sj.npp.1301324

Wen, A., Singhal, N., Jones, B. D. M., Zeifman, R. J., Mehta, S., Shenasa, M. A., Blumberger, D. M., Daskalakis, Z. J., & Weissman, C. R. (2023). A Systematic Review of Study Design and Placebo Controls in Psychedelic Research. *Psychedelic Medicine*. https://doi.org/10.1089/psymed.2023.0028

Wießner, I., Falchi, M., Palhano-Fontes, F., Feilding, A., Ribeiro, S., & Tófoli, L. F. (2023). LSD, madness and healing: Mystical experiences as possible link between psychosis model and therapy model. *Psychological Medicine*, *53*(4), 1151–1165. https://doi.org/10.1017/S0033291721002531

Wittmann, M., Carter, O., Hasler, F., Cahn, B. R., Grimberg, U., Spring, P., Hell, D., Flohr, H., & Vollenweider, F. X. (2007). Effects of psilocybin on time perception and temporal control of behaviour in humans. *Journal of Psychopharmacology*, *21*(1), 50–64. https://doi.org/10.1177/0269881106065859

Zalesky, A., Fornito, A., & Bullmore, E. T. (2010). Network-based statistic: Identifying differences in brain networks. *NeuroImage*, *53*(4), 1197–1207. https://doi.org/10.1016/j.neuroimage.2010.06.041

Zamberlan, F., Sanz, C., Martínez Vivot, R., Pallavicini, C., Erowid, F., Erowid, E., & Tagliazucchi, E. (2018). The Varieties of the Psychedelic Experience: A Preliminary Study of the Association Between the Reported Subjective Effects and the Binding Affinity Profiles of Substituted Phenethylamines and Tryptamines. *Frontiers in Integrative Neuroscience*, *12*, 54. https://doi.org/10.3389/fnint.2018.00054

Ziv, J., & Lempel, A. (1977). A universal algorithm for sequential data compression. *IEEE Transactions on Information Theory*, *23*(3), 337–343. IEEE Transactions on Information Theory. https://doi.org/10.1109/TIT.1977.1055714

Ziv, J., & Lempel, A. (1978). Compression of individual sequences via variable-rate coding. *IEEE Transactions on Information Theory*, *24*(5), 530–536. IEEE Transactions on Information Theory. https://doi.org/10.1109/TIT.1978.1055934

### Supplementary Figures and Tables

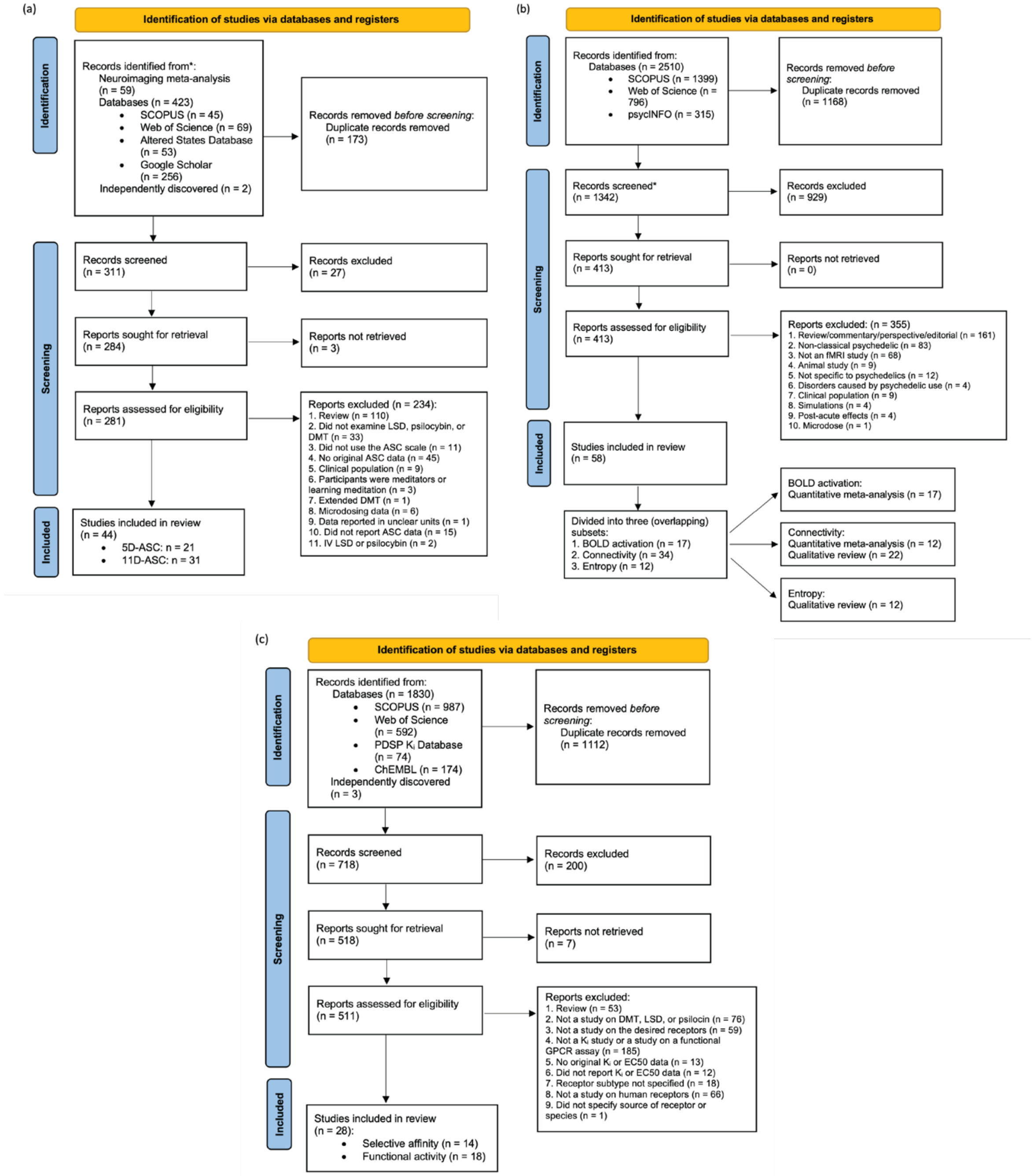

***Figure S1. Literature search flowcharts.*** *These diagrams, created from the PRISMA 2020 flow diagram template (Page et al., 2021), describe the process of identifying the (a) phenomenology studies; (b) neuroimaging studies, and (c) pharmacology studies. A combination of manual and automated techniques were used to remove duplicate records. Screening resulted in the manual deletion of irrelevant papers, such as those that abbreviated "least squares difference" as LSD.*

(a)

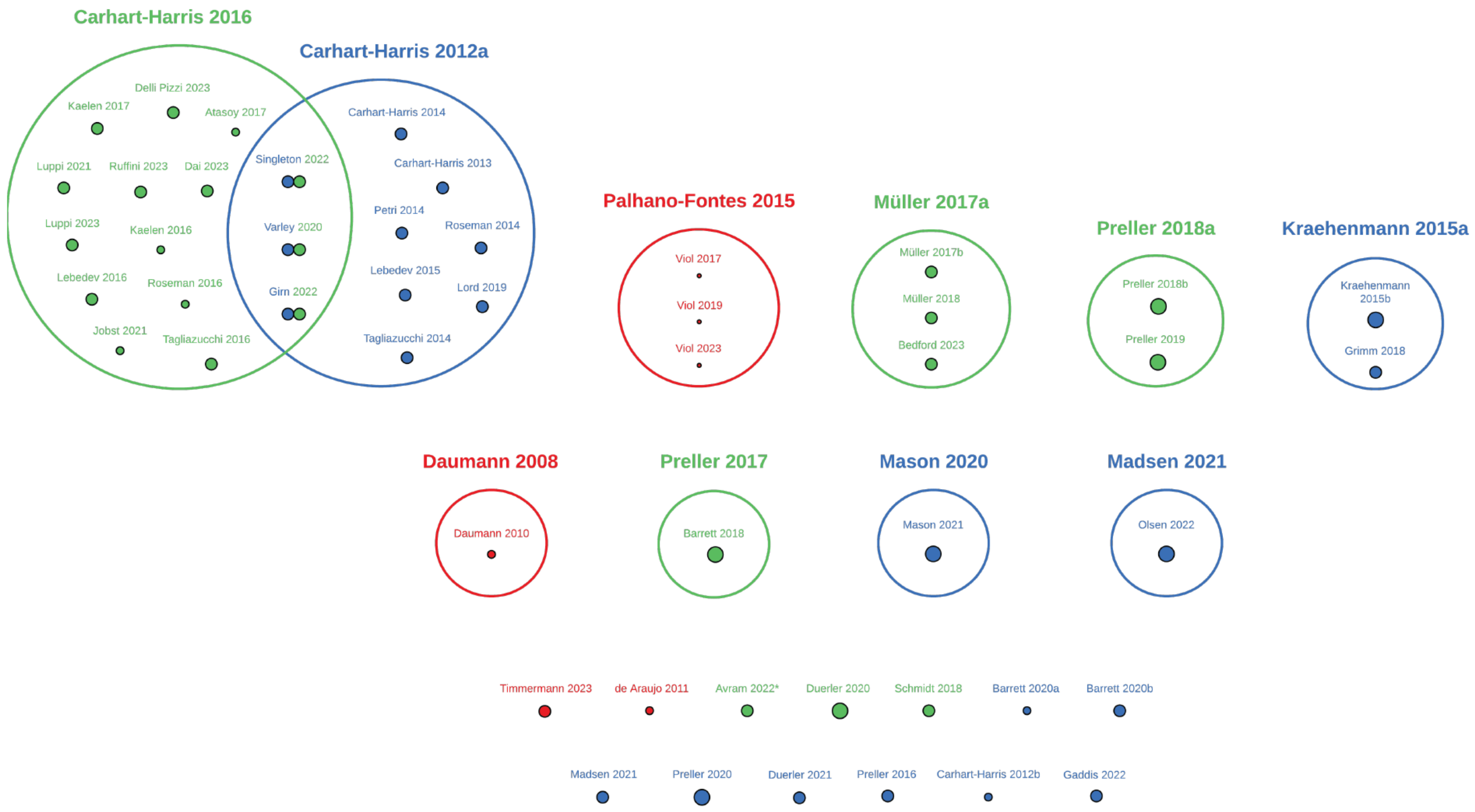

*Avram et al. (2022) does not contain a primary dataset; its fMRI data was collected by Holze et al. (2020), which is not included in this meta-analysis.

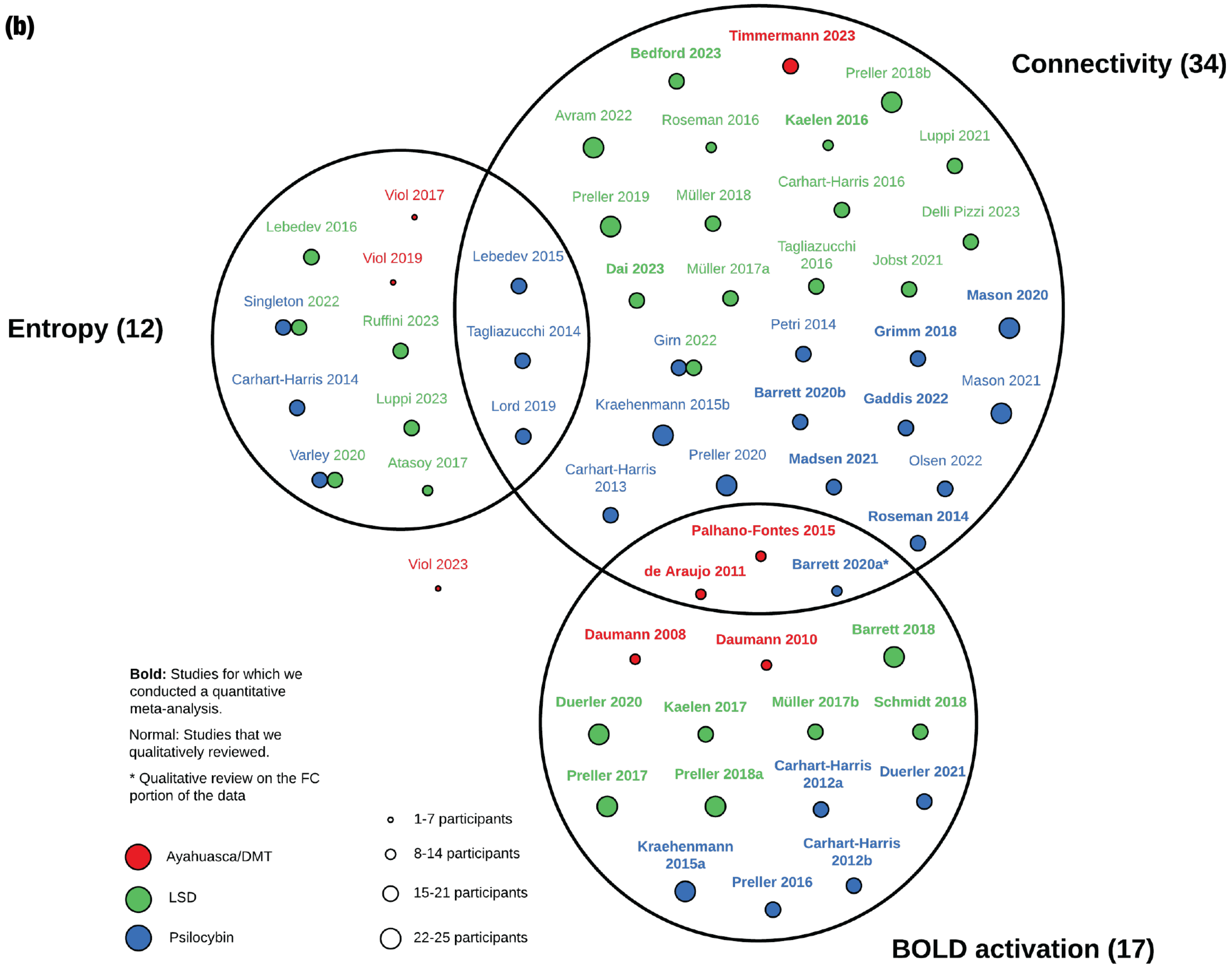
(b)
Connectivity (34)
Entropy (12)
BOLD activation (17)
Timmermann 2023
Bedford 2023
Preller 2018b
Avram 2022
Roseman 2016
Kaelen 2016
Luppi 2021
Carhart-Harris 2016
Preller 2019
Müller 2018
Delli Pizzi 2023
Tagliazucchi 2016
Jobst 2021
Dai 2023
Müller 2017a
Mason 2020
Petri 2014
Grimm 2018
Girn 2022
Mason 2021
Barrett 2020b
Gaddis 2022
Kraehenmann 2015b
Preller 2020
Madsen 2021
Olsen 2022
Carhart-Harris 2013
Roseman 2014
Palhano-Fontes 2015
Barrett 2020a*
de Araujo 2011
Viol 2017
Lebedev 2016
Viol 2019
Lebedev 2015
Singleton 2022
Ruffini 2023
Tagliazucchi 2014
Carhart-Harris 2014
Luppi 2023
Lord 2019
Varley 2020
Atasoy 2017
Viol 2023
Daumann 2008
Daumann 2010
Barrett 2018
Duerler 2020
Kaelen 2017
Müller 2017b
Schmidt 2018
Preller 2017
Preller 2018a
Carhart-Harris 2012a
Duerler 2021
Kraehenmann 2015a
Carhart-Harris 2012b
Preller 2016
Bold: Studies for which we conducted a quantitative meta-analysis.
Normal: Studies that we qualitatively reviewed.
* Qualitative review on the FC portion of the data
Ayahuasca/DMT
LSD
Psilocybin
1-7 participants
8-14 participants
15-21 participants
22-25 participants

***Fig S2. Overview of the fMRI literature on psychedelics.*** *(a) Primary datasets vs. secondary analyses. Primary datasets with at least one secondary analysis are depicted as large circles containing smaller circles. Clearly, the primary datasets in Carhart-Harris et al. (2016) and Carhart-Harris, Erritzoe, et al. (2012) received disproportionately more secondary analyses than any other primary datasets. (b) Venn diagram of the drugs, number of participants, and metric of brain activity (entropy, connectivity, or BOLD activation) analysed by each study. The size of the colored circles represents the number of participants in the study. We conducted a qualitative review of all the entropy studies, any FC studies for which the data was unavailable, and any FC studies whose analysis methods were not suitable for our quantitative meta-analysis. Note that while we did conduct a quantitative analysis of the BOLD activation studies, we did not ultimately report the results because we thought they would be misleading (see* Section 2.2*). Daumann et al. (2008, 2010) and Timmermann et al. (2023) used DMT, whereas the remainder of the studies in the ayahuasca/DMT category measured the effects of ayahuasca on brain activity. Barrett 2020a refers to Barrett, Doss, et al. (2020); Barrett 2020b to Barrett, Krimmel, et al. (2020); Müller 2017a to Müller, Lenz, Dolder, Lang, et al. (2017) and Müller 2017b to Müller, Lenz, Dolder, Harder, et al. (2017); Preller 2018a to Preller, Schilbach, et al. (2018); Preller 2018b to Preller, Burt, et al. (2018); Carhart-Harris 2012a to Carhart-Harris, Erritzoe, et al. (2012); Carhart-Harris 2012b to Carhart-Harris, Leech, et al. (2012).*

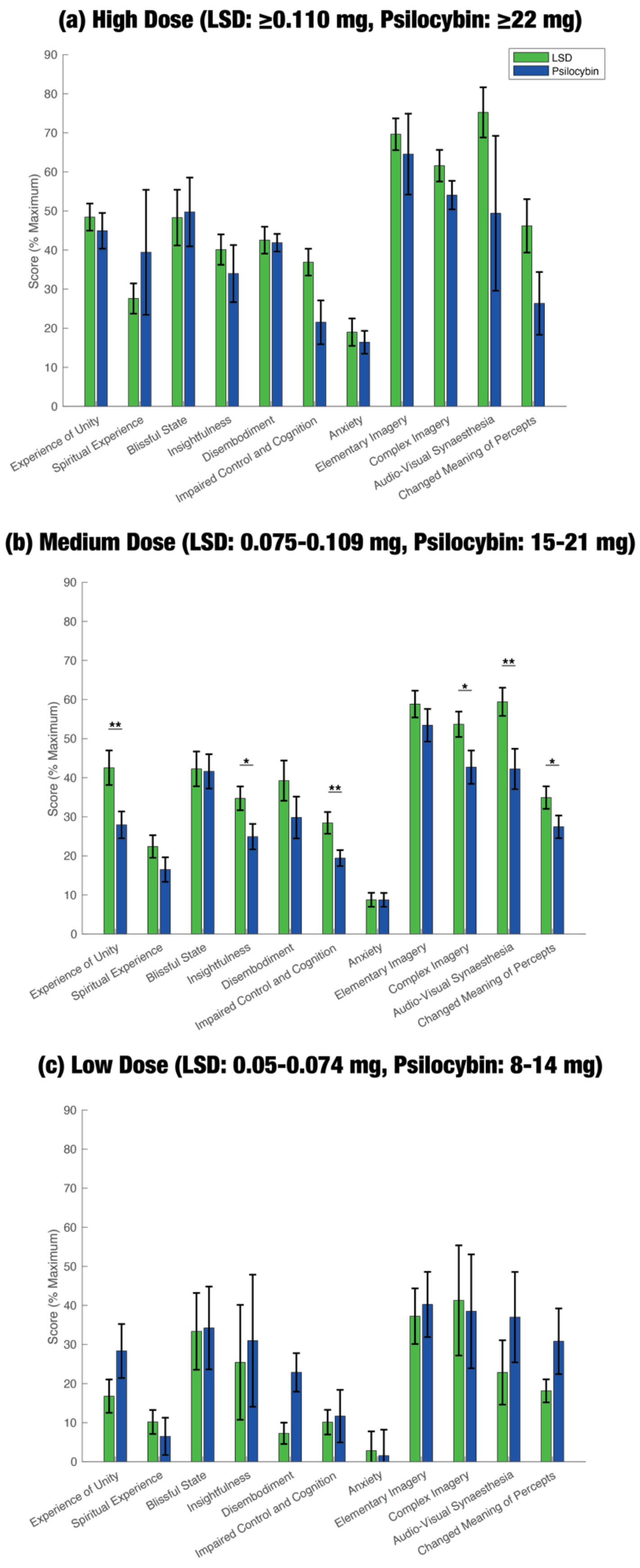
(a) High Dose (LSD: ≥0.110 mg, Psilocybin: ≥22 mg)
LSD
Psilocybin
Score (% Maximum)
0
10
20
30
40
50
60
70
80
90
Experience of Unity
Spiritual Experience
Blissful State
Insightfulness
Disembodiment
Impaired Control and Cognition
Anxiety
Elementary Imagery
Complex Imagery
Audio-Visual Synaesthesia
Changed Meaning of Percepts
(b) Medium Dose (LSD: 0.075-0.109 mg, Psilocybin: 15-21 mg)
Score (% Maximum)
**
*
**
*
**
*
(c) Low Dose (LSD: 0.05-0.074 mg, Psilocybin: 8-14 mg)
Score (% Maximum)

**(d) High Dose (LSD: ≥0.110 mg, Psilocybin: ≥22 mg)**

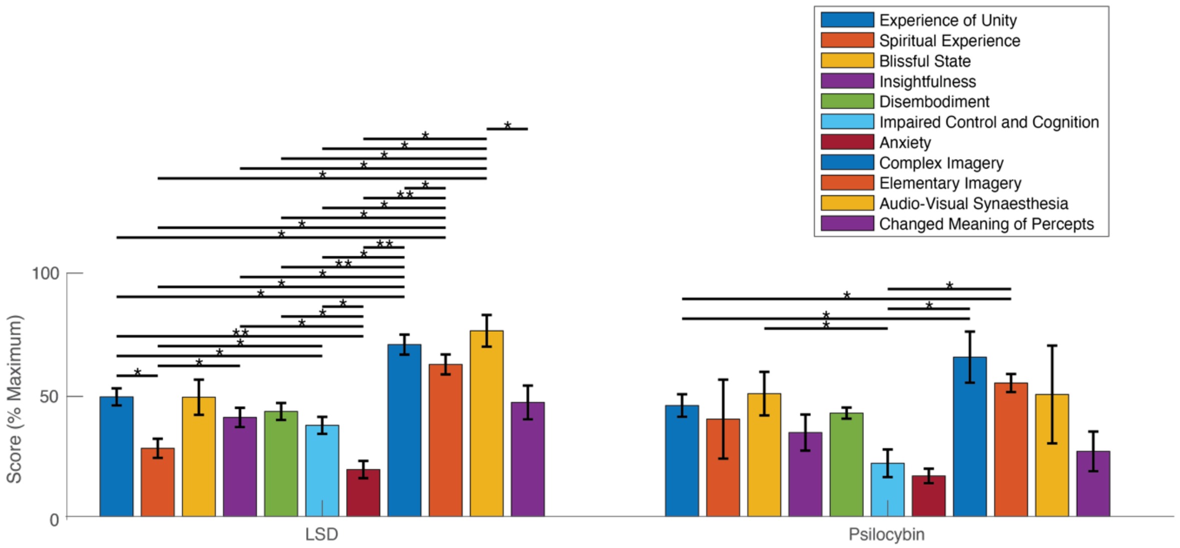

**(e) Medium Dose (LSD: 0.075-0.109 mg, Psilocybin: 15-21 mg)**

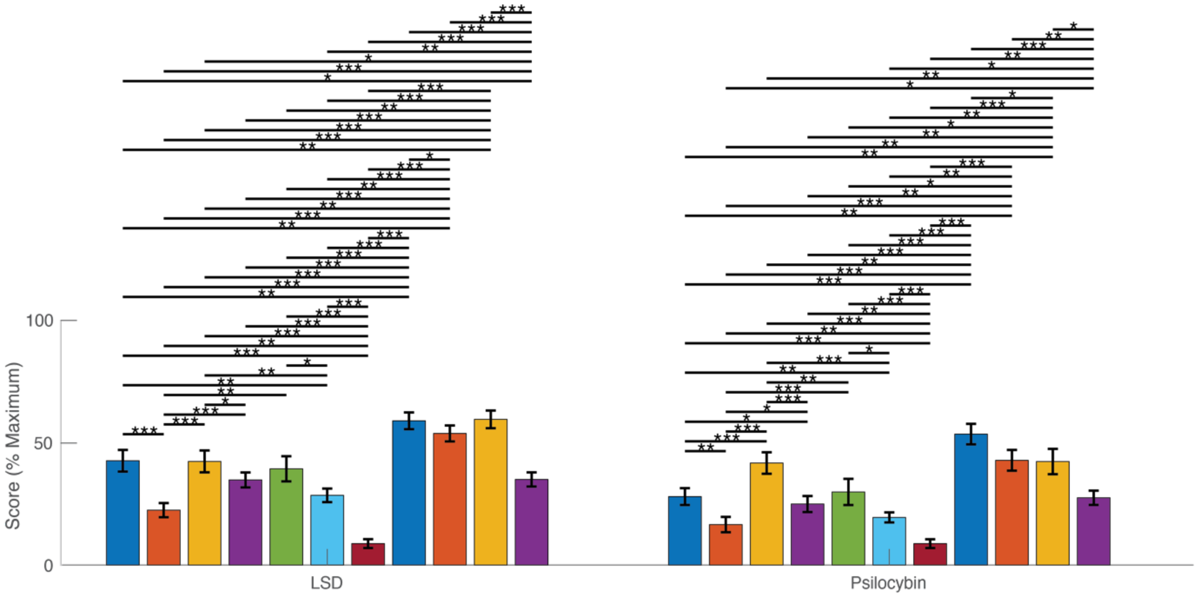

**(f) Low Dose (LSD: 0.05-0.074 mg, Psilocybin: 8-14 mg)**

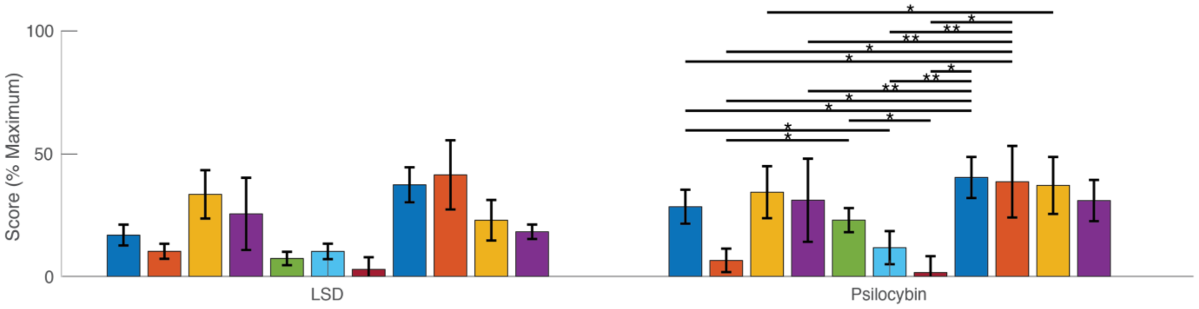

***Fig S3. Meta-analysis of the 11-dimensional Altered States of Consciousness (ASC) data.*** *Our literature search identified 33 studies that reported 11D-ASC data. We performed a multilevel random-effects meta-analysis to account for the lack of statistical independence between measurements of different dimensions in the same group of participants. Within-study and between-study heterogeneity were estimated with the restricted maximum likelihood procedure. Using a Correlated and Hierarchical Effects model to address within-study correlations in sampling error, we analysed the effect of the interaction between dose, dimension, and drug (LSD and psilocybin) on pooled 11D-ASC scores. Note that we could not include DMT in this meta-analysis as there was only one 11D-ASC study on DMT that reported standard errors. We only compared the two drugs for similar doses, assuming that 0.1 mg LSD = 20 mg psilocybin (Ley et al., 2023). (a-c) Between-drug comparisons of pooled 11D-ASC scores. LSD tended to rank higher than psilocybin in the subjective dimensions, except for low doses; however, there was only one study that gave LSD at a low dose (0.005 mg), so our pooled estimates may be unreliable. Differences only reached significance for medium doses, in two subscales of oceanic boundlessness (experience of unity and insightfulness), one subscale of anxious ego dissolution (impaired control and cognition), and three subscales of visionary restructuralisation (complex imagery, audio-visual synaesthesia, and changed meaning of percepts). (d-f) Within-drug comparisons. Subscales of visionary restructuralisation (complex imagery, elementary imagery, audio-visual synaesthesia, changed meaning of percepts), tended to be significantly greater than subscales of anxious ego dissolution (disembodiment, impaired control and cognition, anxiety) and oceanic boundlessness (experience of unity, spiritual experience, blissful state, insightfulness). Numerical data for the above figure are given in* ***Table S2****.* * $p < 0.05$, ** $p < 0.01$, *** $p < 0.001$.

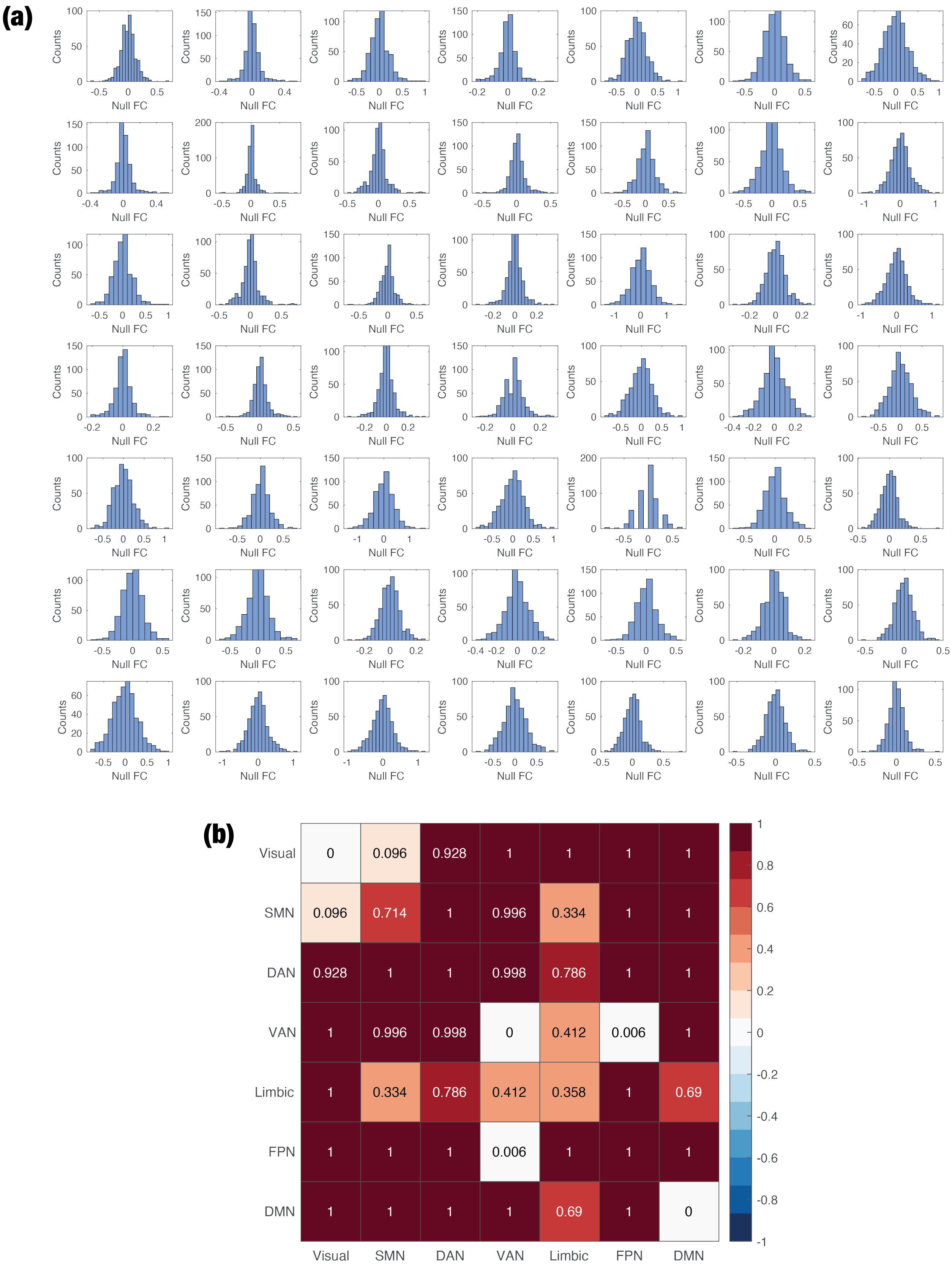
(a)
Counts
Null FC
(b)
Visual 0 0.096 0.928 1 1 1 1
SMN 0.096 0.714 1 0.996 0.334 1 1
DAN 0.928 1 1 0.998 0.786 1 1
VAN 1 0.996 0.998 0 0.412 0.006 1
Limbic 1 0.334 0.786 0.412 0.358 1 0.69
FPN 1 1 1 0.006 1 1 1
DMN 1 1 1 1 0.69 1 0
Visual SMN DAN VAN Limbic FPN DMN
1 0.8 0.6 0.4 0.2 0 -0.2 -0.4 -0.6 -0.8 -1

***Figure S4. Null distribution of functional connectivity (FC).*** *Null distribution was formed by measuring functional connectivity in resting-state fMRI data collected by the Human Connectome Project (HCP) in 100 unrelated, healthy, sober individuals. The data was parcellated with the Schaefer-100 atlas. The FC meta-analysis algorithm was repeatedly applied to randomly chosen subjects and windows of data, resulting in a "surrogate study set." (a) We formed 500 surrogate study sets and produced a null distribution of each element of the aggregate FC matrix. (b) We determined the percentile of each element of the true aggregate FC matrix within the null distribution. If an entry of the true aggregate FC matrix was below the 2.5$^{th}$ percentile or above the 97.5$^{th}$ percentile of the corresponding entry in the null distribution, then it was deemed significant (so long as it also survived multiple comparisons.)*

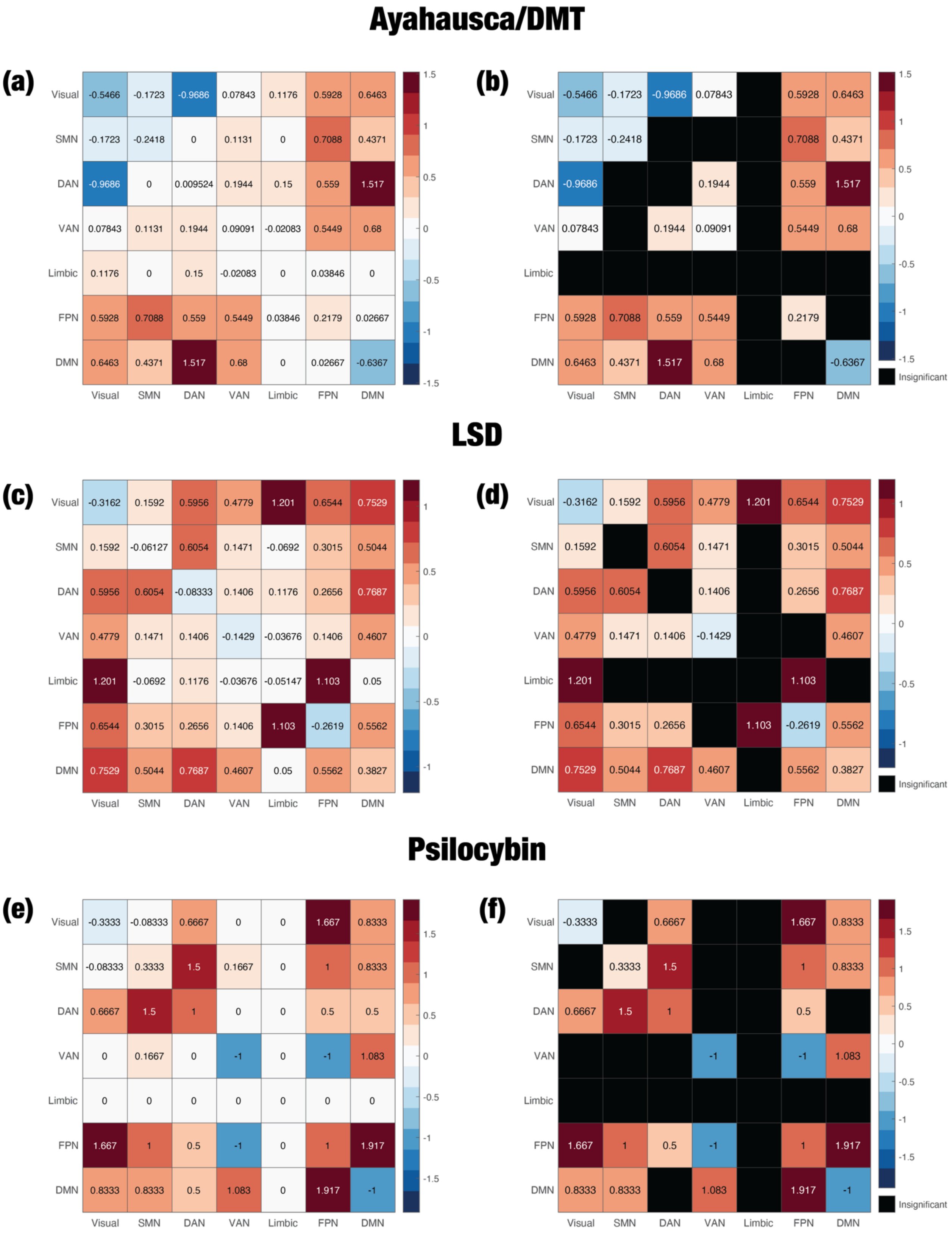
Ayahausca/DMT
(a)
Visual -0.5466 -0.1723 -0.9686 0.07843 0.1176 0.5928 0.6463
SMN -0.1723 -0.2418 0 0.1131 0 0.7088 0.4371
DAN -0.9686 0 0.009524 0.1944 0.15 0.559 1.517
VAN 0.07843 0.1131 0.1944 0.09091 -0.02083 0.5449 0.68
Limbic 0.1176 0 0.15 -0.02083 0 0.03846 0
FPN 0.5928 0.7088 0.559 0.5449 0.03846 0.2179 0.02667
DMN 0.6463 0.4371 1.517 0.68 0 0.02667 -0.6367
Visual SMN DAN VAN Limbic FPN DMN
(b)
Visual -0.5466 -0.1723 -0.9686 0.07843 0.5928 0.6463
SMN -0.1723 -0.2418 0.7088 0.4371
DAN -0.9686 0.1944 0.559 1.517
VAN 0.07843 0.1944 0.09091 0.5449 0.68
Limbic
FPN 0.5928 0.7088 0.559 0.5449 0.2179
DMN 0.6463 0.4371 1.517 0.68 -0.6367
Visual SMN DAN VAN Limbic FPN DMN
Insignificant
LSD
(c)
Visual -0.3162 0.1592 0.5956 0.4779 1.201 0.6544 0.7529
SMN 0.1592 -0.06127 0.6054 0.1471 -0.0692 0.3015 0.5044
DAN 0.5956 0.6054 -0.08333 0.1406 0.1176 0.2656 0.7687
VAN 0.4779 0.1471 0.1406 -0.1429 -0.03676 0.1406 0.4607
Limbic 1.201 -0.0692 0.1176 -0.03676 -0.05147 1.103 0.05
FPN 0.6544 0.3015 0.2656 0.1406 1.103 -0.2619 0.5562
DMN 0.7529 0.5044 0.7687 0.4607 0.05 0.5562 0.3827
Visual SMN DAN VAN Limbic FPN DMN
(d)
Visual -0.3162 0.1592 0.5956 0.4779 1.201 0.6544 0.7529
SMN 0.1592 0.6054 0.1471 0.3015 0.5044
DAN 0.5956 0.6054 0.1406 0.2656 0.7687
VAN 0.4779 0.1471 0.1406 -0.1429 0.4607
Limbic 1.201 1.103
FPN 0.6544 0.3015 0.2656 1.103 -0.2619 0.5562
DMN 0.7529 0.5044 0.7687 0.4607 0.5562 0.3827
Visual SMN DAN VAN Limbic FPN DMN
Insignificant
Psilocybin
(e)
Visual -0.3333 -0.08333 0.6667 0 0 1.667 0.8333
SMN -0.08333 0.3333 1.5 0.1667 0 1 0.8333
DAN 0.6667 1.5 1 0 0 0.5 0.5
VAN 0 0.1667 0 -1 0 -1 1.083
Limbic 0 0 0 0 0 0 0
FPN 1.667 1 0.5 -1 0 1 1.917
DMN 0.8333 0.8333 0.5 1.083 0 1.917 -1
Visual SMN DAN VAN Limbic FPN DMN
(f)
Visual -0.3333 0.6667 1.667 0.8333
SMN 0.3333 1.5 1 0.8333
DAN 0.6667 1.5 1 0.5
VAN -1 -1 1.083
Limbic
FPN 1.667 1 0.5 -1 1 1.917
DMN 0.8333 0.8333 1.083 1.917 -1
Visual SMN DAN VAN Limbic FPN DMN
Insignificant

***Figure S5. Aggregate FC matrices for each psychedelic.*** *Whereas* ***Figure 3*** *displays the aggregate FC matrix across all three psychedelics, here we show the aggregate FC matrix that was computed on each individual psychedelic. The left column displays all of the aggregate FC values, including insignificant edges, whereas the right column shows only significant entries. All psychedelics significantly disintegrate the visual network and generally increase between-network FC, with a couple of exceptions for ayahuasca/DMT and psilocybin.*

(a)

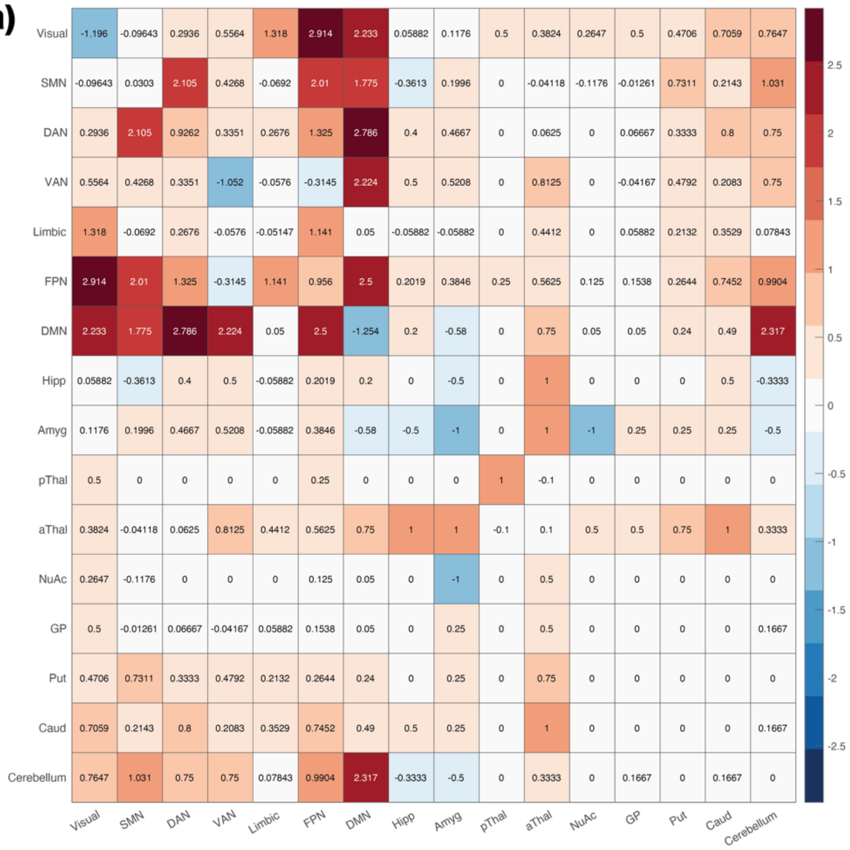
Visual -1.196 -0.09643 0.2936 0.5564 1.318 2.914 2.233 0.05882 0.1176 0.5 0.3824 0.2647 0.5 0.4706 0.7059 0.7647
SMN -0.09643 0.0303 2.105 0.4268 -0.0692 2.01 1.775 -0.3613 0.1996 0 -0.04118 -0.1176 -0.01261 0.7311 0.2143 1.031
DAN 0.2936 2.105 0.9262 0.3351 0.2676 1.325 2.786 0.4 0.4667 0 0.0625 0 0.06667 0.3333 0.8 0.75
VAN 0.5564 0.4268 0.3351 -1.052 -0.0576 -0.3145 2.224 0.5 0.5208 0 0.8125 0 -0.04167 0.4792 0.2083 0.75
Limbic 1.318 -0.0692 0.2676 -0.0576 -0.05147 1.141 0.05 -0.05882 -0.05882 0 0.4412 0 0.05882 0.2132 0.3529 0.07843
FPN 2.914 2.01 1.325 -0.3145 1.141 0.956 2.5 0.2019 0.3846 0.25 0.5625 0.125 0.1538 0.2644 0.7452 0.9904
DMN 2.233 1.775 2.786 2.224 0.05 2.5 -1.254 0.2 -0.58 0 0.75 0.05 0.05 0.24 0.49 2.317
Hipp 0.05882 -0.3613 0.4 0.5 -0.05882 0.2019 0.2 0 -0.5 0 1 0 0 0 0.5 -0.3333
Amyg 0.1176 0.1996 0.4667 0.5208 -0.05882 0.3846 -0.58 -0.5 -1 0 1 -1 0.25 0.25 0.25 -0.5
pThal 0.5 0 0 0 0 0.25 0 0 0 1 -0.1 0 0 0 0 0
aThal 0.3824 -0.04118 0.0625 0.8125 0.4412 0.5625 0.75 1 1 -0.1 0.1 0.5 0.5 0.75 1 0.3333
NuAc 0.2647 -0.1176 0 0 0 0.125 0.05 0 -1 0 0.5 0 0 0 0 0
GP 0.5 -0.01261 0.06667 -0.04167 0.05882 0.1538 0.05 0 0.25 0 0.5 0 0 0 0 0.1667
Put 0.4706 0.7311 0.3333 0.4792 0.2132 0.2644 0.24 0 0.25 0 0.75 0 0 0 0 0
Caud 0.7059 0.2143 0.8 0.2083 0.3529 0.7452 0.49 0.5 0.25 0 1 0 0 0 0 0.1667
Cerebellum 0.7647 1.031 0.75 0.75 0.07843 0.9904 2.317 -0.3333 -0.5 0 0.3333 0 0.1667 0 0.1667 0
Visual SMN DAN VAN Limbic FPN DMN Hipp Amyg pThal aThal NuAc GP Put Caud Cerebellum
2.5 2 1.5 1 0.5 0 -0.5 -1 -1.5 -2 -2.5

(b)

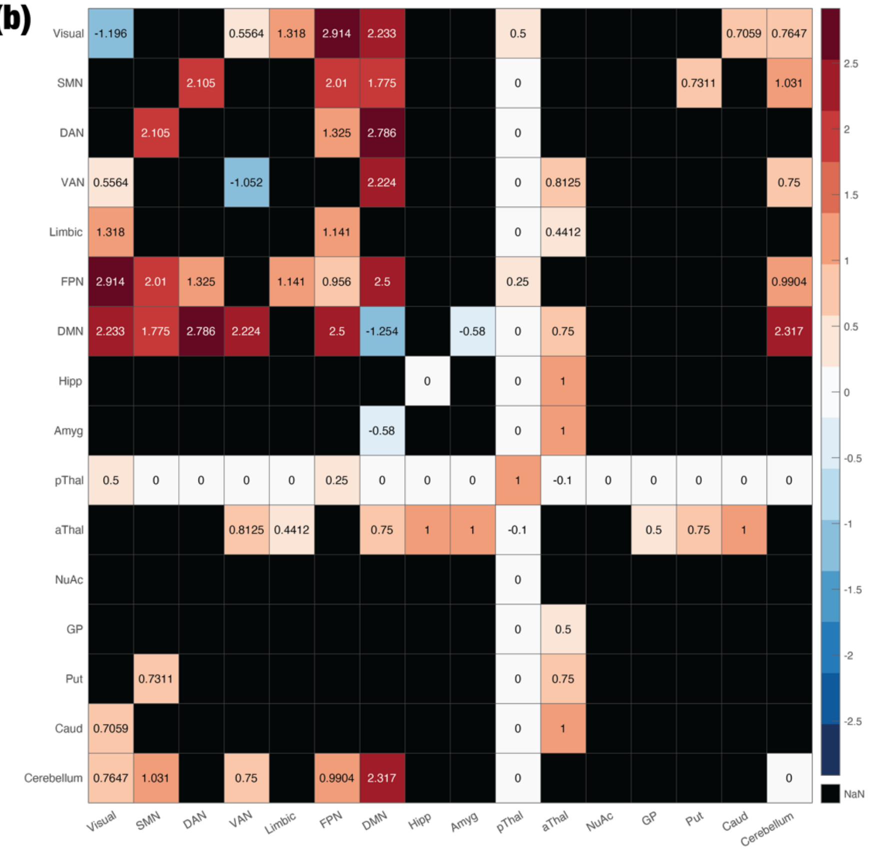
Visual -1.196 0.5564 1.318 2.914 2.233 0.5 0.7059 0.7647
SMN 2.105 2.01 1.775 0 0.7311 1.031
DAN 2.105 1.325 2.786 0
VAN 0.5564 -1.052 2.224 0 0.8125 0.75
Limbic 1.318 1.141 0 0.4412
FPN 2.914 2.01 1.325 1.141 0.956 2.5 0.25 0.9904
DMN 2.233 1.775 2.786 2.224 2.5 -1.254 -0.58 0 0.75 2.317
Hipp 0 0 1
Amyg -0.58 0 1
pThal 0.5 0 0 0 0 0.25 0 0 0 1 -0.1 0 0 0 0 0
aThal 0.8125 0.4412 0.75 1 1 -0.1 0.5 0.75 1
NuAc 0
GP 0 0.5
Put 0.7311 0 0.75
Caud 0.7059 0 1
Cerebellum 0.7647 1.031 0.75 0.9904 2.317 0 0
Visual SMN DAN VAN Limbic FPN DMN Hipp Amyg pThal aThal NuAc GP Put Caud Cerebellum
2.5 2 1.5 1 0.5 0 -0.5 -1 -1.5 -2 -2.5
NaN

***Figure S6. Aggregate FC matrix with subcortical regions included.*** *(a) We measured the aggregate FC matrix using the Tian parcellation, which defines eight subcortical regions: hippocampus (Hipp), posterior thalamus (pThal), anterior thalamus (aThal), nucleus accumbens (NuAc), globus pallidus (GP), putamen (Put), caudate (Caud), and cerebellum (Tian et al., 2020). Note that the cortical portion of this matrix is the same as* ***Figure 3****. (b) The statistical significance of this matrix was determined relative to a null distribution formed from resting-state HCP data. Unlike in the cortical case, the resting-state HCP data was parcellated here with the Desikan-Killiany (DK-80) atlas, which includes subcortical regions. (Due to the change in parcellation, the null distribution of the cortical FC is also altered.) FC between most pairs of subcortical regions, as well as between subcortex and cortex, was generally deemed insignificant, likely due to a lack of subcortical FC analyses in the literature. The increased connectivity between the anterior thalamus and cortex, as well as between the cerebellum and cortex, stand out as exceptions.*

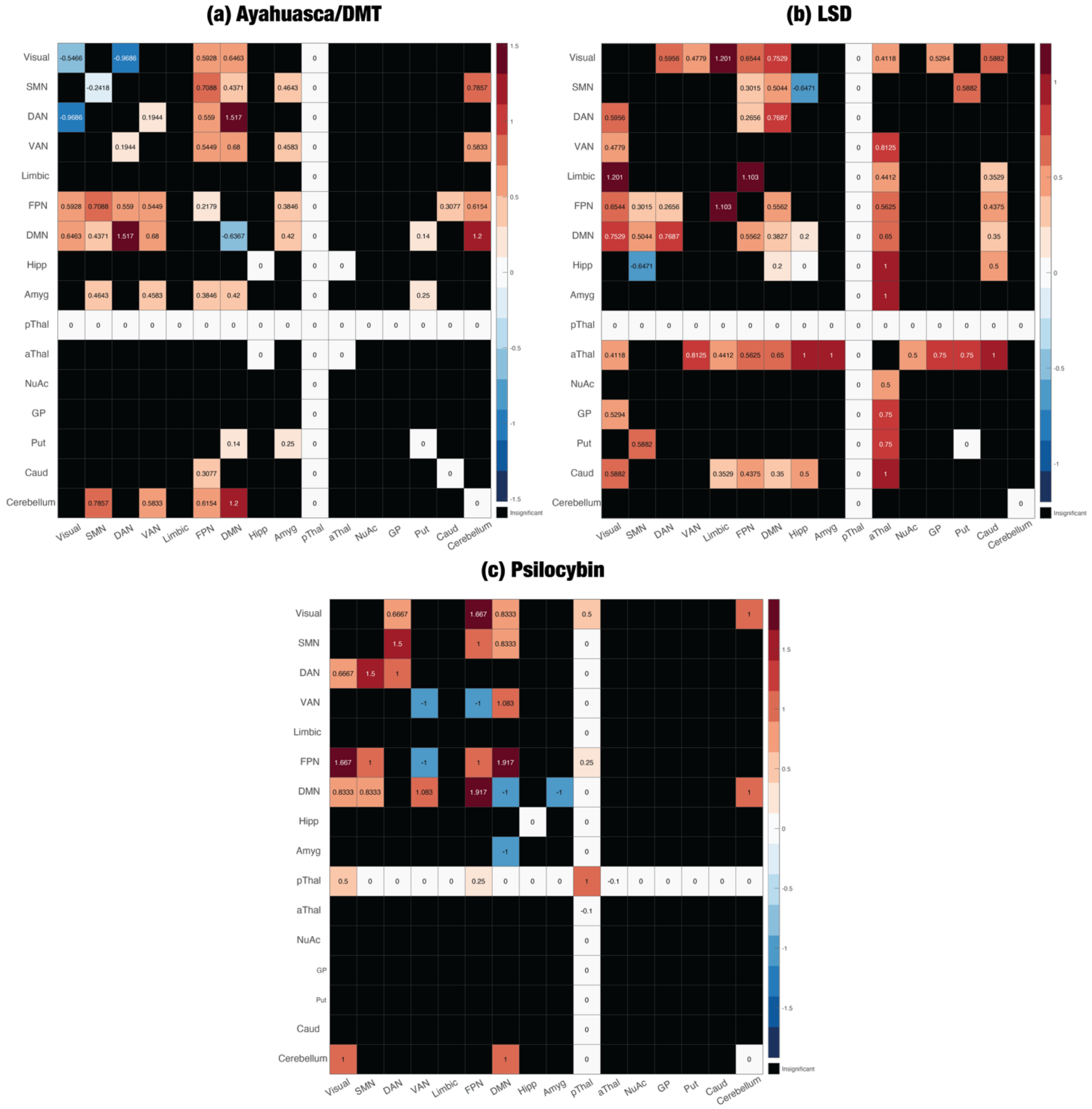

***Figure S7. Aggregate FC matrices for each psychedelic, with subcortical regions included.*** *FC between subcortical regions, as well as FC between cortical and subcortical regions, tended to be insignificant for each psychedelic, especially psilocybin. LSD significantly elevated the FC of the anterior thalamus, both to the subcortex and to the cortex.*

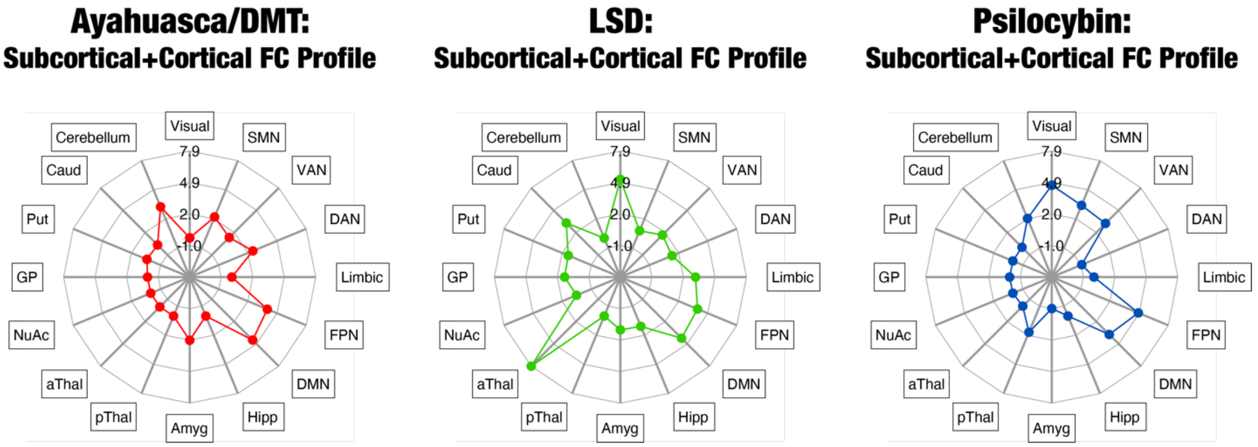

***Figure S8. Functional connectivity profiles with subcortical regions included.*** *We summed across the columns of each of the matrices in **Figure S7** to obtain the corresponding spider plots. Compared to the other psychedelics, LSD much more strongly elevated the total FC of the anterior thalamus and, to a lesser extent, the caudate, even though there was only a single study on the effects of LSD on subcortical FC. Ayahuasca/DMT enhanced the total FC of the amygdala and cerebellum, but psilocybin had a negligible effect on subcortical FC, likely due to the lack of subcortical analyses of psilocybin.*

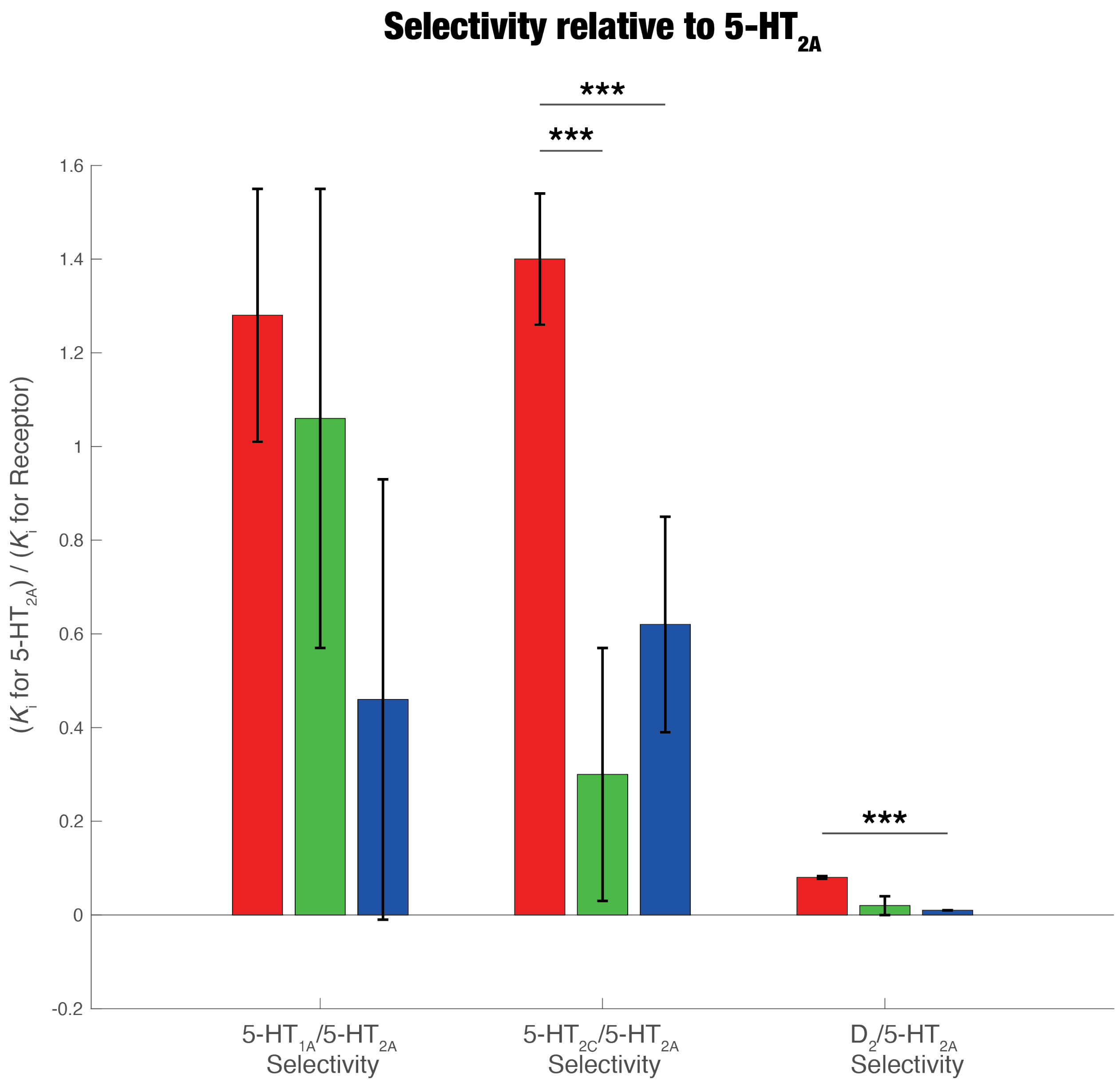

***Figure S9. While there were no significant between-drug differences in selectivity relative to the 5-HT$_{1A}$ receptor, there are significant differences relative to the 5-HT$_{2A}$ receptor.*** *We created random-effects models that modeled between-study variance. DMT was significantly more selective for the 5-HT$_{2C}$ receptor than LSD and psilocin. Additionally, DMT exhibited significantly higher selectivity for the $D_2$ receptor than psilocin.* *** $p < 0.001$, ** $p < 0.01$, * $p < 0.05$.

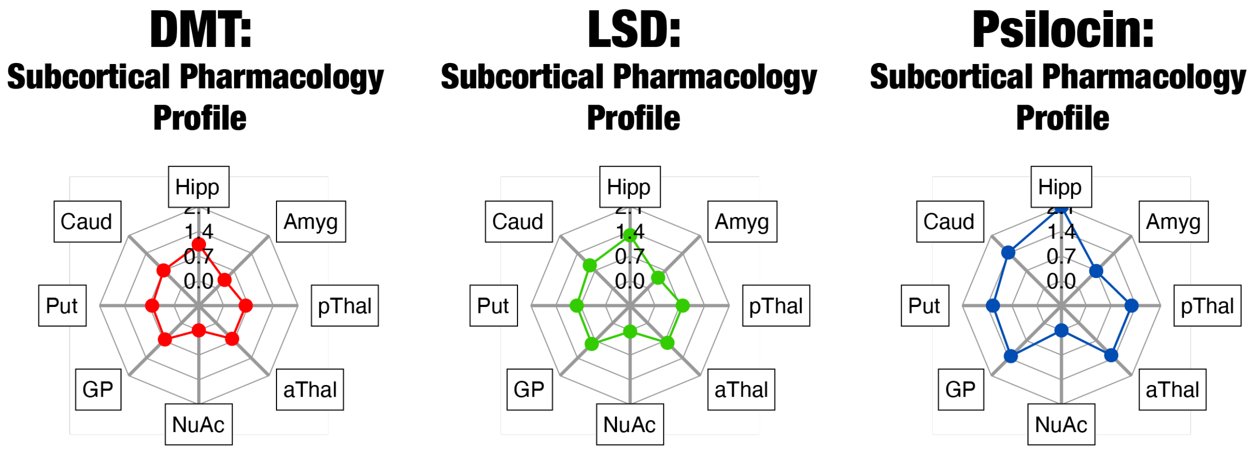

***Figure S10. Pharmacology profiles in subcortical regions.*** *We obtained subcortical maps of 5-HT$_{2A}$ (Beliveau et al., 2017) and D$_2$ receptor expression (Malén et al., 2022). The pharmacology profiles are a weighted sum of the receptor densities across the eight subcortical regions in the Tian atlas, in which the weights are the selectivities of each drug for the corresponding receptor. As in the cortical case, the profiles are dominated by 5-HT$_{2A}$ receptor expression, since the selectivity for 5-HT$_{2A}$ is two orders of magnitude higher than selectivity for D$_2$ across drugs.*

| | DMT | | LSD | | | Psilocybin | | |
|---|---|---|---|---|---|---|---|---|
| | Low Dose (10-15 mg bolus + 0.6-0.99 mg/min IV) | High Dose (≥16 mg bolus + ≥ 1 mg/min IV) | Low Dose (0.05-0.074 mg) | Medium Dose (0.075-0.109 mg) | High Dose (≥0.110 mg) | Low Dose (8-14 mg) | Medium Dose (15-21 mg) | High Dose (≥22 mg) |
| OB | 27.84 ± 2.89 | 36.35 ± 6.92 | 24.37 ± 9.97 | 47.13 ± 5.93 | 47.72 ± 2.55 | 16.77 ± 3.86 | 29.53 ± 2.45 | 41.96 ± 2.45 |
| AED | 9.79 ± 2.13 | 18.45 ± 6.84 | 11.36 ± 3.49 | 17.63 ± 0.93 | 27.85 ± 1.83 | 9.22 ± 1.17 | 15.07 ± 1.32 | 21.85 ± 1.97 |
| VR | 32.44 ± 3.5 | 39.43 ± 3.49 | 40.23 ± 13.11 | 45.31 ± 3.22 | 58.15 ± 2.52 | 15.76 ± 2.17 | 37.45 ± 3.17 | 47.59 ± 2.05 |
| AA | 7.3 ± 2.02 (*) | 10 ± 3.1 (*) | 10.40 ± 3.31 | 12.01 ± 1.17 | 20.02 ± 1.87 | 6.56 ± 1.1 | 9.89 ± 1.08 | 12.82 ± 1.4 |
| RV | 24 ± 4.1 (*) | 32 ± 4.4 (*) | 20.62 ± 5.54 | 25.20 ± 5.57 | 37.1 ± 4.9 | 20.18 ± 4.34 | 28.61 ± 1.7 | 34.96 ± 5.62 |

***Table S1. Numerical data for 5D-ASC meta-analysis.*** *Data given as percentage of maximum score. (*) means that there was only one study that measured the corresponding ASC dimension for the corresponding drug. We did not perform a meta-analysis in this case, so instead we report the data from the single study.*

| | LSD | | | Psilocybin | | |
|---|---|---|---|---|---|---|
| | Low Dose (0.05-0.074 mg) | Medium Dose (0.075-0.109 mg) | High Dose (≥0.110 mg) | Low Dose (8-14 mg) | Medium Dose (15-21 mg) | High Dose (≥22 mg) |
| EU | 16.79 ± 4.26 | 42.55 ± 4.42 | 48.44 ± 3.47 | 28.34 ± 6.89 | 27.92 ± 3.43 | 44.93 ± 4.56 |
| SE | 10.18 ± 3.07 | 22.39 ± 2.89 | 27.6 ± 3.87 | 6.49 ± 4.78 | 16.50 ± 3.14 | 39.42 ± 15.99 |
| BS | 33.36 ± 9.82 | 42.26 ± 4.45 | 48.3 ± 7.13 | 34.24 ± 10.58 | 41.62 ± 4.38 | 49.74 ± 8.82 |
| I | 25.43 ± 14.7 | 34.72 ± 3.04 | 40.12 ± 3.89 | 30.99 ± 16.89 | 24.91 ± 3.25 | 33.98 ± 7.3 |
| D | 7.27 ± 2.73 | 39.24 ± 5.14 | 42.53 ± 3.44 | 22.87 ± 4.92 | 29.81 ± 5.34 | 41.88 ± 2.25 |
| ICC | 10.13 ± 3.15 | 28.43 ± 2.78 | 36.91 ± 3.43 | 11.67 ± 6.73 | 19.43 ± 2.04 | 21.52 ± 5.6 |
| A | 2.84 ± 4.93 | 8.76 ± 1.78 | 18.99 ± 3.50 | 1.56 ± 6.64 | 8.74 ± 1.76 | 16.41 ± 2.92 |
| CI | 37.25 ± 7.12 | 58.84 ± 3.42 | 69.64 ± 4.06 | 40.25 ± 8.33 | 53.42 ± 4.18 | 64.54 ± 10.36 |
| EI | 41.28 ± 14.1 | 53.67 ± 3.25 | 61.59 ± 4.04 | 38.50 ± 14.56 | 42.71 ± 4.25 | 54.06 ± 3.66 |
| AVS | 22.85 ± 8.24 | 59.42 ± 3.61 | 75.22 ± 6.43 | 36.99 ± 11.57 | 42.23 ± 5.17 | 49.42 ± 19.81 |
| CMP | 18.13 ± 2.95 | 34.92 ± 2.88 | 46.19 ± 6.83 | 30.83 ± 8.39 | 27.44 ± 2.89 | 26.36 ± 8.02 |

***Table S2. Numerical data for 11D-ASC meta-analysis.*** *Data given as percentage of maximum score.*

| Reference | Con | SP | CI | DII | MD | MO | SRR | Overall RoB |
|---|---|---|---|---|---|---|---|---|
| Becker et al. (2023) | M | L | L | L | L | M | L | M |
| Bernasconi et al. (2014) | M | L | L | L | L | S | L | S |
| Carbonaro et al. (2018) | M | L | L | L | L | L | L | M |
| Carter, Burr, et al. (2005) | M | L | L | L | L | M | L | M |
| Carter, Pettigrew, et al. (2005) | M | L | L | L | L | M | L | M |
| Carter et al. (2007) | M | L | L | L | L | M | L | M |
| Daumann et al. (2008) | S | L | L | L | L | M | L | S |
| Duerler et al. (2020) | S | L | L | L | L | M | L | S |
| Family et al. (2022) | M | L | L | L | L | M | L | M |
| Gouzoulis-Mayfrank et al. (2005) | M | L | L | L | L | M | L | M |
| Hasler et al. (2004) | L | L | L | L | L | M | L | M |
| Holze et al. (2020) | M | L | L | L | L | M | L | M |
| Holze et al. (2021) | M | L | L | L | L | M | L | M |
| Holze et al. (2022) | M | L | L | L | L | M | L | M |
| Kometer et al. (2012) | M | L | L | L | L | M | L | M |
| Kraehenmann, Preller, et al. (2015) | M | L | L | L | L | S | L | S |
| Kraehenmann et al. (2017) | M | L | L | L | L | M | L | M |
| Lewis et al. (2017) | M | L | L | L | L | S | L | S |

| | | | | | | | | |
|---|---|---|---|---|---|---|---|---|
| Ley et al. (2023) | M | L | L | L | L | L | L | M |
| Madsen et al. (2021) | M | L | L | L | L | M | L | M |
| Mallaroni et al. (2023) | M | L | L | L | L | M | L | M |
| Mason et al. (2020) | M | L | L | L | L | M | L | M |
| McAlpine et al. (2023) | S | L | L | L | L | S | L | S |
| Müller, Lenz, Dolder, Lang, et al., 2017 | M | L | L | L | L | S | L | S |
| Ort et al. (2023) | S | L | L | L | L | L | L | S |
| Pallavicini et al. (2021) | M | L | L | L | L | S | L | S |
| Pokorny et al. (2016) | S | L | L | L | L | M | L | S |
| Pokorny et al. (2017) | S | L | L | L | L | S | L | S |
| Pokorny et al. (2020) | S | L | L | L | L | M | L | S |
| Preller et al. (2016) | S | L | L | L | L | S | L | S |
| Preller et al. (2017) | M | L | L | L | L | M | L | M |
| Preller et al. (2020) | M | L | L | L | L | S | L | S |
| Quednow et al. (2012) | M | L | L | L | L | M | L | M |
| Schmid et al. (2015) | M | L | L | L | L | S | L | S |
| Schmidt et al. (2012) | S | L | L | L | L | M | L | S |
| Schmidt et al. (2018) | M | L | L | L | L | S | L | S |
| Smigielski et al. (2020) | S | L | L | L | L | S | L | S |
| Straumann et al. (2023) | M | L | L | L | L | M | L | M |

| | | | | | | | |
|---|---|---|---|---|---|---|---|
| Timmermann et al. (2023) | M | L | L | L | L | S | L | S |
| Umbricht et al. (2002) | S | L | L | L | L | S | L | S |
| Vogt et al. (2023) | M | L | L | L | L | M | L | M |
| Vollenweider et al. (2007) | M | L | L | L | L | M | L | M |
| Wießner et al. (2023) | M | L | L | L | L | S | L | S |
| Wittmann et al. (2007) | M | L | L | L | L | M | L | M |

***Table S3. Risk of bias for individual studies in the phenomenology literature.*** *Using the ROBINS-I tool, we analysed risk of bias in seven domains: bias due to confounding (Con), bias in selection of participants into the study (SP), bias in classification of interventions (CI), bias due to deviations from intended interventions (DII), bias due to missing data (MD), bias in measurement of outcomes (MO), and bias in selection of the reported result (SRR). The overall risk of bias (RoB) is given in the rightmost column. L, M, and S refer to low, moderate, and serious risk of bias. Risk of bias was deemed low for all studies in the SP, CI, DII, MD, and SRR domains. Most studies received either an M or an S rating for the Con domain due either to confounding from the participants' levels of prior psychedelic use or failure to measure this characteristic, respectively. Many studies were rated M in the MO domain because it was possible for participants to guess the intervention (i.e., drug and, if applicable, dose level) that they had received. Some studies were rated S because they used a within-subjects, cross-over design in which the psychedelic and the placebo were the only two drugs that were administered. Participants who received a psychedelic in the first session were likely to know with certainty that they would get a placebo in the second session; thus, they may respond that they experienced zero effects along all dimensions of the ASC scale, whereas participants who received placebo in the first session may give low but non-zero responses for some dimensions. Overall RoB was either M or S for all studies.*

| Study in which primary dataset was first reported | Secondary analyses | Analysis included in the FC meta-analysis |
|---|---|---|
| Carhart-Harris, Erritzoe, et al. (2012) | Lebedev et al. (2015) | Roseman et al. (2014) |
| | Roseman et al. (2014) | |
| Carhart-Harris et al. (2016) | Carhart-Harris et al. (2016): seed-to-voxel analysis | Dai et al. (2023) |
| | Carhart-Harris et al. (2016): ICA-based analysis | |
| | Dai et al. (2023) | |
| | Luppi et al. (2021) | |
| | Tagliazucchi et al. (2014) | |
| Holze et al. (2020)* | Avram et al. (2022) | Bedford et al. (2023)** |
| | Bedford et al. (2023)** | |
| Mason et al. (2020) | Mason et al. (2021) | Mason et al. (2020)*** |
| Müller, Lenz, Dolder, Lang, et al. (2017) | Bedford et al. (2023)** | Bedford et al. (2023)** |
| | Müller, Dolder, et al. (2018) | |
| | Müller, Lenz, Dolder, Lang, et al. (2017): thalamic seed-to-seed analysis | |
| | Müller, Lenz, Dolder, Lang, et al. (2017): seed-to-voxel analysis | |
| Timmermann et al. (2023) | Timmermann et al. (2023): seed-to-seed analysis | Timmermann et al. (2023): seed-to-seed analysis |
| | Timmermann et al. (2023): ICA-based analysis | |

***Table S4. Studies that were included in the functional connectivity meta-analysis, when there was more than one reported analysis of the corresponding dataset.*** *The existence of multiple studies on the same primary dataset biases the results of the FC meta-analysis. We only included one analysis of each unique dataset in the meta-analysis. This analysis was the one that spanned the largest portion of the brain and used the most fine-grained parcellation, as long as it did not apply the network-based statistic, which underestimates the number of significant connections relative to other techniques. *Holze et al. (2020) was not identified in the neuroimaging literature search because the publication did not report any fMRI findings, though fMRI data was collected as part of the study. **Bedford et al. (2023) assessed data from both Avram et al. (2022) and Müller, Lenz, Dolder, Lang et al. (2017). ***The ICA-based analysis reported in Mason et al. (2021) is almost exactly the same as the one reported in Mason et al. (2020), except it additionally includes the salience network in its analysis.*

| Functional Connectivity Study and Method | Inputs |
|---|---|
| de Araujo et al. (2011): seed-to-seed analysis | 1. Spatial maps of specific seeds from the Brodmann areas |
| Barrett, Krimmel, et al. (2020): seed-to-seed analysis | 1. Spatial maps of left and right claustrum seeds, obtained from the authors<br>2. Networks of 10mm-radius spheres around coordinates of network seeds in the Power atlas |
| Bedford et al. (2023): seed-to-seed analysis | 1. Spatial maps of seeds extracted from the Harvard-Oxford atlas |
| Dai et al. (2023): seed-to-seed analysis | 2. Spatial maps of seeds extracted from the HCP-ICA atlas |
| Gaddis et al. (2022): ICA | 1. Cortical: spatial maps of template group ICs from Smith et al. (2009)*; thalamic: template group ICs obtained from author |
| Grimm et al. (2018): seed-to-voxel analysis | 1. Spatial maps of the right amygdala seed, extracted from the AAL atlas<br>2. 10mm-radius spheres around the coordinates of voxels that had peak FC with the seed |
| Kaelen et al. (2016): seed-to-voxel analysis | 1. Spatial map of the parahippocampus seed, extracted from the Harvard-Oxford atlas<br>2. Spatial map of voxels that had peak FC with the seed, obtained from the authors |
| Madsen et al. (2021): seed-to-seed analysis | 1. Networks of 10mm-radius spheres around coordinates of network seeds in the Raichle atlas |
| Mason et al. (2020/21): ICA | 1. Spatial maps of template group ICs from Smith et al. (2009)* |
| Palhano-Fontes et al. (2015): seed-to-voxel analysis (not DMN-TPN orthogonality test) | 1. 10mm-radius spheres around coordinates of PCC and mPFC seeds<br>2. 10mm-radius spheres around the coordinates of voxels that had peak FC with the seed |
| Roseman et al. (2014): ICA | 1. Spatial maps of template group ICs from Smith et al. (2009)** |
| Timmermann et al. (2023): seed-to-seed analysis | 1. Spatial maps of seeds extracted from the Schaefer-100 atlas and subcortical seeds in the AAL atlas |

***Table S5. Summary of inputs into the functional connectivity meta-analysis.*** *Our scoring algorithm compared data from three kinds of studies: seed-to-seed, seed-to-voxel, and ICA-based analyses. For seed-to-seed studies, we obtained spatial maps of the seeds, either from the relevant parcellation or from the authors themselves. For seed-to-voxel studies, we tried to obtain spatial maps of the clusters of voxels that were significantly connected to the seeds; when this was not possible, we constructed spheres around the MNI coordinates of the clusters, where the radius of the sphere varied depending on the size of the cluster. For ICA-based studies, we used spatial maps of group independent components (ICs). In some ICA-based studies, group ICs were either *cross-correlated with template ICs from Smith et al. (2009); others **used those template ICs for dual regression/back-projection into subject-specific spatial IC maps. Thus, to run the meta-analysis on the ICA-based analyses, we used the template ICs from Smith et al. (2009).*

| Reference | Con | SP | CI | DII | MD | MO | SRR | Overall RoB |
|---|---|---|---|---|---|---|---|---|
| Barrett, Krimmel, et al. (2020) | M | L | L | L | L | M | L | M |
| Bedford et al. (2023) | M | L | L | L | L | L | L | M |
| Dai et al. (2023) | M | L | L | L | L | M | L | M |
| de Araujo et al. (2011) | S | L | L | L | L | M | L | S |
| Gaddis et al. (2022) | M | L | L | L | L | M | L | M |
| Grimm et al. (2018) | S | L | L | L | L | L | L | S |
| Kaelen et al. (2016) | M | L | L | L | L | M | L | M |
| Madsen et al. (2021) | M | L | L | L | L | M | L | M |
| Mason et al. (2020) | M | L | L | L | M | L | L | M |
| Palhano-Fontes et al. (2015) | M | L | L | L | L | M | L | M |
| Roseman et al. (2014) | M | L | L | L | L | M | L | M |
| Timmermann et al. (2023) | L | L | L | L | L | M | L | M |

***Table S6. Risk of bias for individual studies in the neuroimaging literature.*** *Using the ROBINS-I tool, we analysed risk of bias in seven domains: bias due to confounding (Con), bias in selection of participants into the study (SP), bias in classification of interventions (CI), bias due to deviations from intended interventions (DII), bias due to missing data (MD), bias in measurement of outcomes (MO), and bias in selection of the reported result (SRR). The overall risk of bias (RoB) is given in the rightmost column. L, M, and S refer to low, moderate, and serious risk of bias. Risk of bias was deemed low for all studies in the SP, CI, DII, and SRR domains. One study used a between-subjects design and excluded more subjects in the psilocybin group than the placebo group due to head motion, so it was assigned a moderate risk of bias in the MD domain (Mason et al., 2020). Psychedelics are known to be vasoconstrictors (Dyer & Gant, 1973), which may influence the fMRI signal (Özbay et al., 2019). None of the studies explicitly controlled for vasoconstriction, but it is also unclear how to feasibly measure neural vasoconstriction in a study with humans, so most studies were assigned a moderate risk of bias in the Con domain. Timmermann et al. (2023) received an L rating in this domain because it measured EEG and fMRI simultaneously, and EEG signal is less susceptible to vascular changes than fMRI. Studies that did not report the results of head motion analyses, which are known to strongly confound fMRI measurements (Makowski et al., 2019), were given an S rating in the Con domain. Studies that were not double-blind were deemed to have moderate risk of bias in the MO domain. In this case, the outcome assessors were aware of whether participants received a psychedelic or placebo, but this likely did not have a significant effect on the measurement outcomes. Overall risk of bias was moderate for 10/12 studies and serious for 2/12 studies.*

| | 5-$HT_{2A}$/5-$HT_{1A}$ | 5-$HT_{2C}$/5-$HT_{1A}$ | $D_2$/5-$HT_{1A}$ |
|---|---|---|---|
| DMT | 1.03 ± 0.44 | 1.00 ± 0.39 | 0.03 ± 0.003 |
| LSD | 1.28 ± 0.41 | 0.41 ± 0.21 | 0.07 ± 0.01 |
| Psilocin | 2.07 ± 0.68 | 1.02 ± 0.37 | 0.03 ± 0.003 |

***Table S7. Numerical results for meta-analysis of selectivity relative to 5-$HT_{1A}$.*** *Data given as the ratio between the $K_i$ for the receptor of interest and the $K_i$ for 5-$HT_{1A}$.*

| | 5-$HT_{1A}$/5-$HT_{2A}$ | 5-$HT_{2C}$/5-$HT_{2A}$ | $D_2$/5-$HT_{2A}$ |
|---|---|---|---|
| DMT | 1.28 ± 0.27 | 1.40 ± 0.14 | 0.08 ± 0.003 |
| LSD | 1.06 ± 0.49 | 0.30 ± 0.27 | 0.02 ± 0.02 |
| Psilocin | 0.46 ± 0.47 | 0.62 ± 0.23 | 0.01 ± 0.0001 |

***Table S8. Numerical results for meta-analysis of selectivity relative to 5-$HT_{2A}$.*** *Data given as the ratio between the $K_i$ for the receptor of interest and the $K_i$ for 5-$HT_{2A}$.*

| | β-Arrestin2 Recruitment | Calcium Mobilisation | Inositol Phosphate Formation |
|---|---|---|---|
| DMT | 1.32 ± 0.44 | -1.89 ± 0.83 | -1.40 ± 0.22 |
| LSD | 2.59 ± 0.31 | -0.73 ± 0.49 | 1.62 ± 0.16 |
| Psilocin | 2.10 ± 0.43 | -1.02 ± 0.6 | -0.36 ± 0.31 |

***Table S9. Numerical results for functional activity meta-analysis.*** *All assays included in this meta-analysis were performed on the 5-$HT_{2A}$ receptor. Data given as relative activity, which is* $\Delta\log(E_{max}/EC_{50})$. Δ *refers to the difference between the relative activity of the drug of interest and that of a reference ligand. The reference ligand was serotonin for the calcium mobilisation and inositol phosphate formation assays and mescaline for the β-arrestin2 recruitment assays.*
